# A large cross-ancestry meta-analysis of genome-wide association studies identifies 69 novel risk loci for primary open-angle glaucoma and includes a genetic link with Alzheimer’s disease

**DOI:** 10.1101/2020.01.30.927822

**Authors:** Puya Gharahkhani, Eric Jorgenson, Pirro Hysi, Anthony P. Khawaja, Sarah Pendergrass, Xikun Han, Jue Sheng Ong, Alex W. Hewitt, Ayellet Segre, Robert P. Igo, Helene Choquet, Ayub Qassim, Navya S Josyula, Jessica N. Cooke Bailey, Pieter Bonnemaijer, Adriana Iglesias, Owen M. Siggs, Terri Young, Veronique Vitart, Alberta A.H.J. Thiadens, Juha Karjalainen, Steffen Uebe, Ronald B. Melles, K. Saidas Nair, Robert Luben, Mark Simcoe, Nishani Amersinghe, Angela J. Cree, Rene Hohn, Alicia Poplawski, Li Jia Chen (CUHK), Ching-Yu Cheng, Eranga Nishanthie Vithana, NEIGHBORHOOD consortium, ANZRAG consortium, Biobank Japan project, FinnGen study, UK Biobank Eye and Vision Consortium, GIGA study group, 23andMe Research Team, Gen Tamiya, Yukihiro Shiga, Masayuki Yamamoto, Toru Nakazawa, John Rouhana, Hannah Currant, Ewan Birney, Xin Wang, Adam Auton, Adeyinka Ashaye, Olusola Olawoye, Susan E. Williams, Stephen Akafo, Michele Ramsay, Kazuki Hashimoto, Yoichito Kamatani, Masato Akiama, Yukihide Momozawa, Paul J. Foster, Peng T. Khaw, James E. Morgan, Nicholas G. Strouthidis, Peter Kraft, Jae Hee Kang, Calvin Chi Pui Pang (CUHK), Francesca Pasutto, Paul Mitchell, Andrew J. Lotery, Aarno Palotie, Cornelia van Duijn, Jonathan Haines, Chris Hammond, Louis R. Pasquale, Caroline C.W. Klaver, Michael Hauser, Chiea Chuen Khor, David A. Mackey, Michiaki Kubo, Tin Aung, Jamie Craig, Stuart MacGregor, Janey Wiggs

## Abstract

We conducted a large multi-ethnic meta-analysis of genome-wide association studies for primary open-angle glaucoma (POAG) on a total of 34,179 cases vs 349,321 controls, and identified 127 independent risk loci, almost doubling the number of known loci for POAG. The majority of loci have broadly consistent effect across European, Asian and African ancestries. We identify a link, both genome-wide and at specific loci, between POAG and Alzheimer’s disease. Gene expression data and bioinformatic functional analyses provide further support for the functional relevance of the POAG risk genes. Several drug compounds target these risk genes and may be potential candidates for developing novel POAG treatments.

---

Primary open-angle glaucoma (POAG) is the leading cause of irreversible blindness worldwide^1, 2^. The disease is characterized by progressive optic nerve degeneration that is usually accompanied by elevated intraocular pressure (IOP). Neuroprotective therapies are not available and current treatments are limited to lowering IOP which can slow disease progression at early disease stages. However over 50% of glaucoma is not diagnosed until irreversible optic nerve damage has occurred ^2, 3^.

POAG is highly heritable^4, 5^, and previous genome-wide association studies (GWAS) have identified important loci associated with POAG risk and IOP^6–14^. Despite this success, the POAG genetic landscape remains incomplete and identification of novel risk loci is required to further define contributing disease mechanisms that could be targets of novel preventative therapies.

The majority of known risk loci for POAG have been identified through GWAS in participants of European descent, followed by replication in other ethnic populations. However, previous observational studies have shown that individuals of African ancestry, followed by Latinos and Asians, have higher POAG disease burden compared to those of European ancestry^3, 15–17^, suggesting important differences in genetic risk and highlighting the need to compare the genetic architecture of these ethnic groups.

In this study, we report the results of GWAS studies on a large collection of POAG cases and controls of European, Asian, and African descent. We performed the largest multi-ethnic meta-analysis of POAG GWAS to date, identifying 127 risk loci (69 not previously reported at genome-wide significant levels for POAG). The majority of loci have broadly consistent effects on glaucoma risk, regardless of ancestry, with our large scale approaches uncovering a genetic link between POAG and Alzheimer’s disease (AD).

## Results

### Discovery of novel POAG risk loci in Europeans

We performed a four-stage meta-analysis (Fig 1). In the first stage, we conducted a fixed-effect meta-analysis of 16,677 POAG cases vs. 199,580 controls of European descent. The participating studies are detailed in Supplementary Table 1. We identified 66 independent genome-wide significant (P<5e-08) SNPs (Supplementary Table 2), of which 25 were novel (i.e. uncorrelated with previously reported SNPs). There was no evidence indicating that the results were affected by population structure (LD score regression^18^ intercept 1.03, se=0.01, Supplementary Fig 1A).

**Figure 1.**
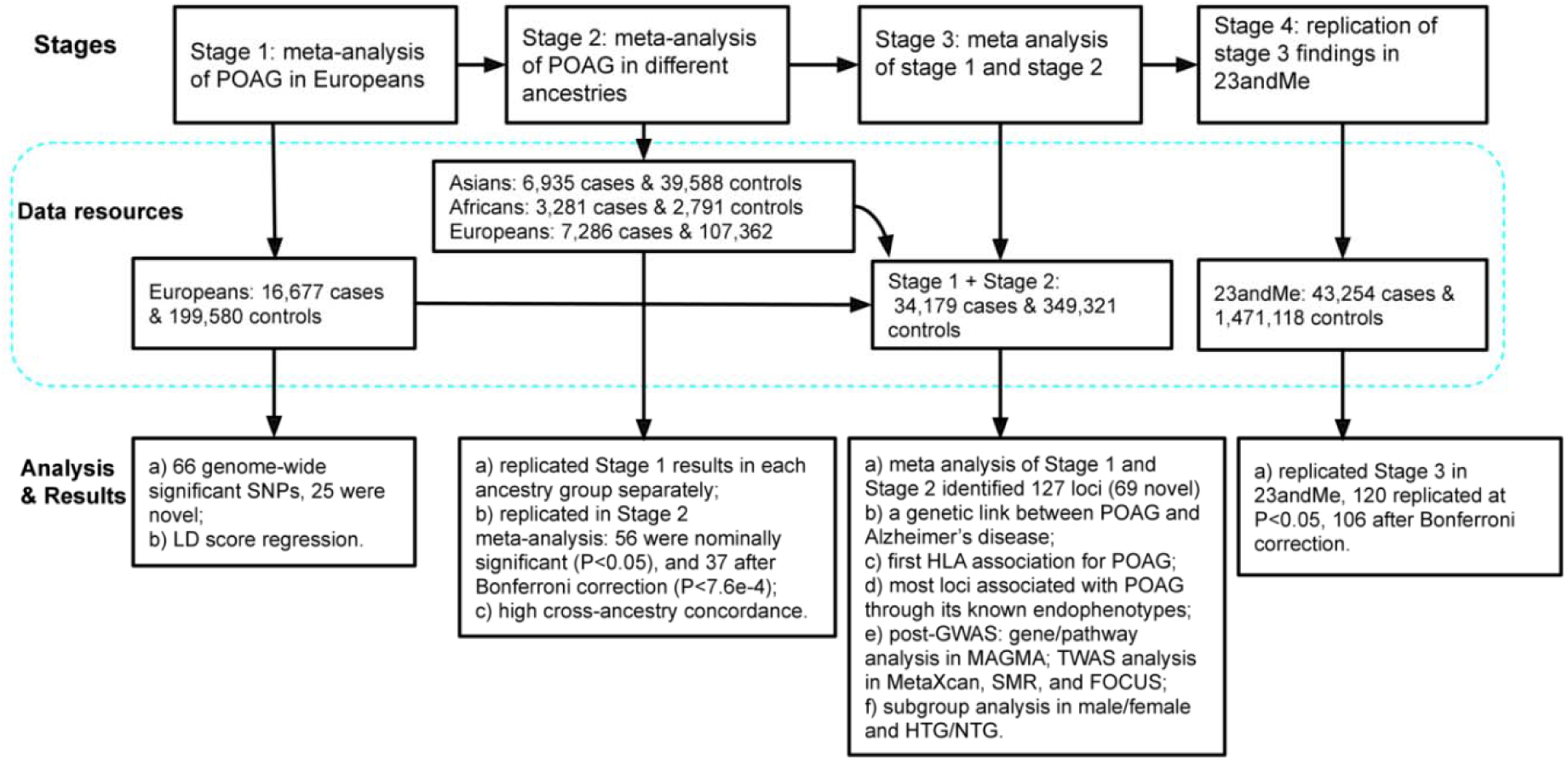
Study design. This Figure summarises the four stages of this study as well as the data resources and main analyses/results for each stage.

### Consistent genetic effect across ancestries

In the second stage, we replicated the stage 1 findings in 3 independent datasets comprising 7,286 self-reported cases and 107,362 controls of European descent from the UK Biobank study (UKBB)^19^, 6,935 POAG cases and 39,588 controls of Asian descent, and 3,281 POAG cases and 2,791 controls of African ancestry (Supplementary Table 1). We replicated the results in each ancestry group separately (Supplementary Table 2, Supplementary Fig 2), followed by combining the three replication datasets in a fixed-effect meta-analysis (17,502 cases and 149,741 controls) to maximize statistical power. Of the 66 loci significant in the discovery GWAS, 56 were nominally significant (P<0.05), and 37 after Bonferroni correction (P<7.6e-04) in the meta-analyzed replication cohort. The effect sizes had a Pearson correlation coefficient (r)=0.93 (Fig 2).

**Figure 2.**
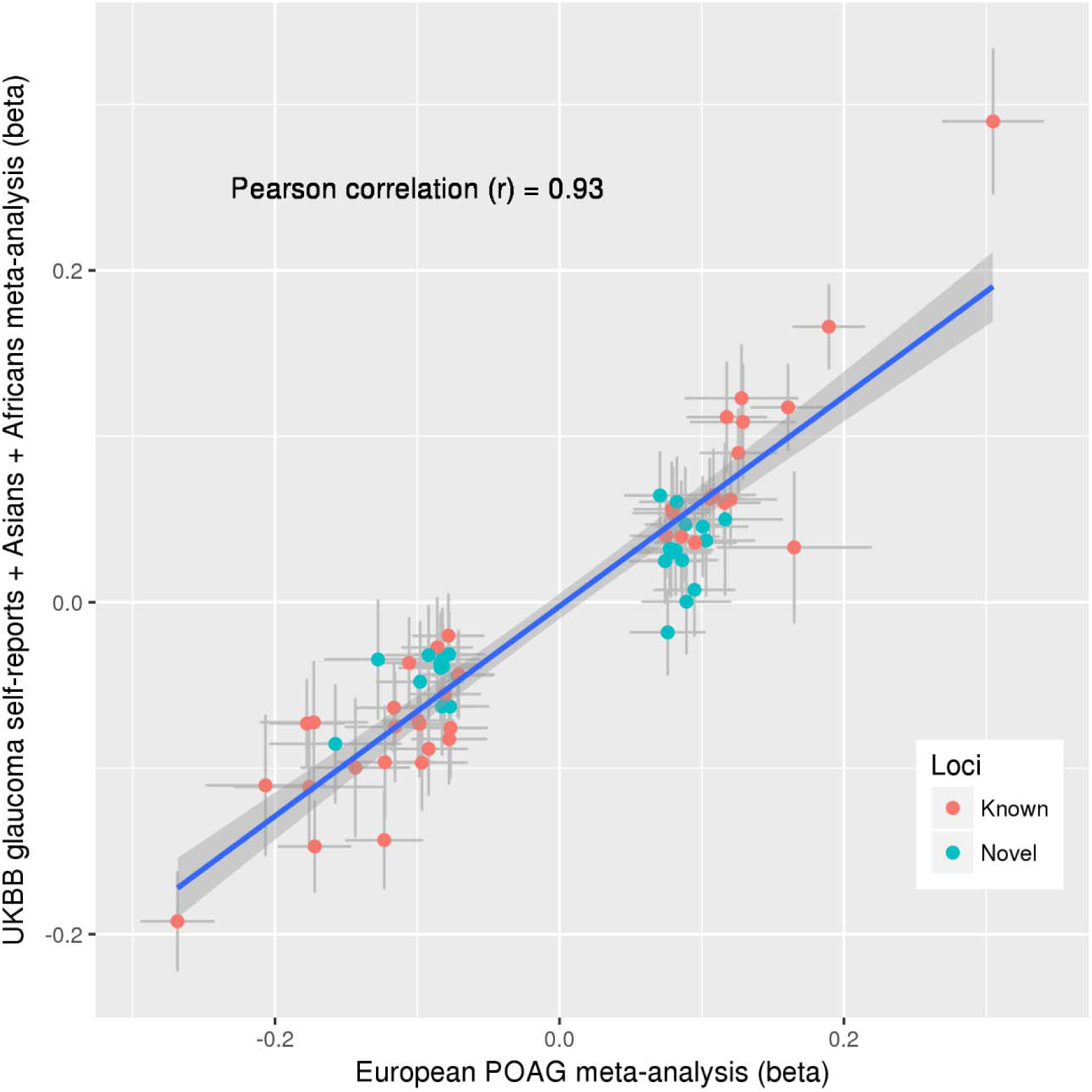
Correlation of SNP effect estimates between the European POAG meta-analysis and the replication dataset. The X-axis shows effect estimates in log(OR) scale for the independent genome-wide significant loci obtained from the meta-analysis of POAG in Europeans (16,677 POAG cases vs 199,580 controls). The Y-axis shows the effect estimates in log(OR) scale for the same SNPs obtained from meta-analysis of the following three GWAS data: glaucoma self-reports in UKBB, POAG in Asians, and POAG in Africans (the overall sample size of 17,502 cases and 149,741 controls). Red dots are the previously-identified risk loci and blue dots are the novel risk loci identified in this study. Horizontal grey bars on each dot represent the 95% confidence intervals (CIs) for the effect estimates in Europeans, and vertical grey bars shows the 95% CIs in the replication dataset. The blue line is the linear regression line best fitting the data. The Pearson correlation between the effect estimates is 0.93 (95% CIs 0.89-0.96).

There was high cross-ancestry concordance both for genome-wide significant loci and across the genome. For the genome-wide significant SNPs, the European SNP effects were correlated (Pearson correlation coefficient (r)=0.78 [95% confidence intervals (CIs) 0.66-0.86] and r=0.68 [95% CIs 0.52-0.79]) with Asian and African ancestries, respectively (Supplementary Fig 2B and 2C). The genome-wide genetic correlation estimated using the approach implemented in Popcorn^20^ was even higher: r=0.85 (95% CIs 0.70-1.00) for European-Asian and r=0.75 (95% CIs, −0.93 to 2.43) for European-African. Although the concordance amongst the top SNPs was clear for the European-African comparison, larger sample sizes will be required to narrow the CIs on the European-African genome-wide correlation estimate.

### Significant risk loci in the meta-analysis of POAG in Asians and Africans

Ten loci were genome-wide significant for POAG in the meta-analysis of Asian studies (Supplementary Table 3), all of which are known POAG loci, and all associated in the European meta-analysis or UKBB self-reports at Bonferroni-corrected P<0.05/10. While only one of these loci had a P<0.05 in Africans, 8 had consistent direction of effect. For the African meta-analysis, one locus (rs16944405 within *IQGAP1*) reached the genome-wide significance level (P=3e-08). This locus has not been previously reported for POAG. In this study, it was not associated with POAG in Europeans but was nominally associated in Asians (Supplementary Table 3). The LD score regression intercept was 0.99 (se=0.009) for Asians and 0.95 (se=0.006) for Africans, suggesting that the results were not influenced by population structure.

### Discovery of novel POAG risk loci in a multi-ancestry meta-analysis

In the third stage, given the large genetic correlation between ancestries, we performed a fixed-effect meta-analysis of GWAS results from stage 1 and 2 (34,179 cases vs 349,321 controls). We identified 127 independent genome-wide significant loci, located at least >1Mb apart (Fig 3, Supplementary Fig 3). Of these, 69 loci were not previously associated with POAG at genome-wide significant levels (Supplementary Table 4). Of note, four of the novel risk loci (*MXRA5-PRKX*, *GPM6B*, *NDP-EFHC2,* and *TDGF1P3-CHRDL1*) are on the X chromosome, representing the first POAG risk loci on a sex chromosome.

**Figure 3.**
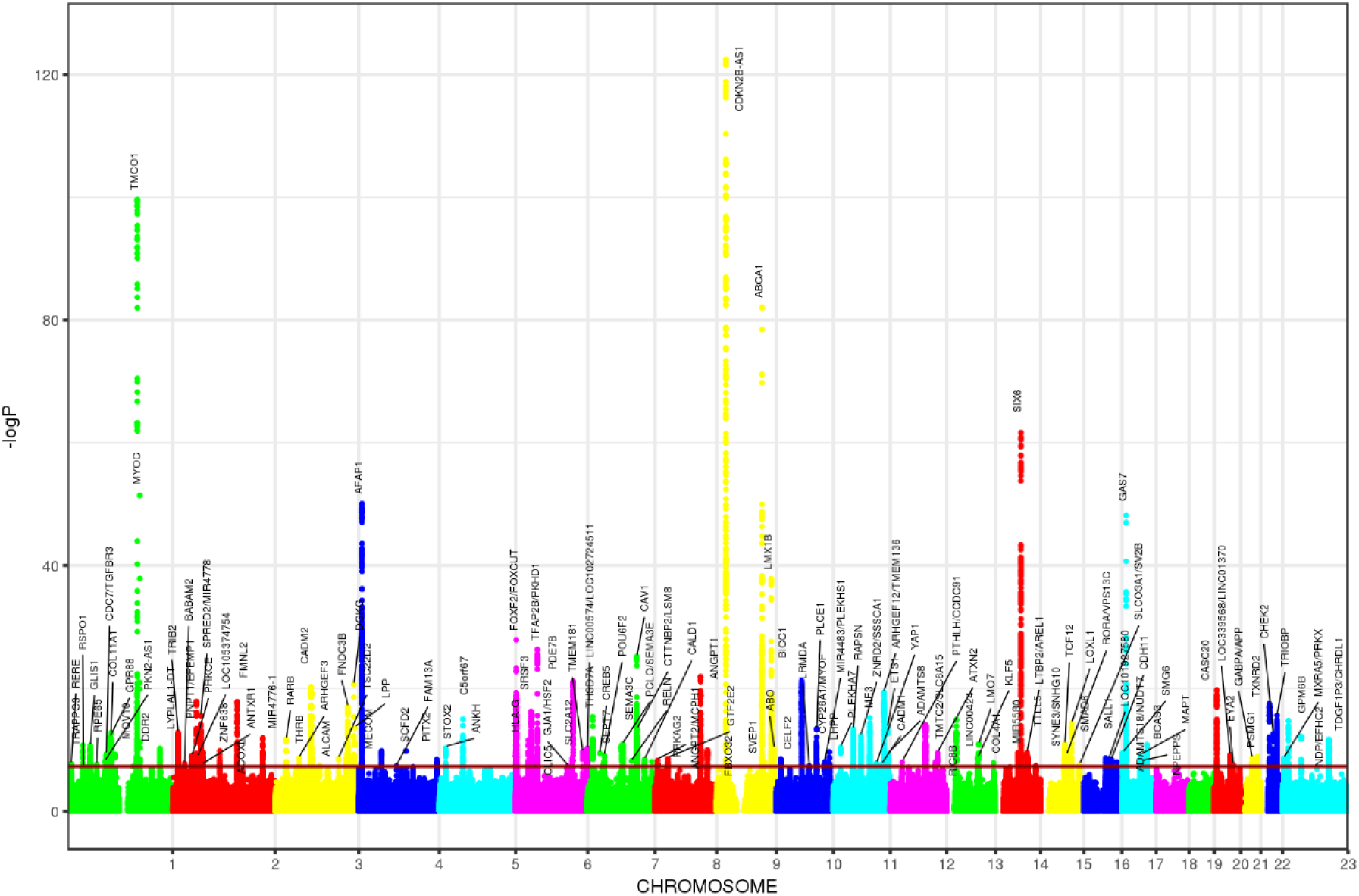
Manhattan plots for the cross-ancestry meta-analysis. Each dot represents a SNP, the X-axis shows the chromosomes where each SNP is located, and the Y-axis shows -log10 P-value of the association of each SNP with POAG in the cross-ancestry meta-analysis. The red horizontal line shows the genome-wide significant threshold (P-value=5e-8; -log10 P-value=7.30). The nearest gene to the most significant SNP in each locus has been labeled.

### The POAG risk loci were strongly replicated in 23andMe

In the fourth stage, we validated the association of the genome-wide significant SNPs from stage 3 in a dataset comprising 43,254 participants with self-reported POAG (defined as those who reported having glaucoma excluding angle-closure glaucoma or other types of glaucoma) and 1,471,118 controls from 23andMe, Inc. Of the 127 loci, the association results for 125 SNPs were available in 23andMe, 120 of which (96%) were replicated at P<0.05, and 106 (85%) after Bonferroni correction for 125 independent tests (Supplementary Table 4). The correlation of the effect size was r=0.98 (95% CIs 0.977-0.989).

### A genetic link between POAG and Alzheimer’s disease

Three POAG risk loci we identified (*MAPT, CADM2,* and *APP)* have also been implicated in AD and dementia^21–23^. *MAPT -* a lead SNP for AD (rs2732703) – was highly correlated with the POAG lead SNP (rs242559; LD r2=0.89). The *MAPT* rs2732703 had showed a genome-wide significant association with AD (P=6e-09) in 4-subjects^22^ although in our meta-analysis, it did not reach the genome-wide significance threshold for POAG (P=1.2e-06). Both SNPs are strong eQTLs for *MAPT* and several other genes in this region in multiple GTEx tissues^24^. To further investigate the genetic link between POAG and AD, we used LD-Score^25^ to estimate their genome-wide genetic correlation using GWAS summary data from our POAG meta-analysis in Europeans (excluding self-report) and the publicly available GWAS data^26^ for 71,880 AD cases and 383,378 controls of European descent. The genome-wide genetic correlation was 0.14 (95% CI: 0.003-0.28; P=0.049) between glaucoma and AD, suggesting that multiple loci across the genome may contribute to the risk of both diseases. Given the AD link, we performed a post hoc lookup of rs4420638[A] which tags the well-known *APOE* □4 AD risk haplotype; the SNP showed nominal association with POAG in the all-ancestry meta-analysis (P=0.00153).

### First HLA association for POAG

We also, for the first time, identified an association of a human leukocyte antigen (HLA gene (HLA-G/HLA-H) with POAG. The HLA system is a gene complex encoding the major histocompatibility complex proteins in humans. These cell-surface proteins are responsible for the regulation of the human immune system. The most significant SNP in this region (rs407238) has been associated at the genome-wide significance level for other traits such as Celiac disease, intestinal malabsorption, disorders of iron metabolism, multiple blood traits, hyperthyroidism, multiple sclerosis, hip circumference, and weight (https://genetics.opentargets.org/variant/6_29839124_C_G). The mechanism of action of the lead SNP appears to be via IOP (UKBB IOP GWAS P=8.8e-06).

### Most of the risk loci associated with POAG act via its known endophenotypes

All POAG-risk variants identified to date have also been associated with either IOP or with altered optic nerve morphology (typically, measured as vertical cup-to-disc ratio (VCDR)) or both. To investigate the association of the POAG loci with IOP and VCDR, we used GWAS of IOP (N=133,492)^11^ and VCDR (N=23,899)^27^. Fig 4A shows that the majority of loci (89 of 123; four were unavailable for IOP) act primarily via IOP (red and green dots on Fig 4A - a putative “IOP route to glaucoma”) but that there were a set of 34 loci with unclear effect on IOP (purple and blue dots on Fig 4A, full data in Supplementary Table 5). Plotting the POAG effect sizes against the VCDR effect sizes for 32 of these 34 SNPs (2 SNPs unavailable for VCDR), we can see that the majority of the POAG loci not acting via IOP have a clear effect on VCDR (purple dots on Fig 4B - a putative “VCDR route to glaucoma”). There were a small number (13) of POAG loci where there was no clear effect on IOP or VCDR, although in three cases the peak SNP was not associated with glaucoma in 23andMe and in the remaining cases, it was difficult to comprehensively rule out an effect on glaucoma via a small change in IOP or VCDR.

**Figure 4.**
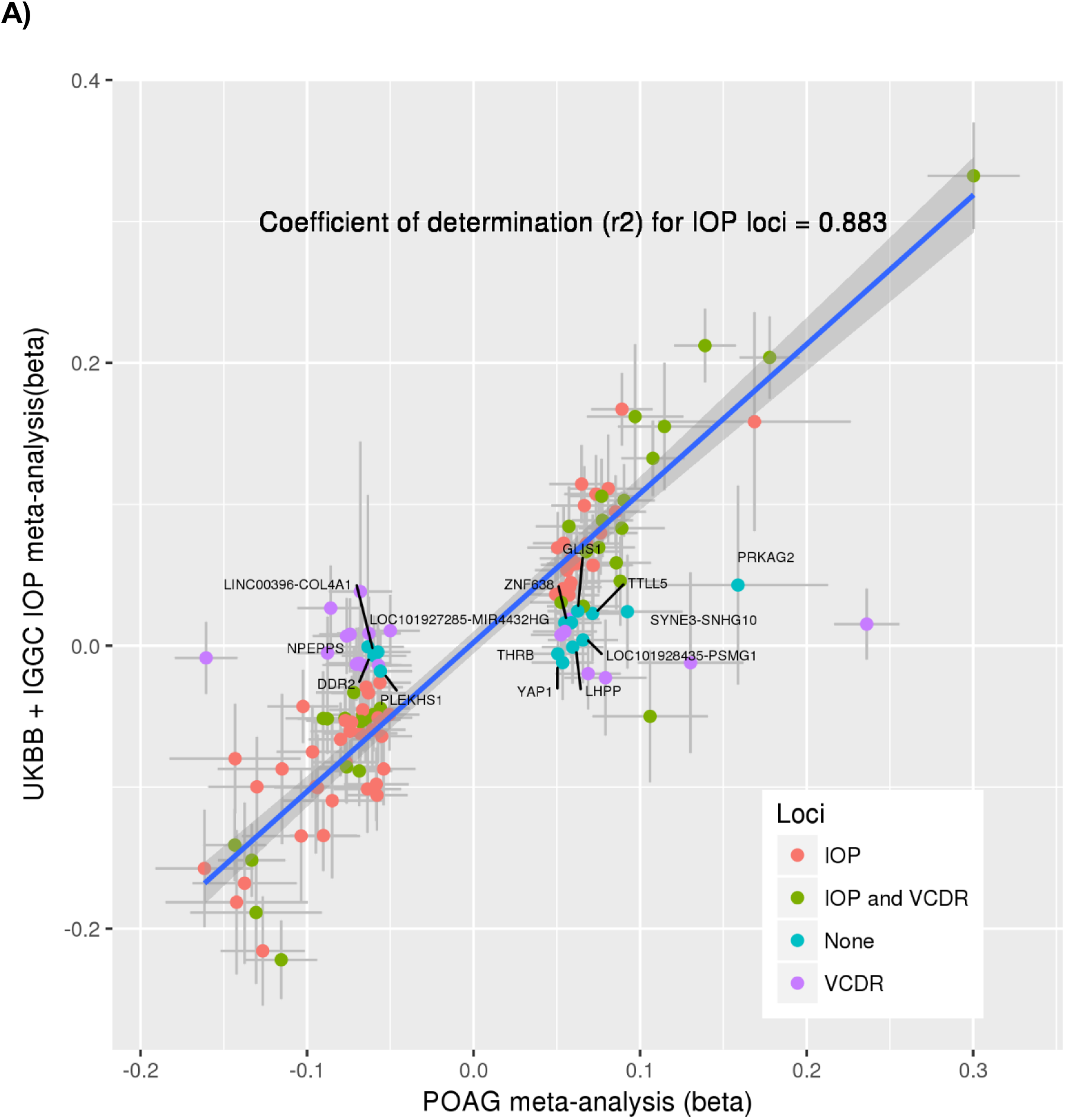
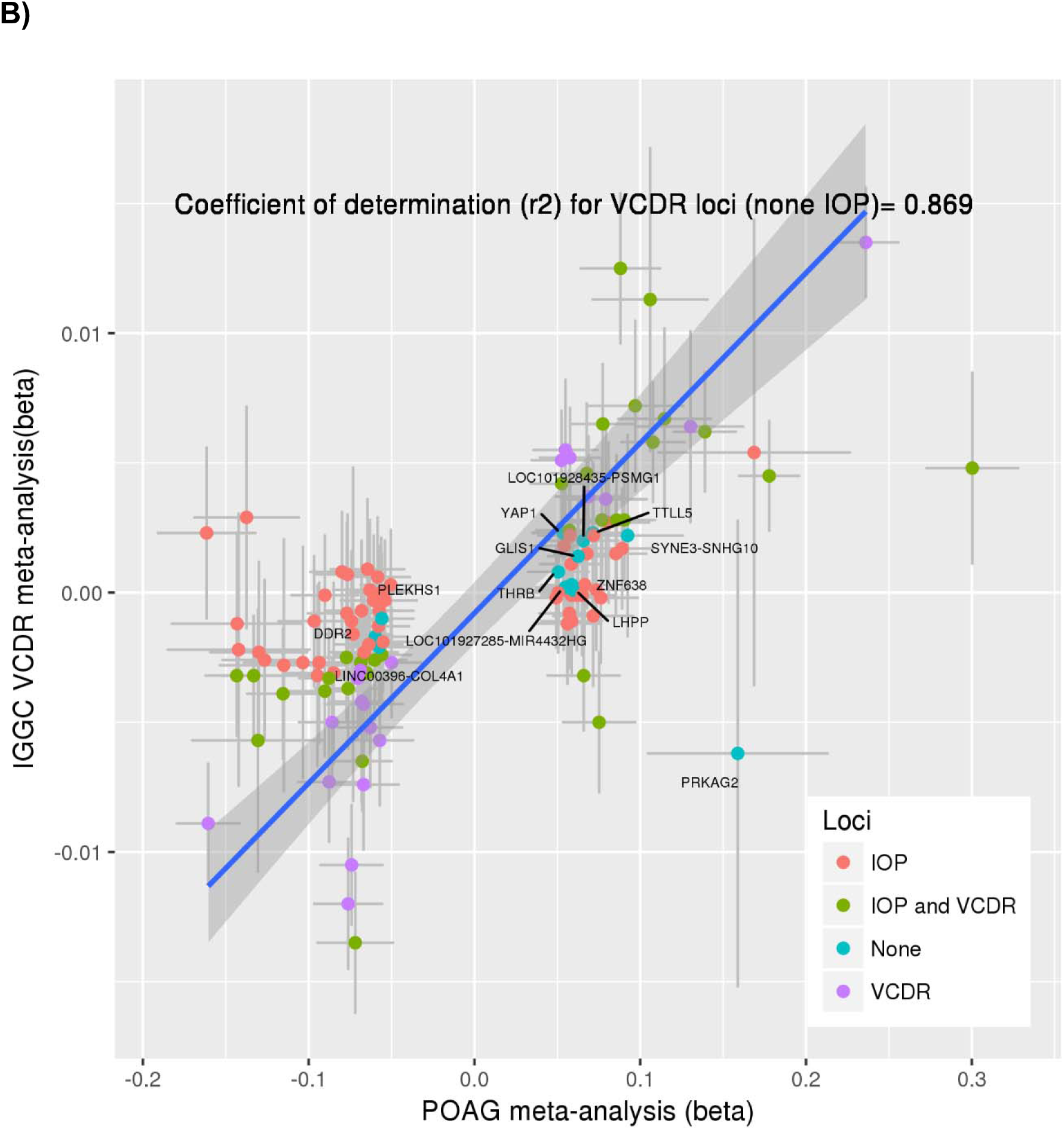
Association of the POAG risk loci with IOP and VCDR. The X-axes show POAG effect estimates in log(OR) scale for the independent genome-wide significant loci obtained from the cross-ancestry meta-analysis. The Y-axes shows the effect estimates for the same SNPs obtained from the meta-analysis of IOP in UKBB + IGGC (mmHg scale; Figure A) and the meta-analysis of VCDR in IGGC (Figure B). Blue line shows the regression line for IOP (Figure A) and VCDR loci (Figure B). Blue dots represent SNPs having P>0.05 for both IOP and VCDR. Horizontal grey bars on each dot represent the 95% confidence intervals (CIs) for the POAG effect estimates, and vertical grey bars shows the 95% CIs for IOP (Figure A) and VCDR (Figure B). Although none of the blue dots show an expected trend of association with IOP in Figure A (their 95% CIs do not overlap with the regression line), the majority of them show a trend of association for VCDR in Figure B.

Another structural endophenotype for POAG in addition to VCDR is retinal ganglion cell-inner plexiform layer (GCIPL) and retinal nerve fiber layer (RNFL) thickness. Of the 127 POAG genome-wide significant lead SNPs, 116 were available in a GWAS of GCIPL/RNFL thickness of the macula (N=31,536; Supplementary Table 6). Of these, 14 loci were associated with GCIPL thickness at nominal significance threshold (P<0.05) and four (*PLEKHA7*, *MAPT*, *LINC01214-TSC22D2*, and *POU6F2*) after Bonferroni correction for the number of tests (P<0.05/116). Similarly, 13 loci were nominally associated with RNFL, and three (*PLEKHA7*, *MAPT*, and *SIX6*) after Bonferroni-correction. These results highlight the possible involvement of *PLEKHA7* and *MAPT* in glaucoma pathogenesis through modulation of retinal thickness.

### Post-GWAS discoveries and functional relevance of the risk loci

We performed gene-based and pathway-based tests using MAGMA^28^ for each ancestry separately, followed by combining P-values across ancestries using Fisher’s combined probability test^29^. We identified 205 genes that passed the gene-based Bonferroni-corrected threshold (P<0.05/20174), corresponding to an additional 11 independent novel risk loci located at least >1Mb apart from the risk loci identified in the single variant-based test. Table 1 presents significant genes within these 11 loci. Expression of the risk genes identified in MAGMA gene-based analysis were significantly enriched in artery and nerve tissues, reflecting the widely recognized neuronal and vascular character of glaucoma (Supplementary Fig 4A and 4B).

**Table 1.**
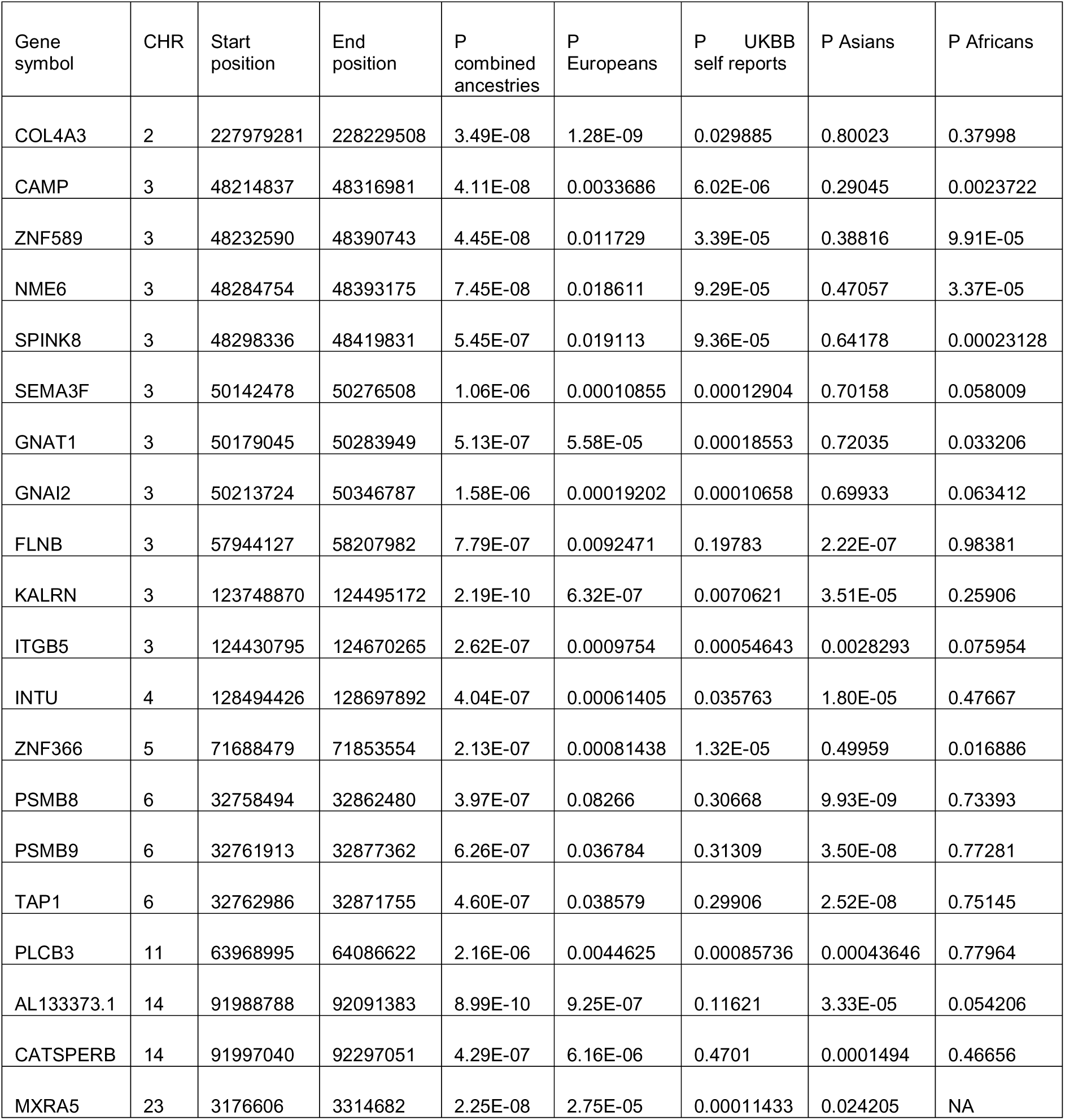
Significant MAGMA gene-based results. The genome-wide significant (P<2.5e-06) gene-based results obtained from MAGMA, which are independent of the risk loci identified in the single variant-based tests (i.e., located at least >1Mb apart from the risk loci identified in the meta-analysis). The corresponding results for each subgroup (Europeans, UKBB self-reports, Asians, and Africans) have also been provided. CHR, chromosome.

| Gene symbol | CHR | Start position | End position | P combined ancestries | P Europeans | P UKBB self reports | P Asians | P Africans |
| --- | --- | --- | --- | --- | --- | --- | --- | --- |
| COL4A3 | 2 | 227979281 | 228229508 | 3.49E-08 | 1.28E-09 | 0.029885 | 0.80023 | 0.37998 |
| CAMP | 3 | 48214837 | 48316981 | 4.11E-08 | 0.0033686 | 6.02E-06 | 0.29045 | 0.0023722 |
| ZNF589 | 3 | 48232590 | 48390743 | 4.45E-08 | 0.011729 | 3.39E-05 | 0.38816 | 9.91E-05 |
| NME6 | 3 | 48284754 | 48393175 | 7.45E-08 | 0.018611 | 9.29E-05 | 0.47057 | 3.37E-05 |
| SPINK8 | 3 | 48298336 | 48419831 | 5.45E-07 | 0.019113 | 9.36E-05 | 0.64178 | 0.00023128 |
| SEMA3F | 3 | 50142478 | 50276508 | 1.06E-06 | 0.00010855 | 0.00012904 | 0.70158 | 0.058009 |
| GNAT1 | 3 | 50179045 | 50283949 | 5.13E-07 | 5.58E-05 | 0.00018553 | 0.72035 | 0.033206 |
| GNAI2 | 3 | 50213724 | 50346787 | 1.58E-06 | 0.00019202 | 0.00010658 | 0.69933 | 0.063412 |
| FLNB | 3 | 57944127 | 58207982 | 7.79E-07 | 0.0092471 | 0.19783 | 2.22E-07 | 0.98381 |
| KALRN | 3 | 123748870 | 124495172 | 2.19E-10 | 6.32E-07 | 0.0070621 | 3.51E-05 | 0.25906 |
| ITGB5 | 3 | 124430795 | 124670265 | 2.62E-07 | 0.0009754 | 0.00054643 | 0.0028293 | 0.075954 |
| INTU | 4 | 128494426 | 128697892 | 4.04E-07 | 0.00061405 | 0.035763 | 1.80E-05 | 0.47667 |
| ZNF366 | 5 | 71688479 | 71853554 | 2.13E-07 | 0.00081438 | 1.32E-05 | 0.49959 | 0.016886 |
| PSMB8 | 6 | 32758494 | 32862480 | 3.97E-07 | 0.08266 | 0.30668 | 9.93E-09 | 0.73393 |
| PSMB9 | 6 | 32761913 | 32877362 | 6.26E-07 | 0.036784 | 0.31309 | 3.50E-08 | 0.77281 |
| TAP1 | 6 | 32762986 | 32871755 | 4.60E-07 | 0.038579 | 0.29906 | 2.52E-08 | 0.75145 |
| PLCB3 | 11 | 63968995 | 64086622 | 2.16E-06 | 0.0044625 | 0.00085736 | 0.00043646 | 0.77964 |
| AL133373.1 | 14 | 91988788 | 92091383 | 8.99E-10 | 9.25E-07 | 0.11621 | 3.33E-05 | 0.054206 |
| CATSPERB | 14 | 91997040 | 92297051 | 4.29E-07 | 6.16E-06 | 0.4701 | 0.0001494 | 0.46656 |
| MXRA5 | 23 | 3176606 | 3314682 | 2.25E-08 | 2.75E-05 | 0.00011433 | 0.024205 | NA |

Pathway analysis identified 21 significant gene-sets surviving the Bonferroni-corrected threshold (P<0.05/10678). These included previously identified pathways such as collagen formation and vascular development^10, 11^, and highlighted novel pathways involved in lipid binding and transportation such as apolipoprotein binding and negative regulation of lipid storage (Table 2). Genes involved in these pathways that demonstrated suggestive association (P<5e-05) with POAG in MAGMA gene-based test are summarized in Supplementary Table 7. Of note, *ABCA1* (an established risk gene for POAG)^6^ and *MAPT* (one of the AD genes associated with POAG) were the most significant POAG-associated genes in the apolipoprotein binding pathway, suggesting that these genes contribute to the development of POAG through disrupting protein interactions within the lipoprotein complex.

**Table 2.** Significant MAGMA gene-set results. The genome-wide significant (P<4.7e-06) pathways obtained from MAGMA. The corresponding results for each subgroup (Europeans, UKBB self-reports, Asians, and Africans) have also been provided.

| Gene set name | P-value Combined | P-value Europeans POAG | P-value UKBB self-report | P-value Asians | P-value Africans |
| --- | --- | --- | --- | --- | --- |
| GO_bp:go_response_to_laminar_fluid_shear_stress | 3.45E-12 | 8.07E-07 | 0.017482 | 4.64E-07 | 0.065546 |
| Curated_gene_sets:matzuk_postimplantation_and_postpartum | 1.30E-09 | 0.023602 | 0.00051684 | 3.14E-06 | 0.0076395 |
| GO_bp:go_phagocytosis_engulfment | 3.98E-08 | 0.026931 | 0.0026067 | 0.00057104 | 0.00033657 |
| Curated_gene_sets:cairo_pml_targets_bound_by_myc_dn | 1.81E-07 | 0.0011074 | 0.00013864 | 0.003352 | 0.14637 |
| Curated_gene_sets:schmidt_por_targets_in_limb_bud_dn | 1.86E-07 | 0.012594 | 0.023226 | 0.00034103 | 0.00077633 |
| GO_bp:go_intracellular_lipid_transport | 3.93E-07 | 0.0078614 | 0.0018918 | 0.0039474 | 0.0031041 |
| GO_bp:go_negative_regulation_of_macrophage_derived_foam_cell_differentiation | 5.54E-07 | 5.15E-05 | 0.023922 | 0.00095691 | 0.22956 |
| GO_bp:go_blood_vessel_morphogenesis | 9.66E-07 | 7.22E-06 | 0.0066377 | 0.012922 | 0.82897 |
| Curated_gene_sets:pasini_suz12_targets_dn | 1.19E-06 | 2.30E-05 | 0.20525 | 0.0032145 | 0.043053 |
| GO_bp:go_regulation_of_endothelial_cell_proliferation | 1.46E-06 | 0.028819 | 2.01E-05 | 0.0053734 | 0.26674 |
| Curated_gene_sets:naba_collagens | 1.47E-06 | 4.77E-06 | 0.0012882 | 0.37129 | 0.3668 |
| Curated_gene_sets:wierenga_stat5a_targets_group2 | 1.65E-06 | 0.00070316 | 0.82593 | 7.09E-05 | 0.023142 |
| Curated_gene_sets:reactome_collagen_formation | 1.86E-06 | 3.51E-06 | 0.0057866 | 0.2652 | 0.20407 |
| GO_bp:go_vasculature_development | 1.98E-06 | 1.04E-05 | 0.012533 | 0.023042 | 0.39096 |
| Curated_gene_sets:piccaluga_angioimmunoblastic_lymphoma_up | 2.11E-06 | 0.00011406 | 0.0025541 | 0.021162 | 0.20531 |
| GO_bp:go_regulation_of_cellular_response_to_growth_factor_stimulus | 2.20E-06 | 0.0036872 | 0.0001049 | 0.0048688 | 0.70563 |
| GO_mf:go_apolipoprotein_binding | 2.22E-06 | 0.027315 | 0.018225 | 2.84E-05 | 0.095355 |
| GO_bp:go_negative_regulation_of_lipid_storage | 2.46E-06 | 0.0014418 | 0.0037001 | 0.0009871 | 0.28762 |
| GO_bp:go_positive_regulation_of_endothelial_cell_proliferation | 3.66E-06 | 0.045899 | 2.63E-05 | 0.015101 | 0.13198 |
| GO_bp:go_positive_regulation_of_cholesterol_efflux | 3.69E-06 | 0.0053921 | 0.0014091 | 0.0018782 | 0.17048 |
| Curated_gene_sets:kinsey_targets_of_ewsr1_flii_fusion_dn | 4.47E-06 | 2.30E-05 | 0.0088699 | 0.06373 | 0.23426 |

To investigate which risk loci are more likely to affect POAG by modulating gene expression, we used two TWAS approaches, Summary Mendelian randomization (SMR)^30^ and MetaXcan^31^. For MetaXcan, we used our POAG cross-ancestry meta-analysis statistics, RNA-seq and genotype data from peripheral retina (EyeGEx)^32^ and 44 GTEX tissues. Following Bonferroni correction for the maximum number of genes tested (N=7,209) in 45 tissues (P<1.5e-07), we identified 100 significant genes, five of which (*AKR1A1*, *DDIT4L*, *LAMTOR3*, *VARS2*, and *ZMAT5*) were located >1Mb apart from (the other loci) identified using single-variant and other gene-based tests in this study (Supplementary Table 8). In a post hoc analysis looking solely at retina, two additional novel genes (*CNTF* and *MPHOSPH9*) were significant (given Bonferroni correction threshold for 6,508 genes in retinal tissue). Of note, *MAPT* and several other genes located within 500kb distance from *MAPT* had suggestive association (P<5e-05) in MetaXcan analyses in multiple tissues. After correction for multiple testing, the predicted expression of *CRHR1* (58.5kb downstream of *MAPT*) was significantly associated with POAG in retina (P=3.9e-08).

Additionally, we integrated our GWAS meta-analysis summary statistics with eQTL data from blood (CAGE eQTL summary data, N=2,765) and retina (EyeGEx eQTL data, N = 406) using SMR. Given that these eQTL data were obtained from people of European descent, we restricted this analysis to our European meta-analysis to ensure that different gene expression and LD structure patterns between ancestries did not influence the SMR findings. In retina and blood, 16 genes passed the SMR significance threshold corrected for the maximum number of 8,516 genes tested in 2 tissues (P<2.9e-06), of which 8 had a P>0.05 in the HEIDI test^30^ implemented in SMR, suggesting that the same association signals drive both gene expression and POAG risk, at these loci (Supplementary Table 9). Although the majority of the risk loci identified through the MetaXcan and SMR approaches were also identified in the meta-analysis, these analyses help with prioritizing the most likely functionally relevant genes. To further identify the most plausible causal genes based on gene expression data, we used the approach implemented in FOCUS^33^, a probabilistic framework that assigns a posterior probability for each gene causally driving TWAS associations in multiple tissues (Supplementary Table 10 summarizes the genes with a posterior probability > 0.6).

The relevance of the identified risk loci was further investigated by examining chromatin interactions in addition to their roles in regulation of gene expression. 72% (92 out of 127) of the lead SNPs or those in high LD (r2>0.8) with the lead SNPs have also been reported to be significant eQTLs (FDR<0.05) in various tissues (Supplementary Table 11). Moreover, the identified risk loci have 34,724 unique significant (FDR<1e-6) chromatin interactions in various tissues/cell lines involving 4,882 genes (Supplementary Fig 5). Of these, 425 genes overlap with the eQTL genes (286 genes if only considering eQTL results for the genome-wide significant SNPs). In addition, SNPs within 105 risk loci identified in this study overlap with one end of these chromatin interactions. Additional support for the pathogenicity of these loci comes from the predicted pathogenicity scores: 20 SNPs had CADD scores >12.37, suggesting that these SNPs have deleterious effects (Supplementary Table 12).^34^ Of the 20, five are coding variants, two of which are novel POAG-associated SNPs (rs61751937 a missense variant in *SVEP1*, and rs8176749 a synonymous variant in *ABO*). Interestingly, *SVEP1* encodes an extracellular matrix protein which is essential for lymphangiogenesis in mice, binds to ANGPT2 (the product of another POAG risk gene identified in this study), and modulates expression of *TEK* and *FOXC2* in Knockout mice^35^. Twenty-four SNPs had RegulomeDB^36^ scores <=3, supporting regulatory roles for these SNPs (Supplementary Table 12).

We investigated the expression of the novel risk genes in eye tissues (Supplementary Table 13 and Supplementary Fig 6). Clustering analysis shows that the majority of the novel genes were expressed in eye tissues (Supplementary Fig 6A). We examined the differential expression of the novel genes in ocular tissues likely to be involved in POAG pathogenesis, namely trabecular meshwork, ciliary body and optic nerve head, and found 50/69 (72%) of the genes differentially expressed in these tissues compared to the other eye tissues tested in this study (Supplementary Table 13 and Supplementary Fig 6B).

At least 16 of the POAG risk genes are targeted by existing drugs, some of which are already in use/clinical trials for several eye or systemic diseases (Supplementary Table 14). The functional relevance of 14 of these 16 drug target genes is supported by the bioinformatic functional analyses we used in this study (i.e. eQTL, chromatin interaction, CADD scores, etc; Supplementary Table 14). We discuss the relevance of some of these drugs in the discussion section below.

### Sex-stratified meta-analysis

We identified a very high genetic correlation (rg=0.99, se=0.06) between POAG in men versus women (European stage 1 and UKBB self-reports combined). We also performed cross-ancestry, sex-stratified meta-analyses using a subset of the overall study with sex-stratified GWAS available (Supplementary Table 1; Supplementary Table 15 and 16; Supplementary Fig 1C and 1D). Only one signal near *DNAH6* appeared to have a female-specific effect (2:84828363[CA], OR=1.6, P=3.28e-09 for women; OR=1.05, P=0.56 for men).

### Subtype-stratified meta-analysis

Finally, we performed cross-ancestry subtype-specific meta-analyses using 3,247 cases and 47,997 controls for NTG (Normal-tension glaucoma defined as glaucoma with IOP <21 mmHg), and 5,144 cases and 47,997 controls for HTG (high-tension glaucoma with IOP > 21 mmHg (Supplementary Table 17 and 18, Supplementary Fig 1E and 1F). All NTG and HTG loci were also genome-wide significant in the overall POAG meta-analysis except for one locus near *FLNB* that was genome-wide significant for NTG (lead SNP rs12494328[A], OR= 1.18, P=1.7e-08) but did not reach the significance threshold in the overall POAG meta-analysis (P=7.5e-07). However, this SNP was genome-wide significant in the 23andMe replication study (rs12494328[A], OR= 1.06, P=1.35-e12), and has previously been associated with optic nerve head changes^27^. Overall, all NTG loci were at least nominally associated (P<0.05) with HTG (and vice versa) except for rs1812974 (top SNP near *ARHGEF12*). Although this SNP had the same direction of effect for NTG, the effect was significantly larger for HTG than NTG (P=0.007). Similarly, several other loci had significantly larger effect on one subtype (e.g., *CDKN2B-AS1*, *FLNB*, and *C14orf39* had larger effects on NTG than HTG) (Supplementary Table 17 and 18). Overall, the genetic correlation between NTG and HTG was estimated to be 0.58 (se=0.08) using LD score regression and the meta-analysis summary data from Europeans.

## Discussion

In this large multi-ethnic meta-analysis for POAG, we identified 127 risk loci for POAG, 69 of which reached the genome-wide significant level for the first time. We also identified additional risk loci using gene-based tests and highlighted genetic pathway involved in the pathogenesis of POAG. We observed relatively consistent genetic effects for POAG across ancestries. We identified a direct link between POAG risk loci and those known to affect AD. The risk loci include genes that are highly expressed in relevant eye tissues, nerves, arteries, as well as tissues enriched with these components. Functional relevance of the identified risk loci were further supported by eQTL and chromatin interaction data.

We identified a strong correlation between the POAG effect sizes of genome-wide significant SNPs, as well as all the SNPs throughout the genome, across Europeans, Asians, and Africans. Although previous studies have suggested that the genetic architecture of POAG might differ between Africans and Europeans^37^, we observed a high correlation (r ∼ 0.7) between effect sizes of the POAG risk loci in Europeans and Africans (Supplementary Fig 2C). Despite the broad concordance, a subset of the known (e.g. *TMCO1*, *COL11A1, CASC20, BICC1,* and *LMO7*) and novel (e.g. *BCAS3*, and *OVOL2*) loci showed significant difference (P<0.05) of effect sizes between Europeans and Africans. There were several examples of loci in this study where including African GWASs could help with fine mapping the GWAS signals in Europeans and Asians (as the LD blocks were relatively shorter in Africans). However, this was not applicable to the majority of the loci partly due to the lower statistical power of our African studies.

This study identified a genetic correlation throughout the genome between POAG and AD, with overlap of three POAG genome-wide significant risk loci (*MAPT*, *CADM2*, and *APP*) with those also implicated in AD risk^21–23^. Common variants within or near *MAPT* have been associated with AD^21, 22^, but mutations in *MAPT* have been found to be even more prominent in to frontotemporal dementia^38^. It is of note that the most significant POAG SNP (rs242559) in this region is located within an intron of *MAPT* and is in high LD (r2=0.89) with the common variant associated with AD near *MAPT,* within *ARL17B*. rs242559 was also associated with retinal GCIPL and RNFL thickness in UKBB. Interestingly, RNFL thinning is associated with increased cognitive decline in UKBB^39, 40^. rs242559 has also been significantly associated with many other traits including general cognitive ability and Parkinson’s disease (https://genetics.opentargets.org/variant/17_45948522_C_A), further supporting the functional importance of this variant.

For *CADM2,* although the lead SNPs for POAG (rs13101042) was not in high LD with the AD lead SNP (rs71316816; LD r2=0.03), rs71316816 showed some evidence of association with POAG (P=6e-04). In addition, SNPs in high LD with rs13101042 have been shown to be associated with cognitive function traits; for example, rs2220243 which is in LD r2=0.8 with rs13101042 is genome-wide significant (P=4e-08) for mathematical ability^41^.

Finally, mutations in *APP* lead to early-onset AD by increasing the production of toxic forms of amyloid β (Aβ) peptide. Here, our POAG GWAS identified common variants near *APP,* with the POAG SNP also showing an association with IOP (P=0.00004). Interestingly, a POAG GWAS in individuals of African descent identified a variant in Amyloid Beta Precursor Protein Binding Family B Member 2 (*APBB2*)^43^; the reported SNP at this locus (rs59892895) was also associated with POAG in our African meta-analysis (rs59892895[C], OR=1.27, P=1.0e-07, although there is sample overlap between the two studies). The associated allele at rs59892895 in Africans has allele frequency <0.1% in Europeans and Asians. Although the causal variants for POAG may not be the same variants associated with AD in *CADM2* and *APP*, further fine mapping and functional studies to confirm the biological relevance of those genes for POAG may advance and inform the physiologic bases of both POAG and AD.

The gene-sets enriched for the risk loci identified in this study indicate two major pathogenic mechanisms for POAG: 1) vascular system defects, mainly the molecular mechanisms contributing to blood vessel morphogenesis, vasculature development, and regulation of endothelial cell proliferation, and; 2) lipid binding and transportation - mainly the molecular mechanisms involved in intracellular lipid transport, apolipoprotein binding, negative regulation of lipid storage, and positive regulation of cholesterol efflux. Involvement of the vascular system in the pathogenesis of POAG is further supported by our results showing enrichment of the expression of the POAG risk genes in arteries and vessels. Molecular targets in these pathways can be potential candidates for treatment of POAG. Two of the significant gene-sets in this study (phagocytosis engulfment and negative regulation of macrophage derived foam cell differentiation) suggest an important role of immune system defects in increasing the risk of POAG.

Several genes within the identified POAG risk loci are targets of some currently approved drugs. For instance, COL4A1 is targeted by ocriplasmin, a collagen hydrolytic enzyme that is currently used to treat vitreomacular adhesion (adherence of vitreous to retina). This drug can degrade the structural proteins including those involved in vitreoretinal surface^44^. Ocriplasmin is currently under clinical trials for several eye conditions including macular degeneration, diabetic macular edema, macular hole, and retinal vein occlusion. Some other drug candidates are currently under consideration for treating dementia and cardiovascular diseases. For example, acitretin is a retinoid receptor agonist targeting RARB which has been considered for treatment of AD in ongoing clinical trials. Dipyridamole, a 3’,5’-cyclic phosphodiesterase inhibitor, targets PDE7B and is currently under clinical trials for diseases such as stroke, coronary heart disease, ischemia reperfusion injury, and internal carotid artery stenosis. Given that our pathway analyses highlighted the involvement of vasculature development and blood vessel morphogenesis in pathogenesis of POAG, these drugs might be capable of controlling the abnormal angiogenesis and vasculature processes occurring in POAG. Further studies to confirm the functionality of these POAG risk genes *in-vivo* and *in-vitro* may support the suitability of repurposing these drugs as novel treatments for POAG.

This study has several strengths and limitations. The main strength includes identification of risk loci contributing to the development of POAG across ethnic groups, as opposed to most POAG GWAS that have focused on individuals of European descent. We showed that combining GWAS data across ancestries increases the power of gene mapping for POAG. Another strength is the integration of GWAS, gene expression, and chromatin interaction data to investigate the functional relevance of the identified loci, as well as to identify the most plausible risk genes.

A limitation of this study is that although the majority of the cases were confirmed POAG, our data included >7,200 glaucoma cases from the UKBB obtained through self-reports. However, we observed a very high concordance between the GWAS results for clinically validated cases versus self-report. Additionally, the vast majority of our results replicated in self-report data from 23andMe. Finally, although we investigated the functional relevance of the identified risk loci using bioinformatic analyses, we did not confirm their functionality *in-vitro* and *in-vivo*. Further studies to investigate the biological roles of these risk loci with respect to POAG pathogenesis in relevant eye tissues will further shed light on the molecular etiology of POAG.

In conclusion, this study identified a strong cross-ancestry genetic correlation for POAG between Europeans, Asians, and Africans, and identified 127 genome-wide significant loci by combining GWAS results across these ancestries. We found novel and interesting biology including a genetic link of POAG with AD. Further dissecting the shared biology behind the genetic link with AD is warranted to reveal whether similar prevention and management strategies can be applied. Finally, some genes within POAG risk loci are targets of currently approved drugs used for the treatment of other diseases, making them potential candidates for drug repurposing for the treatment of POAG.

## Online Methods

### Study design and participants

We obtained 34,179 POAG cases and 349,321 controls including participants of European, Asian, and African descent from 21 independent studies across the world. Number of cases and controls, genotyping platform, and distribution of age and sex for each study are summarized in Supplementary Table 1. The phenotype definition and additional details for each study are provided in Supplementary Notes. For most of the studies, we restricted glaucoma to POAG based on the ICD9/ICD10 criteria. However, considering that POAG constitutes the majority of glaucoma cases in Europeans^45^, we also included 7,286 glaucoma self-reports from UK Biobank to replicate findings from the ICD9/ICD10 POAG meta-analysis in Europeans and to maximize the statistical power of the final stage meta-analysis (please see below). Informed consent was obtained from all the participants, and ethics approval was obtained from the ethics committee of all the participating institutions.

We performed a three-stage meta-analysis to combine GWAS data from the participating studies. In the first stage, we conducted a meta-analysis of the POAG GWAS in Europeans (16,677 POAG cases vs 199,580 controls). In the second stage, we performed a meta-analysis of POAG GWAS in Asians (6,935 cases vs 39,588 controls) and Africans (3,281 cases vs 2,791 controls) (Supplementary Table 1). These data along with a GWAS of 7,286 self-report glaucoma cases vs 107,362 controls of European descent from UKBB were used to validate the findings from the first stage. The UKBB self-report GWAS was completely independent of the UKBB IC9/ICD10 POAG GWAS; all the UKBB POAG cases and controls from the first stage, as well as their relatives (Pi hat >0.2), were removed from the self-report GWAS dataset. In the third stage, we combined the results from stage 1 and 2 to increase our statistical power to identify POAG risk loci across ancestries.

To investigate sex-specific loci for POAG, we also conducted a meta-analysis of POAG in males and females separately. For this analysis, we had GWAS data from a subset of the overall POAG meta-analysis, including 10,775 cases vs 123,644 controls for males, and 10,977 cases vs 144,606 controls for females (Supplementary Table 1). Similarly, to identify risk loci for the HTG and NTG subtypes, we performed a subtype-specific meta-analysis using 3,247 NTG cases vs 47,997 controls, and 5,144 HTG cases vs 47,997 controls.

### Quality control (QC) and imputation

Study-specific QC and imputation details have been provided in Supplementary Notes. Overall, SNPs with >5% missing genotypes, minor allele frequency (MAF) < 0.01, and evidence of significant deviations from Hardy-Weinberg equilibrium (HWE) were excluded. In addition, individuals with >5% missing genotypes, one of each pair of related individuals (detected based on a p-hat>0.2 from identity by descent calculated from autosomal markers), and ancestry outliers from each study (detected based on principal component analysis including study participants and reference samples of known ancestry) were excluded from further analysis (for more details please see Supplementary Notes).

Imputation for studies involving participants of European descent was performed in Minimac3 using the Haplotype Reference Consortium (HRC) r1.1 as reference panel through the Michigan Imputation Server^46^. However, for a study of Finnish population (FinnGen study), whole-genome sequence data from 3,775 Finnish samples were used as reference panel for better population-specific haplotype matching, which results in a more accurate imputation. For studies involving Asian and African participants, imputation was performed using the 1000 Genomes samples of relevant ancestry. SNPs with MAF > 0.001 and imputation quality scores (INFO or r^2^)> 0.3 were taken forward for association analysis.

### Association testing

Association testing was performed assuming an additive genetic model using dosage scores from imputation, adjusting for age, sex, and study-specific principal components as covariates, using software such as PLINK^46, 47^, SNPTEST^48, 49^, SAIGE^50^, EPACTS (https://genome.sph.umich.edu/wiki/EPACTS), and RVtests^51^. For studies with a large number of related individuals, mixed-model association testing was performed to account for relatedness between people. For the X chromosome analysis, we used the following approach to allow for dosage compensation: females were coded as 0 (homozygous for non-effect allele), 1 (heterozygous), and 2 (homozygous for effect allele) while males were coded as 0 (no effect allele) and 2 (one effect allele). The covariates were the same as for the association testing for the autosomes except removing sex.

To confirm the validity of combining GWAS results across populations comprising different ancestries, we estimated the genome-wide genetic correlation for POAG between the populations of European, Asian, and African descent participated in this study. For this purpose, we used Popcorn^20^, a toolset that provides estimates of genetic correlation while accounting for different genetic effect and LD structure between ancestries. For this analysis, the LD scores for each ancestry population were estimated using the 1000G populations (Europeans, Asians, and Africans), and SNPs were filtered based on the default MAF=0.05 in Popcorn.

We performed within and between ancestry meta-analyses using a fixed-effects inverse-variance weighting approach in METAL^52^ using SNP effect point estimates and their standard errors. The presence of heterogeneity between SNP effect estimates across studies were investigated using the Cochran’s Q test implemented in METAL. To identify multiple independent risk variants within the same locus using GWAS summary statistics obtained from the meta-analysis, we used the Conditional and Joint (COJO) analysis implemented in GCTA^53^. Q-Q and Manhattan plots were created in R, and regional association plots in LocusZoom^54^.

We used the univariate LD score regression^18^ intercept for each study separately as well as for the meta-analysed results to ensure that the test statistics did not include model or structural biases such as population stratification, cryptic relatedness, and model misspecification. To investigate the genetic correlation between POAG and AD, we used bivariate LD score regression^25^ using our POAG meta-analysis in Europeans and 71,880 AD cases and 383,378 controls of European descent obtained from meta-analysis of Psychiatric Genomics Consortium (PGC-ALZ)^55^, the International Genomics of Alzheimer’s Project (IGAP)^56^, and the Alzheimer’s Disease Sequencing Project (ADSP)^57^.

The association of the POAG risk loci identified in this study with its major endophenotypes, IOP and VCDR, was investigated using summary statistics from a recent GWAS meta-analysis for IOP (N=133,492)^11^, and the IGGC GWAS meta-analysis for VCDR (N=23,899)^27^.

### 23 andMe replication

We validated the genome-wide significant risk loci from our cross-ancestry meta-analysis (127 independent SNPs) and subtype analyses (7 independent SNPs) in a subset of 23andMe research participants of European descent comprising 43,254 POAG cases and 1,471,118 controls. POAG cases were defined as those who reported glaucoma, excluding those who reported angle-closure glaucoma or other types of glaucoma. Controls did not report any glaucoma. Association testing was performed using logistic regression assuming an additive genetic model, adjusting for age, sex, top five principal components, and genotyping platform as covariates.

### Gene-based and pathway-based tests

Gene-based and gene-set (pathway) based tests were performed using the approach implemented in MAGMA^28^. We performed this analysis for each ancestry separately, and the P values were then combed across ancestries using Fisher’s combined probability test. The significance threshold for gene-based test was set to P<0.05/20174, and for pathway-based tests to P<0.05/10678, accounting for the maximum number of independent genes/pathways tested. In addition, MAGMA was used to investigate the enrichment of the expression of the significant risk genes in GETX v6 tissues (P<1e-03 accounting for 53 tissues tested).

To identify loci whose effect on POAG risk is due to modulation of gene expression, we also used alternative gene-based tests that integrate GWAS summary statistics with eQTL data throughout the genome (transcriptome-wide association study (TWAS)-based approaches). For this purpose, we used MetaXcan^31^ and SMR^30^. MetaXcan uses GWAS summary statistics to impute the genetic component of gene expression in different tissues based on a reference eQTL panel. We used the EyeGEx eQTL data from retina^32^ as well as 44 GTEX tissues as reference eQTL data for this study. The GWAS input for this analysis included the summary statistics obtained from the cross-ancestry meta-analysis. We set the significance threshold to Bonferroni-corrected P<1.5e-07, accounting for the maximum number of genes tested (N=7,209) in 45 tissues.

SMR uses a Mendelian Randomization framework to identify genes whose expression is likely modulated through the same variants associated with the outcome of interest (POAG). For the SMR analysis, we used the following eQTL data: CAGE eQTL summary data from blood (N=2,765) and EyeGEx eQTL data from retina (N = 406). The SMR significance threshold was set to P<2.9e-06, accounting for the maximum number of 8,516 genes tested in 2 tissues. A heterogeneity P>0.05 from the HEIDI test implemented in SMR implies that we cannot reject the null hypothesis that a single causal variant is likely to affect both gene expression and POAG risk for these loci.

### Gene expression

RNA was extracted from the corneal layers, trabecular meshwork, ciliary body, retina, and optic nerve tissues from 21 healthy donor eyes of 21 individuals. After quality control, the RNA was sequenced using Illumina NextSeq® 500 (San Diego, USA). Trimgalore (v0.4.0) was used to trim low-quality bases (Phred score < 28) and reads shorter than 20 bases after trimming were discarded. Data analysis was done with edgeR (version 3.22.5)^58^. Only genes expressed a minimum of 10 times (1.5 counts per million) in at least 5 dissected tissues were kept. The RNA count libraries were normalised using trimmed mean of M-values method^59^. Two-group differential expression analysis was done via negative binomial generalised linear model in edgeR^60^. The RNA expression in ciliary body, trabecular meshwork, and optic nerve head which are involved in aqueous production, drainage and principal site of glaucoma injury respectively was compared to the remaining eye tissues.

### Drug targets

We used Open Targets^61^ to search for drugs currently in use or in clinical trials for treating other ocular or systemic diseases that target the POAG risk genes identified in this study. These drugs can be potentially repurposed as novel treatment for POAG, owing to *in-vivo* and *in-vitro* confirmation of the functionality of the target genes in the pathogenesis of POAG.

### Bioinformatic functional analyses

The bioinformatic functional analysis to investigate the functional relevance of the identified risk loci for POAG were performed through FUMA^62^ using the following dataset/toolsets: GTEX eQTL v6^24^; Blood eQTL browser^63^; BIOS QTL Browser^64^; BRAINEAC^65^; RegulomeDB^36^; CADD^34^; ANNOVAR^66^; and Hi-C data from 21 tissue/cell types from GSE87112^67^, PsychENCODE^68^, Giusti-Rodriguez et al 2019 (https://www.biorxiv.org/content/10.1101/406330v2), and FANTOM5 Human Enhancer Tracks (http://slidebase.binf.ku.dk/human_enhancers/presets).

## Acknowledgements

This work was conducted using the UK Biobank Resource (application number 25331) and publicly available data from the International Glaucoma Genetics Consortium. The UK Biobank was established by the Wellcome Trust medical charity, Medical Research Council (UK), Department of Health (UK), Scottish Government, and Northwest Regional Development Agency. It also had funding from the Welsh Assembly Government, British Heart Foundation, and Diabetes UK. The eye and vision dataset has been developed with additional funding from The NIHR Biomedical Research Centre at Moorfields Eye Hospital and the UCL Institute of Ophthalmology, Fight for Sight charity (UK), Moorfields Eye Charity (UK), The Macula Society (UK), The International Glaucoma Association (UK) and Alcon Research Institute (USA). This work was also supported by grants from the National Health and Medical Research Council (NHMRC) of Australia (#1107098; 1116360, 1116495, 1023911), the Ophthalmic Research Institute of Australia, the BrightFocus Foundation, UK and Eire Glaucoma Society and Charitable Funds from Royal Liverpool University Hospital, and International Glaucoma Association-Royal College of Ophthalmologists. SM, JEC, KPB, and AWH are supported by NHMRC Fellowships. We thank Scott Wood, John Pearson and Scott Gordon from QIMR Berghofer for support. The NEIGHBORHOOD consortium is supported by NIH grants P30 EY014104, R01 EY015473 and R01 EY022305. The EPIC-Norfolk study (DOI 10.22025/2019.10.105.00004) has received funding from the Medical Research Council (MR/N003284/1 and MC-UU_12015/1) and Cancer Research UK (C864/A14136). The genetics work in the EPIC-Norfolk study was funded by the Medical Research Council (MC_PC_13048). We are grateful to all the participants who have been part of the project and to the many members of the study teams at the University of Cambridge who have enabled this research. We thank the research participants and employees of 23andMe who contributed to this study.

## Author contributions

PG, EJ, PH, APK, SP, AWH, RPJI, HC, JNCB, PB, TY, VV, AHJT, JK, RBM, KSN, RL, MS, NA, AJC, RH, AP, LJC, CC, ENV, GT, YS, MY, TN, JR, EB, KH, YK, MA, YM, PJF, PTK, JEM, NGS, PK, JHK, CCP, FP, PM, AJL, AP, CvD, JH, CH, LRP, CCK, MH, CK, DAM, MK, TA, JC, SM, and JW were involved in data collection and contributed to genotyping. PG, PH, XH, JSO, AS, RPJI, HC, AQ, NSJ, PB, AI, JK, SU, MS, GT, HC, XW, AA, MA, and JHK were involved in data analysis. PG, SM, and JW wrote the first draft of the paper. EJ, APK, PJF, PTK, AJL, AP, MH, CCK, DAM, MK, TA, JC, SM, and JW designed the study and obtained the funding.

## Data sharing

UK Biobank data are available through the UK Biobank Access Management System https://www.ukbiobank.ac.uk/.

## Supplementary Tables

**Supplementary Table 1. Study overview of the meta-analysis.** This table summarises the sample size for the overall POAG, sex-stratified, and subtype-stratified analyses for each contributing site.

**Supplementary Table 2. Independent genome-wide significant loci in Europeans.** This table provides summary statistics for the most significant genome-wide associated SNPs from the GWAS meta-analysis of POAG in Europeans. Independent loci were identified through the Conditional and Joint (COJO) analysis in GCTA. The corresponding GWAS summary statistics in UKBB (glaucoma self-report), Asian, and African studies have also been provided. Two loci (*PKHD1* and *TFEC-TES*) had suggestive association in the meta-analysis (P<5e-06) but became significant after performing the COJO analysis. CHR, chromosome; EUR, European; P_COJO, P-value from the Conditional and Joint analysis; Novel: risk loci that, for the first time, become genome-wide significant (P<5e-08) for POAG (some of these loci were genome-wide significant for IOP or VCDR previously, but did not reach genome-wide significance for POAG risk).

**Supplementary Table 3.** Genome-wide significant Loci in Asians and Africans. ASN, Asian; EUR, European; AFR, African.

**Supplementary Table 4. Independent genome-wide significant loci in the GWAS meta-analysis of POAG across ancestries.** This table provides summary statistics for the most significant SNPs from the GWAS meta-analysis of POAG in Europeans, Asians, and Africans. The corresponding GWAS summary statistics for each subgroup (Europeans, UKBB self-reports, Asians, and Africans) as well as 23andMe (the replication data; GWAS of 43,254 glaucoma cases vs 1,471,118 controls) have also been provided. CHR, chromosome; meta, meta-analysis; EUR, European; Novel: risk loci that, for the first time, become genome-wide significant (P<5e-08) for POAG (some of these loci were genome-wide significant for IOP or VCDR previously, but did not reach genome-wide significance for POAG risk).

**Supplementary Table 5. The association of the POAG risk SNPs with IOP and VCDR.** The results are from GWAS meta-analysis of N=133,492 IOP and N=23,899 VCDR. CHR, chromosome; meta, meta-analysis. *rs6755023 in LD r2=0.98 with rs200621439. **rs327736 in LD r2=0.90 with rs59101260. ***rs35220810 in LD r2=1.0 with rs35740987. ****rs62063276 in LD r2=0.95 with rs114919114.

**Supplementary Table 6. The association of the POAG risk SNPs with macular thickness.** Freq, frequency of the effect allele; info, imputation quality score; GCIPL, ganglion cell-inner plexiform layer; RNFL, retinal nerve fiber layer.

**Supplementary Table 7. The most significant POAG genes within the significant pathways.** Genes within the significant pathways identified by MAGMA that are at least suggestively associated (P<5e-05) with POAG in the MAGMA gene-based test.

**Supplementary Table 8. The significant POAG genes identified by MetaXcan.** eQTL data from retina (EyeGEx study, N = 406) and 44 GTEX tissues were used for this analysis. Var_g, variance of genes expression; pred_perf_r2, R2 of the predicted gene expression to gene’s measured transcriptome (prediction performance); pred_perf_pval, P-value of the prediction performance; pred_perf_qval, q-value of the prediction performance; n_snps_used, number of SNPs used from GWAS; n_snps_in_cov, number of SNPs in the covariance matrix; n_snps_in_model; number of SNPs used in the model.

**Supplementary Table 9. The significant POAG genes identified by SMR.** The POAG meta-analysis results in Europeans were integrated with eQTL data from blood (CAGE study, N=2,765) and retina (EyeGEx study, N = 406) using the SMR approach. A1, Effect allele; A2, Other allele; Freq, frequency of the effect allele.

**Supplementary Table 10. Gene prioritisation by FOCUS.** Tx_start, transcription start site; tx_stop, transcription end site; cv.R2, cross-validation predictive R Squared; cv.R2.pval, P-value of cross-validation; twas_z, marginal Z score from TWAS; pip, marginal posterior inclusion probability; in_cred_set, whether or not model is included in the credible set.

**Supplementary Table 11. eQTL results for the POAG risk SNPs.** This table shows eQTL results for the most significant POAG risk SNPs or those in high LD (r2>0.8) with the most significant SNPs.

**Supplementary Table 12.** CADD and RegulomeDB pathogenicity scores for the POAG risk SNPs.

**Supplementary Table 13. Expression of the previously unreported POAG risk genes in eye tissues.** Diff_express, differential expression in trabecular meshwork, ciliary body, and optic nerve head vs the rest of the eye tissues presented in the table; Diff_express_P, Differential expression P-value (Bonferroni adjusted).

**Supplementary Table 14. Candidate drugs for the POAG risk genes.** pLI, this is the pLI score from the ExAC database showing the probability of being loss-of-function intolerant. The higher the score, the more intolerant the variant; ncRVIS, this is the non-coding residual variation intolerance score. The higher the score, the more intolerant the variant; posMapSNPs, this shows the number of SNPs which were mapped to genes based on FUMA positional mapping; posMapMaxCADD, the maximum CADD score of SNPs which were mapped to genes based on FUMA positional mapping; eqtlMapSNPs, the number of SNPs which were mapped to genes based on FUMA eQTL mapping; eqtlMapminP, the minimum eQTL P-value of the mapped SNPs by FUMA eQTL mapping; eqtlMapminQ, the minimum eQTL FDR of the mapped genes by FUMA eQTL mapping; ciMap, this indicates whether or not the gene is mapped by FUMA chromatin interaction mapping; ciMapts, this shows which tissues or cell types if there is a “yes” to ciMap column.

**Supplementary Table 15. The top genome-wide significant SNPs for the male-stratified analysis.** The corresponding statistics for the female-stratified analysis as well as the overall POAG meta-analysis has also been shown. The SNP presented in bold was not genome-wide significant in the meta-analysis. CHR, chromosome; meta, meta-analysis.

**Supplementary Table 16. The top genome-wide significant SNPs for the female-stratified analysis.** The corresponding statistics for the male-stratified analysis as well as the overall POAG meta-analysis has also been shown. The SNPs presented in bold were not genome-wide significant in the meta-analysis. CHR, chromosome; meta, meta-analysis.

**Supplementary Table 17. The top genome-wide significant SNPs for the NTG-stratified analysis.** The corresponding statistics for the HTG-stratified analysis as well as the overall POAG meta-analysis has also been shown. The SNP presented in bold was not genome-wide significant in the meta-analysis. CHR, chromosome; meta, meta-analysis; P_diff, P-value for the difference between the estimated effect sizes for NTG vs HTG.

**Supplementary Table 18. The top genome-wide significant SNPs for the HTG-stratified analysis.** The corresponding statistics for the NTG-stratified analysis as well as the overall POAG meta-analysis has also been shown. CHR, chromosome; meta, meta-analysis; P_diff, P-value for the difference between the estimated effect sizes for NTG vs HTG.

