## Supplementary Figures for "A large cross-ancestry meta-analysis of genome-wide association studies identifies 69 novel risk loci for primary open-angle glaucoma and includes a genetic link with Alzheimer’s disease"

**Supplementary Figure 1. QQ and Manhattan plots.**

**A) European meta-analysis.**

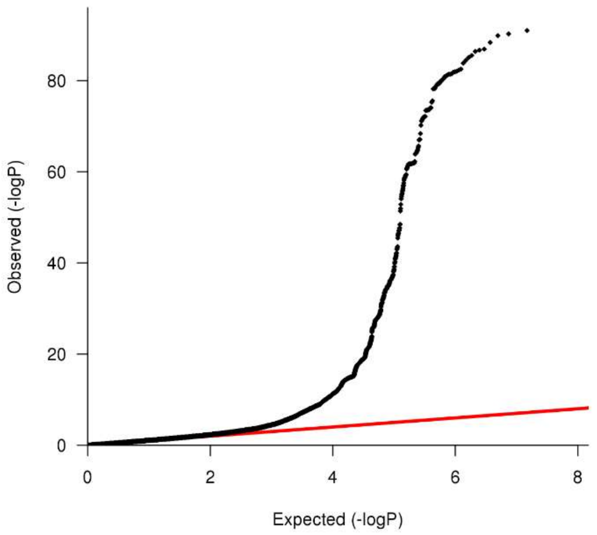


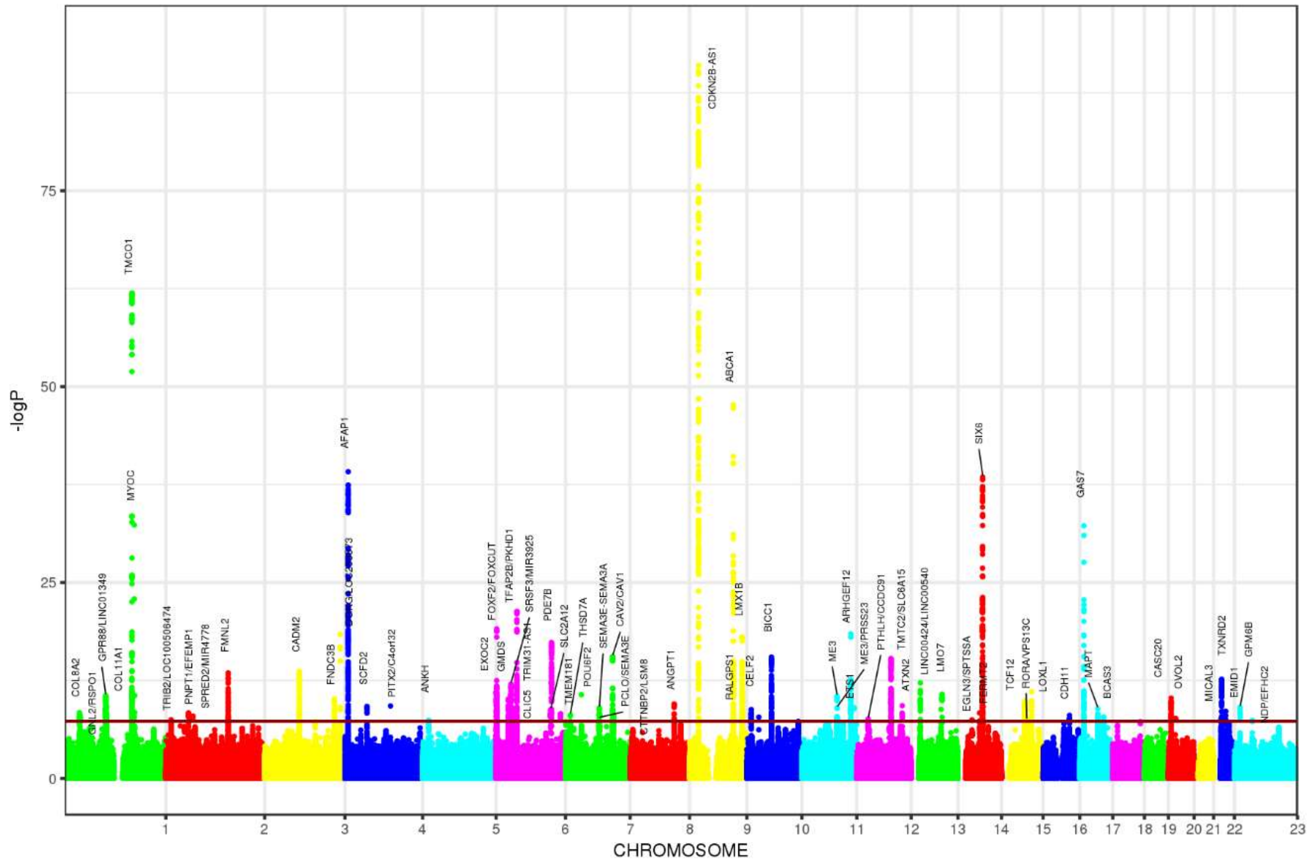

**B) Cross-ancestry meta-analysis.**

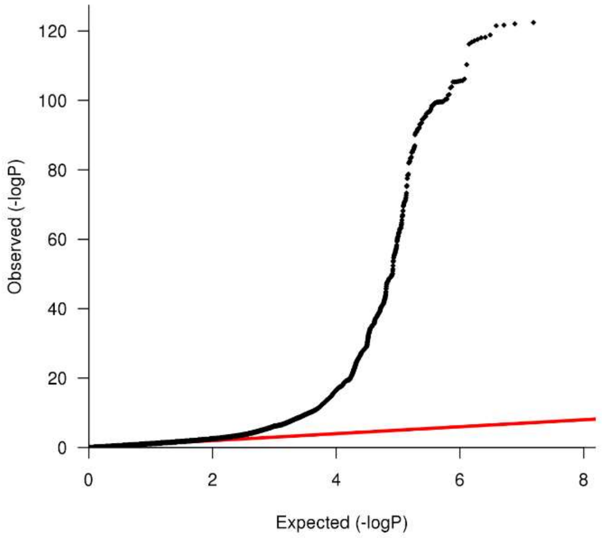


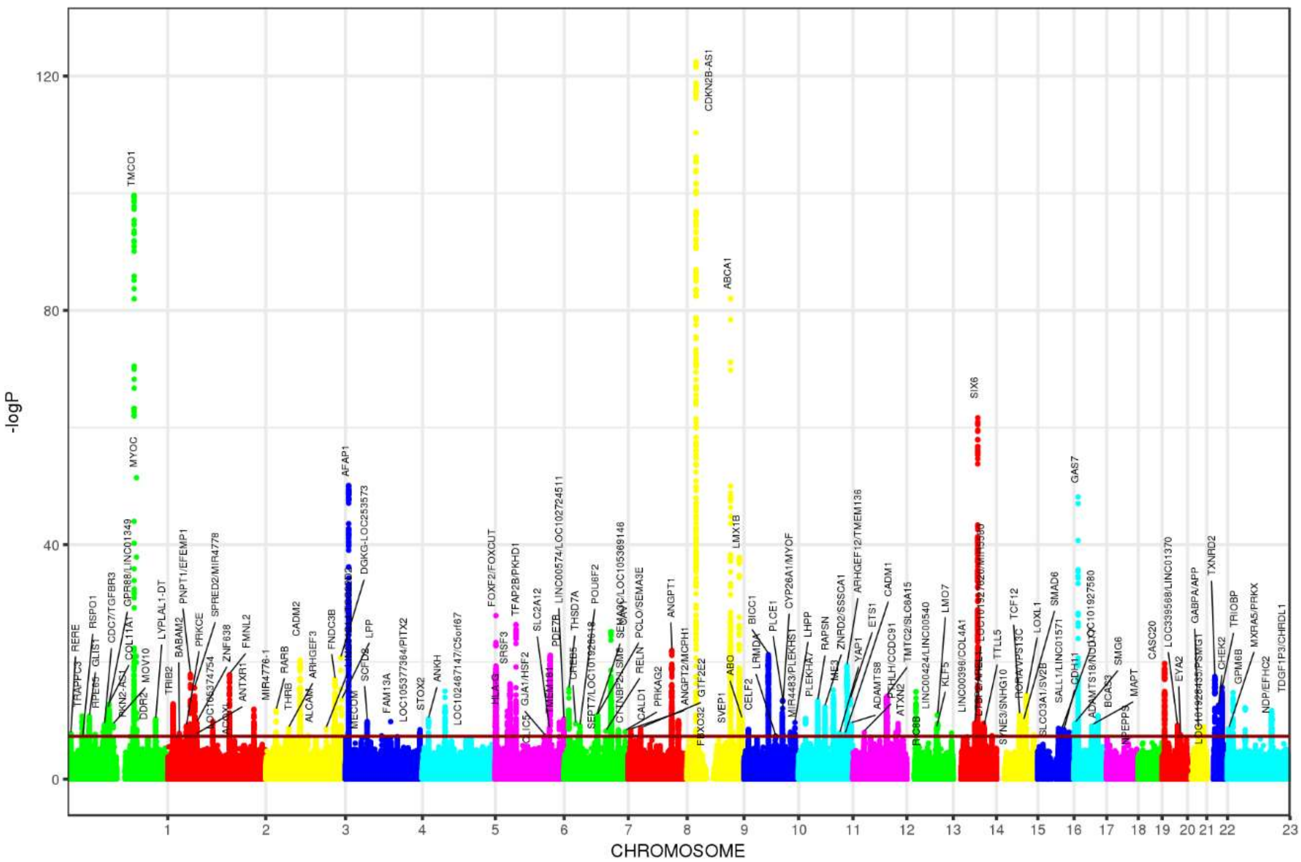

#### C) Male-stratified meta-analysis.

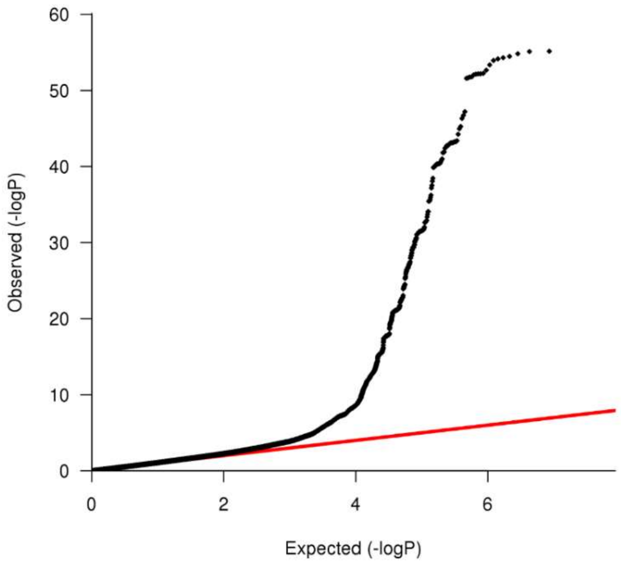


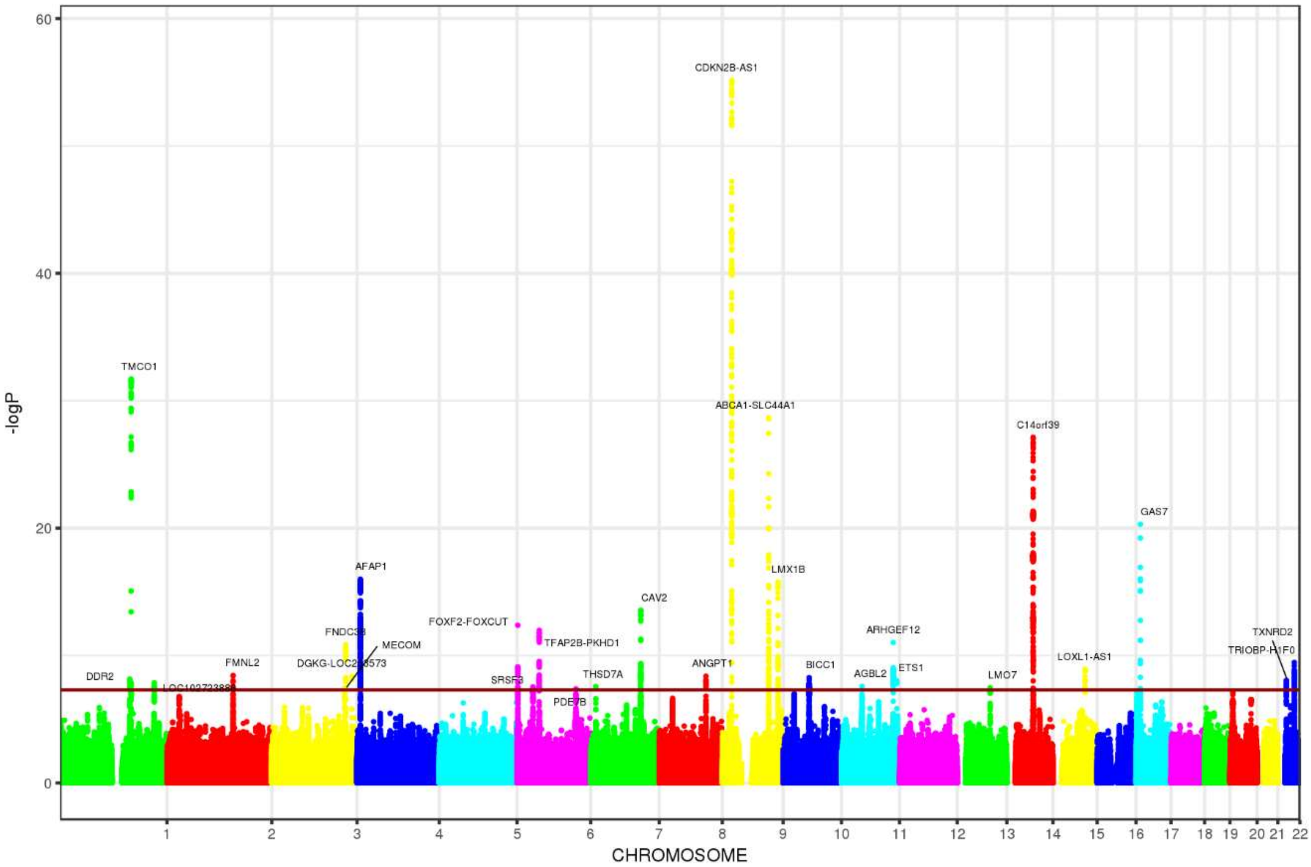

**D) Female-stratified meta-analysis.**

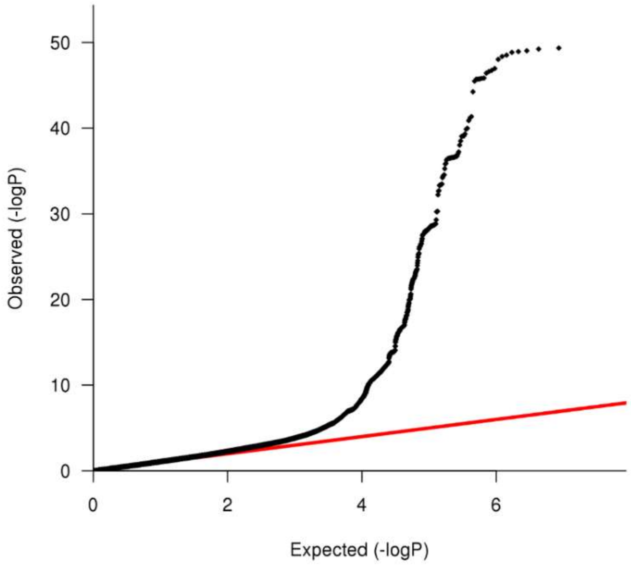


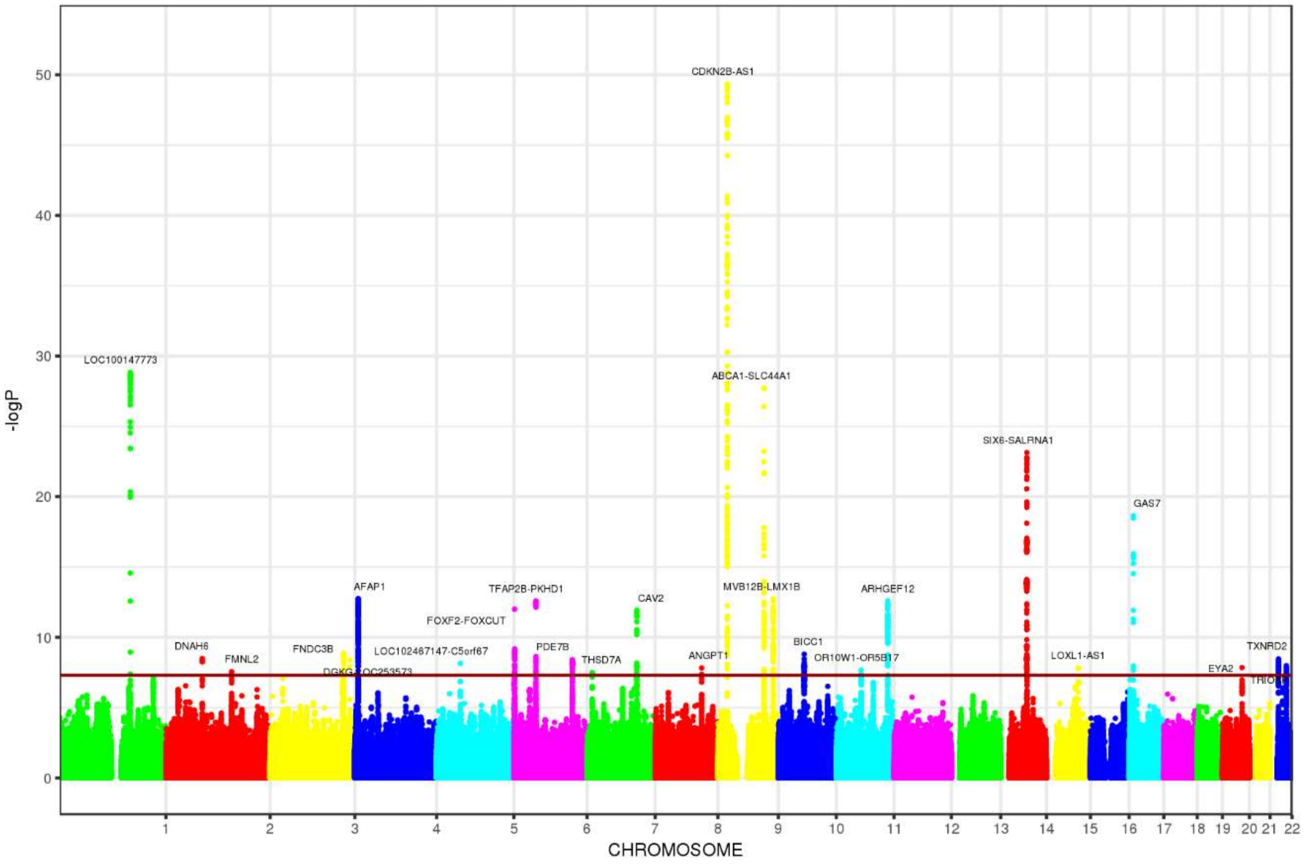

**E) NTG-stratified meta-analysis.**

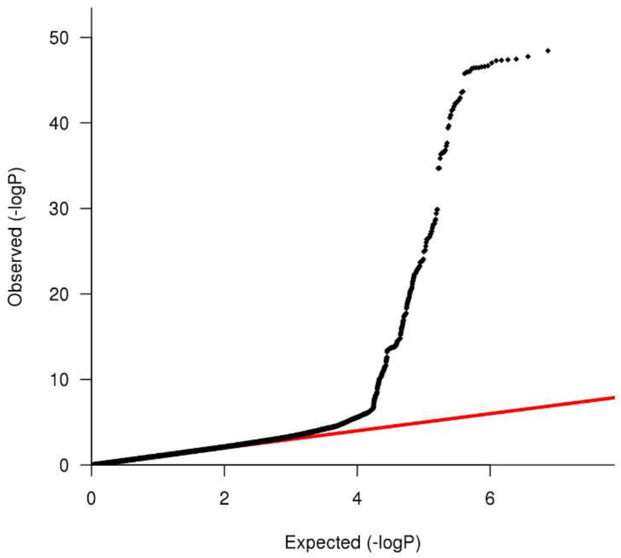


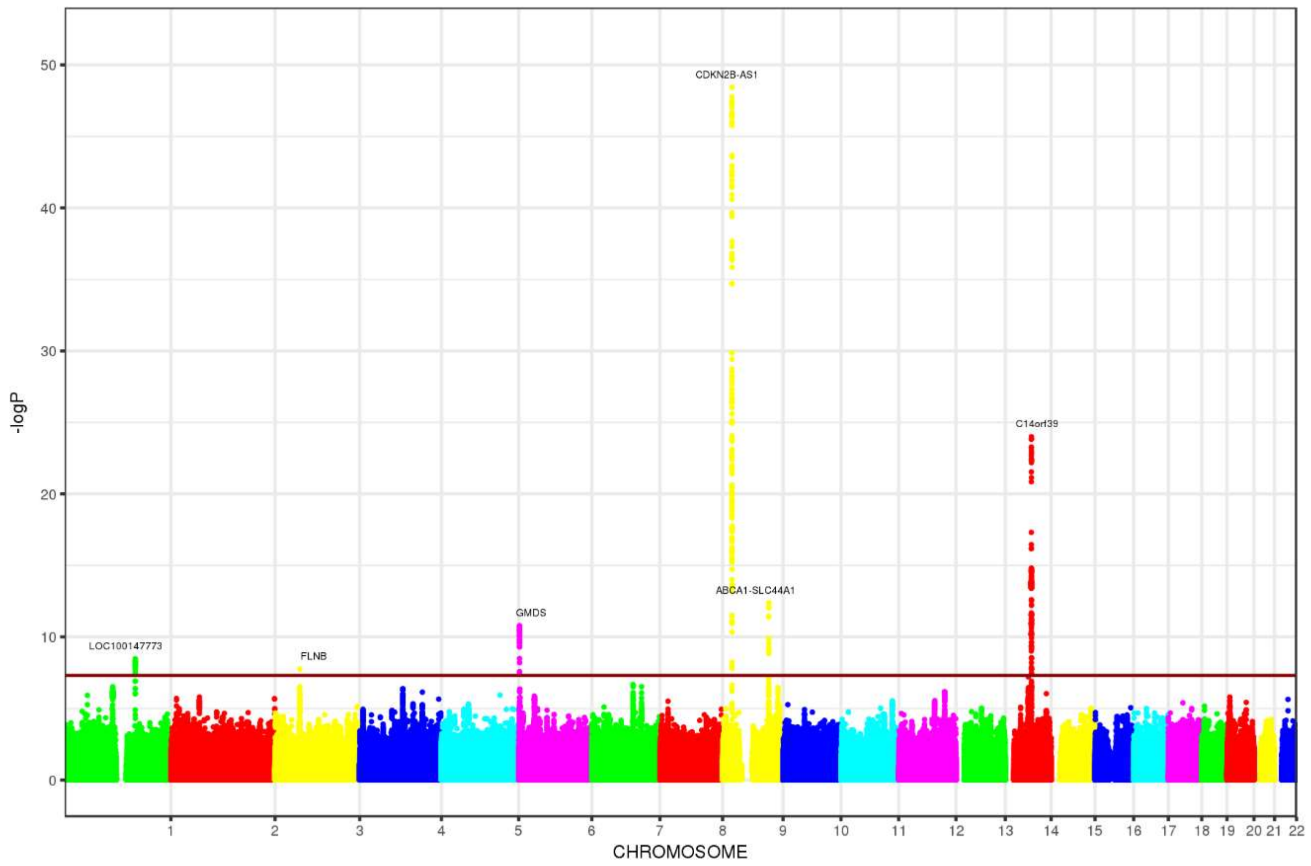

**F) HTG-stratified meta-analysis.**

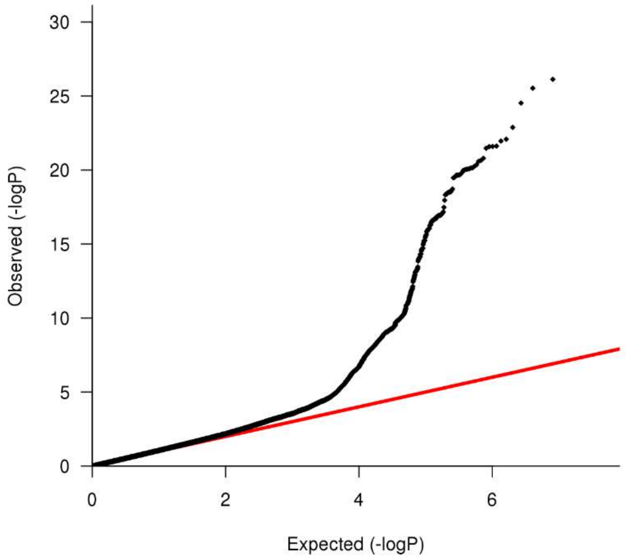


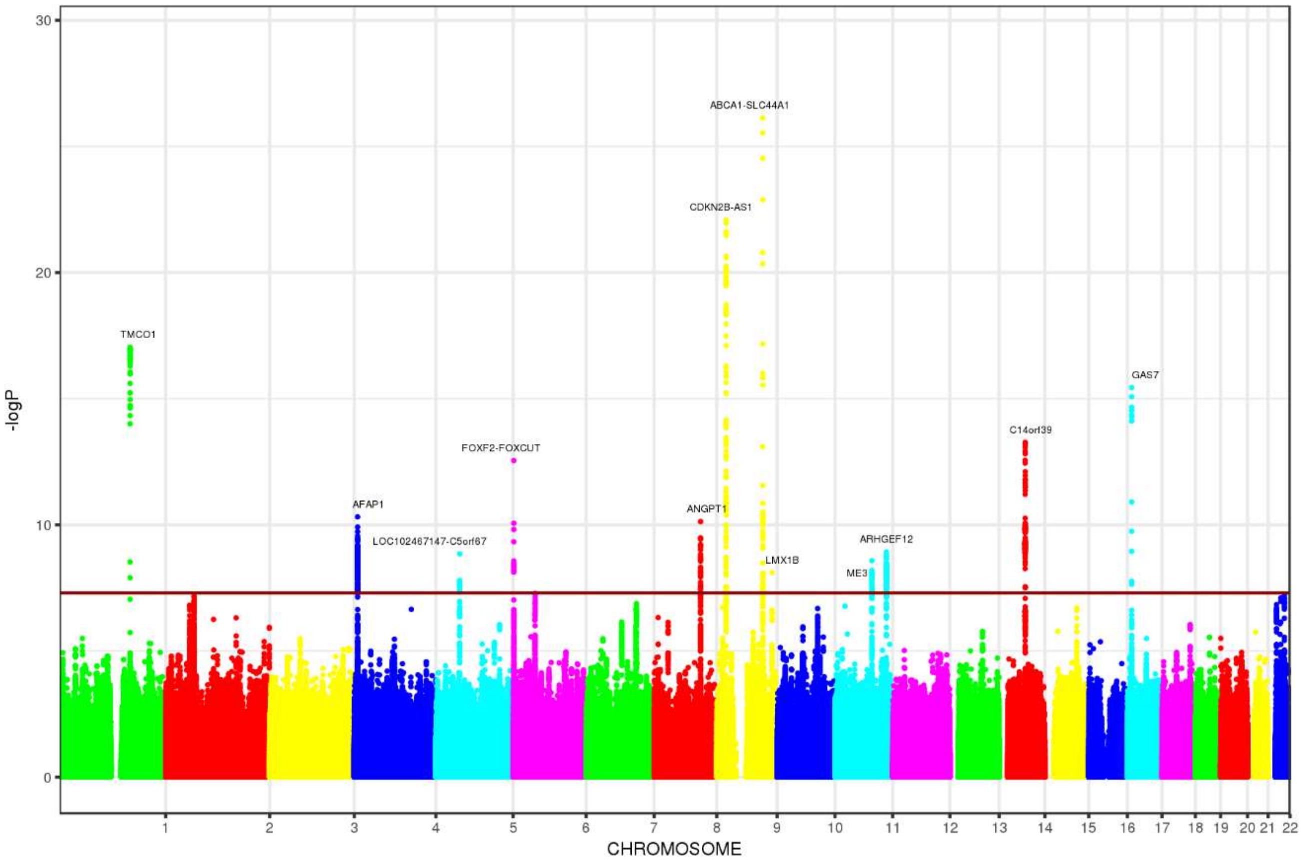

**Supplementary Figure 2. Correlation of SNP effect estimates between European POAG meta-analysis and each of the replication datasets.** The X-axis shows effect estimates in log(OR) scale for the European genome-wide significant SNPs. The Y-axis shows estimates for the same SNPs in UKBB self-reports (panel A), Asians (panel B), and Africans (panel C). The previously-identified risk loci are shown in red and the novel loci in blue. Horizontal grey bars on each dot represent the 95% confidence intervals (CIs) for the effect estimates in Europeans, and vertical grey bars shows the 95% CIs in the replication datasets. The blue line is the linear regression line best fitting the data.

**A) European POAG vs. UKBB self-reports.**

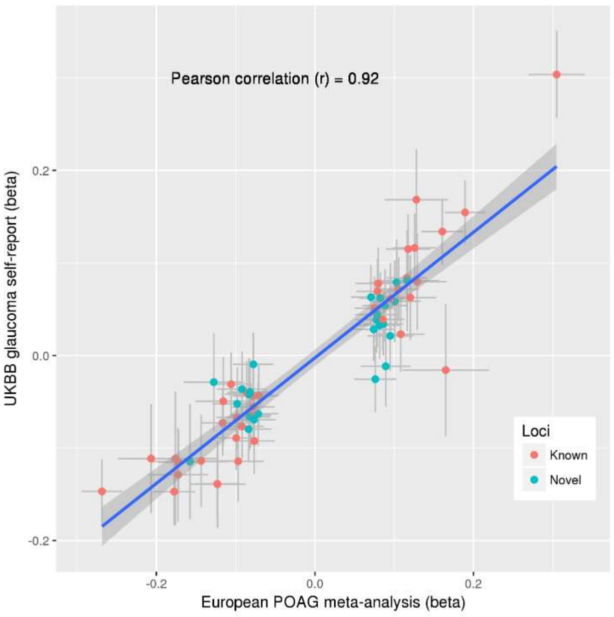


**B) European POAG vs. Asians.**

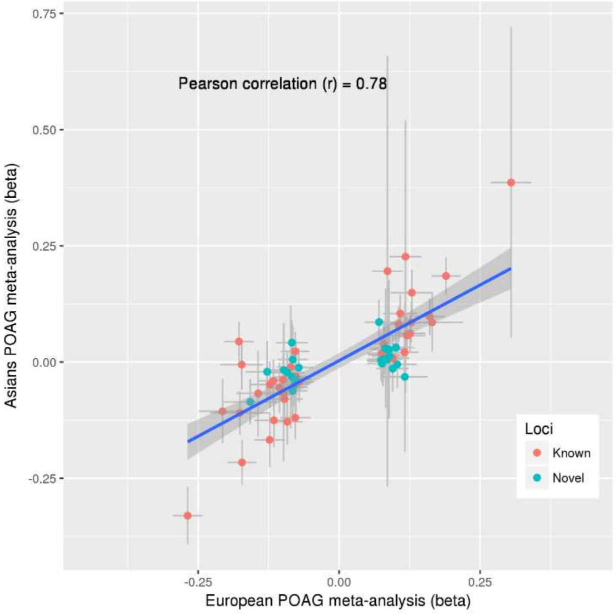


**C) European POAG vs. Africans.**

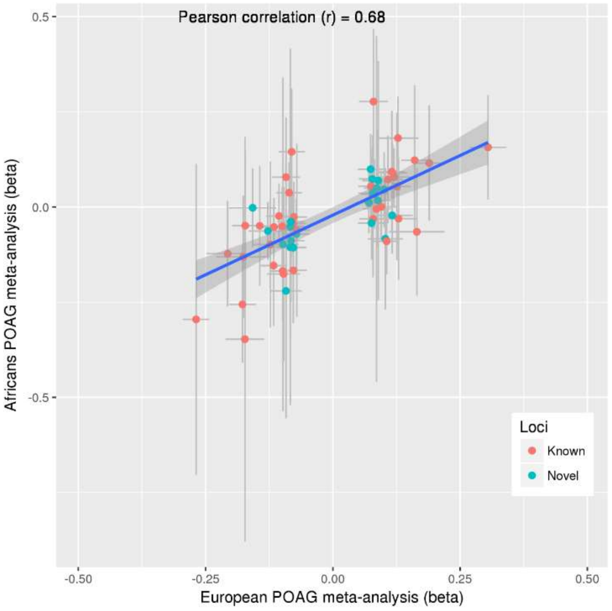

**Supplementary Figure 3. LocusZoom plots for the genome-wide significant SNPs from the cross-ancestry meta-analysis.** On each row, the plots show the same region in Europeans (first plot), Asians (second plot), and Africans (third plot). However, there are only two plots per row for the last four rows (X chromosome loci) because X chromosome data was not available for the African studies. The index SNPs are the most significant SNPs in the cross-ancestry meta-analysis while P-value of associations and LD structures are shown for each ancestry separately.

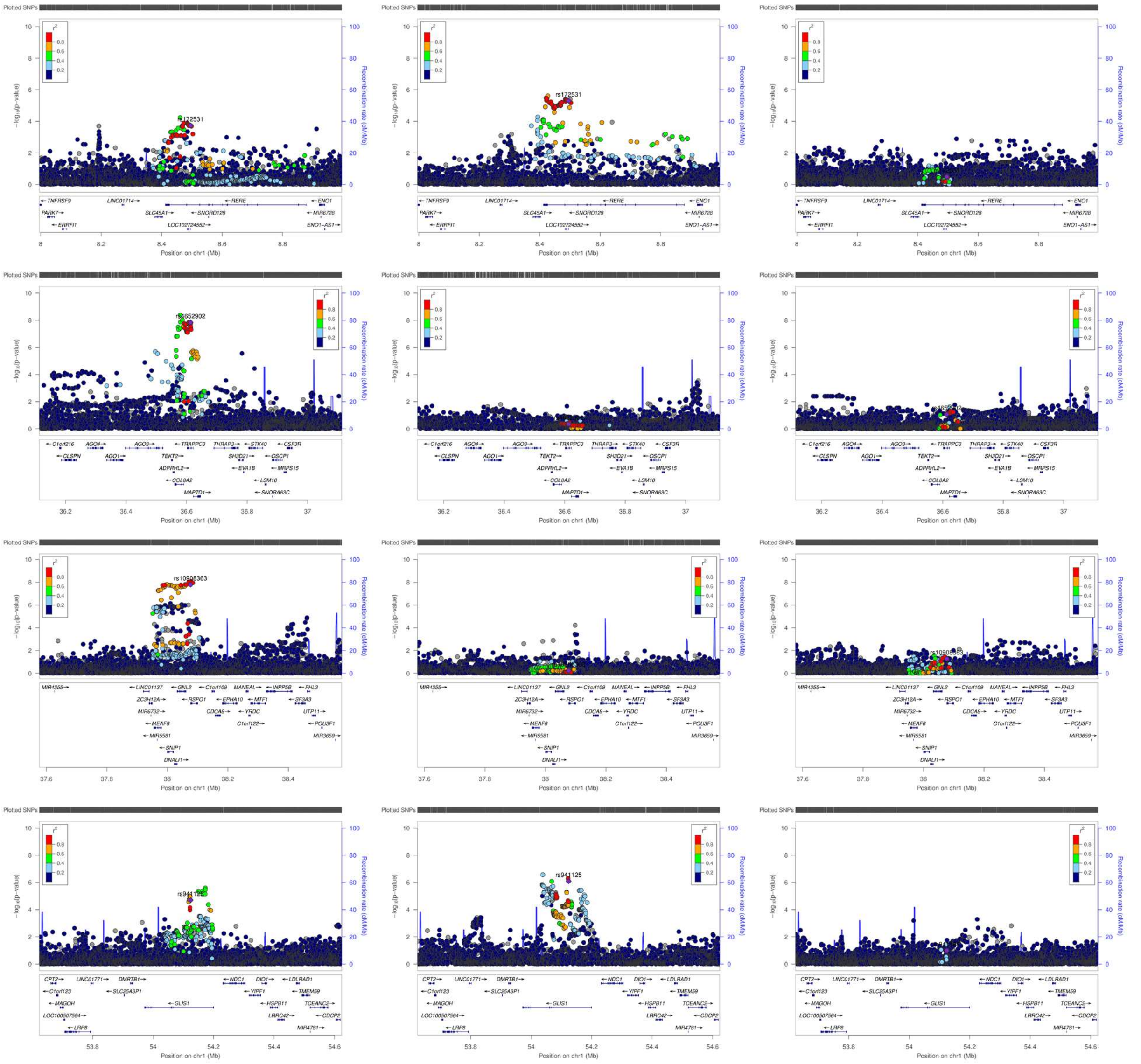

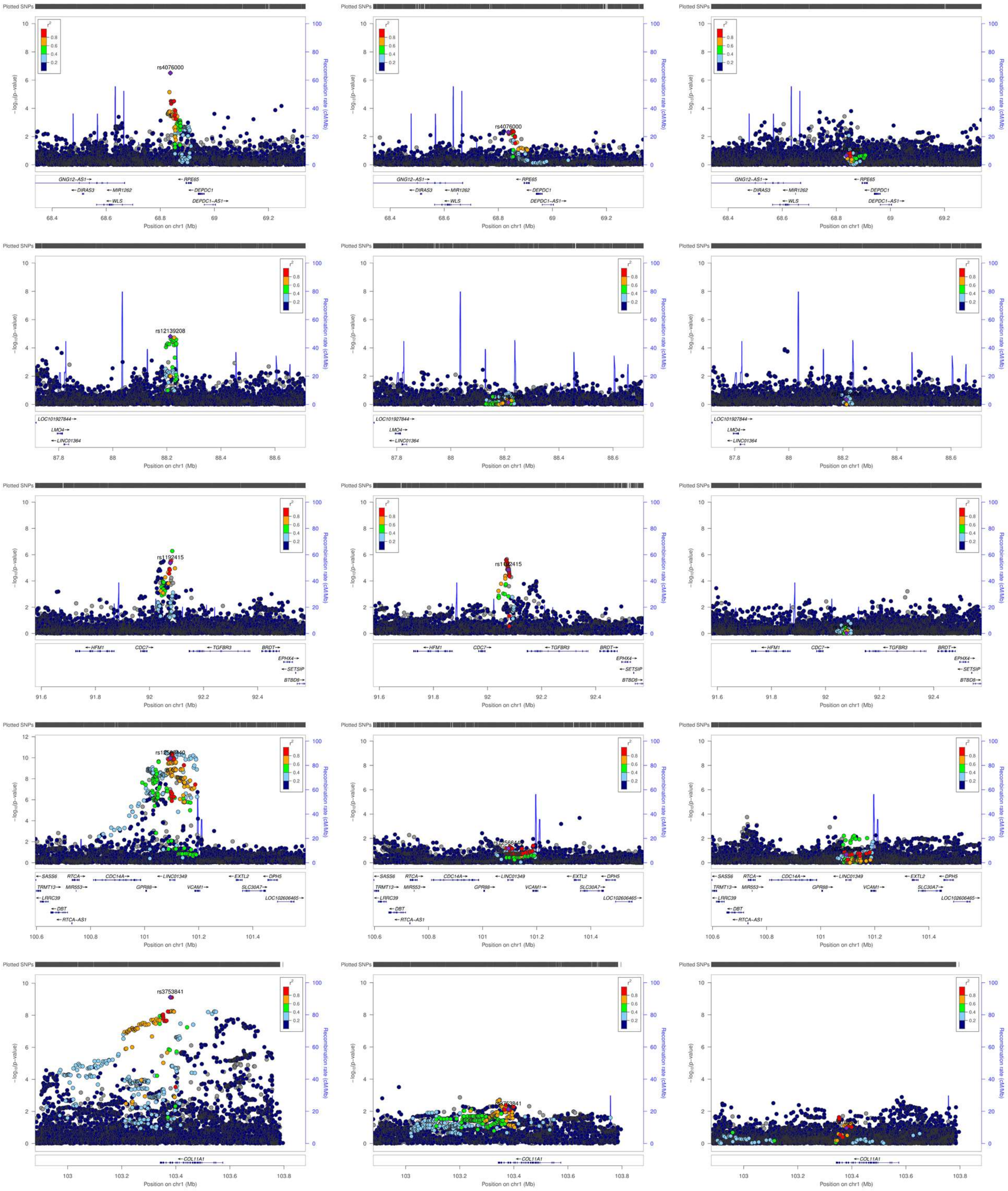
Plotted SNPs
$-\log_{10}(p\text{-value})$
Recombination rate (cM/Mb)
$r^2$
0.8
0.6
0.4
0.2
rs4076000
GNG12-AS1
DIRAS3
MIR1262
WLS
RPE65
DEPDC1
DEPDC1-AS1
Position on chr1 (Mb)
rs12139208
LOC101927844
LMO4
LINC01364
rs1192415
HFM1
CDC7
TGFBR3
BRDT
EPHX4
SETSIP
BTBD8
SASS6
RTCA
CDC14A
LINC01349
EXTL2
DPH5
TRMT13
MIR553
GPR88
VCAM1
SLC30A7
LOC102606465
LRRC39
DBT
RTCA-AS1
rs3753841
COL11A1

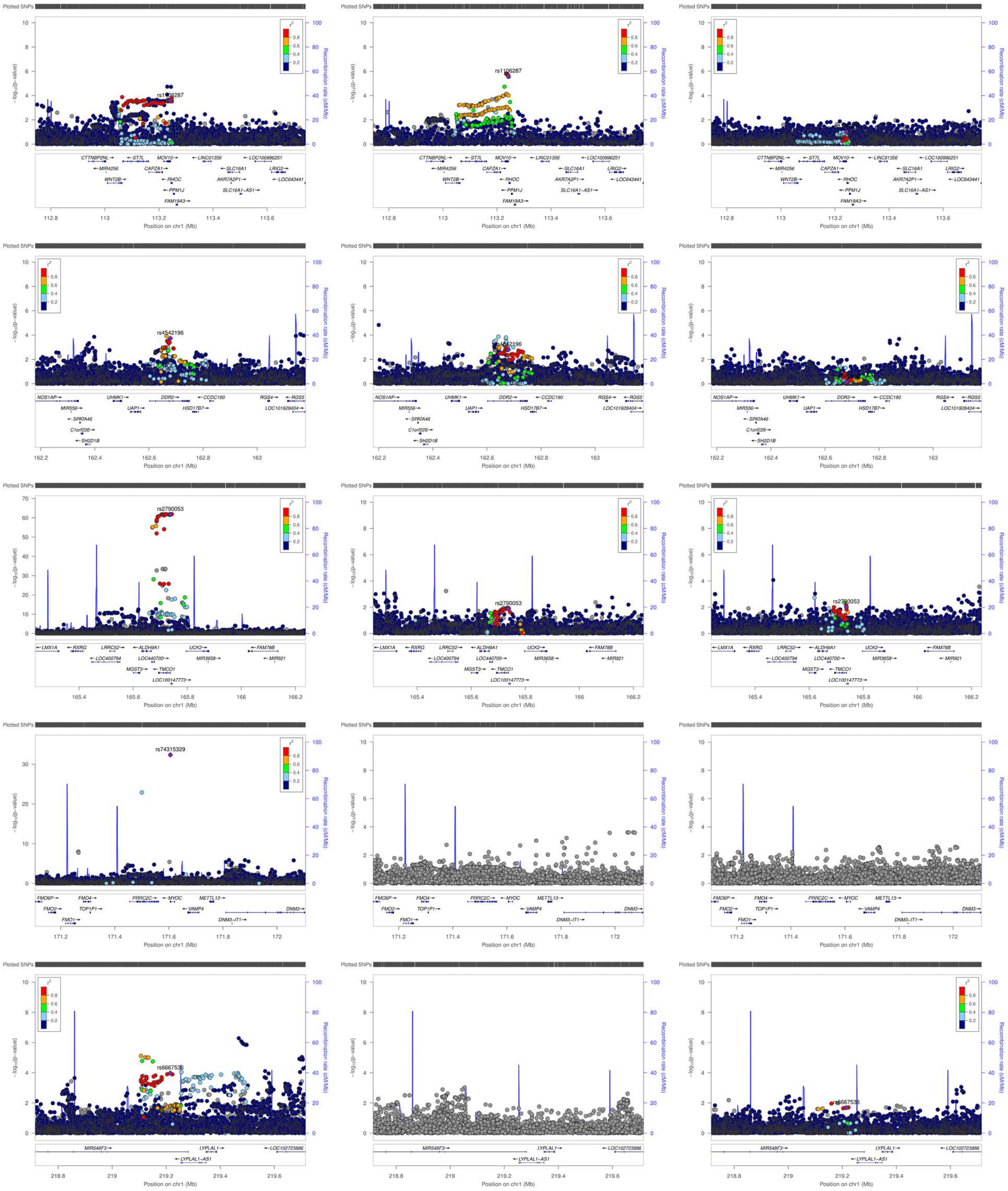

Plotted SNPs
rs1106287
rs4542196
rs2790053
rs74315329
rs6667536
-log10(p-value)
Recombination rate (cM/Mb)
r²
Position on chr1 (Mb)

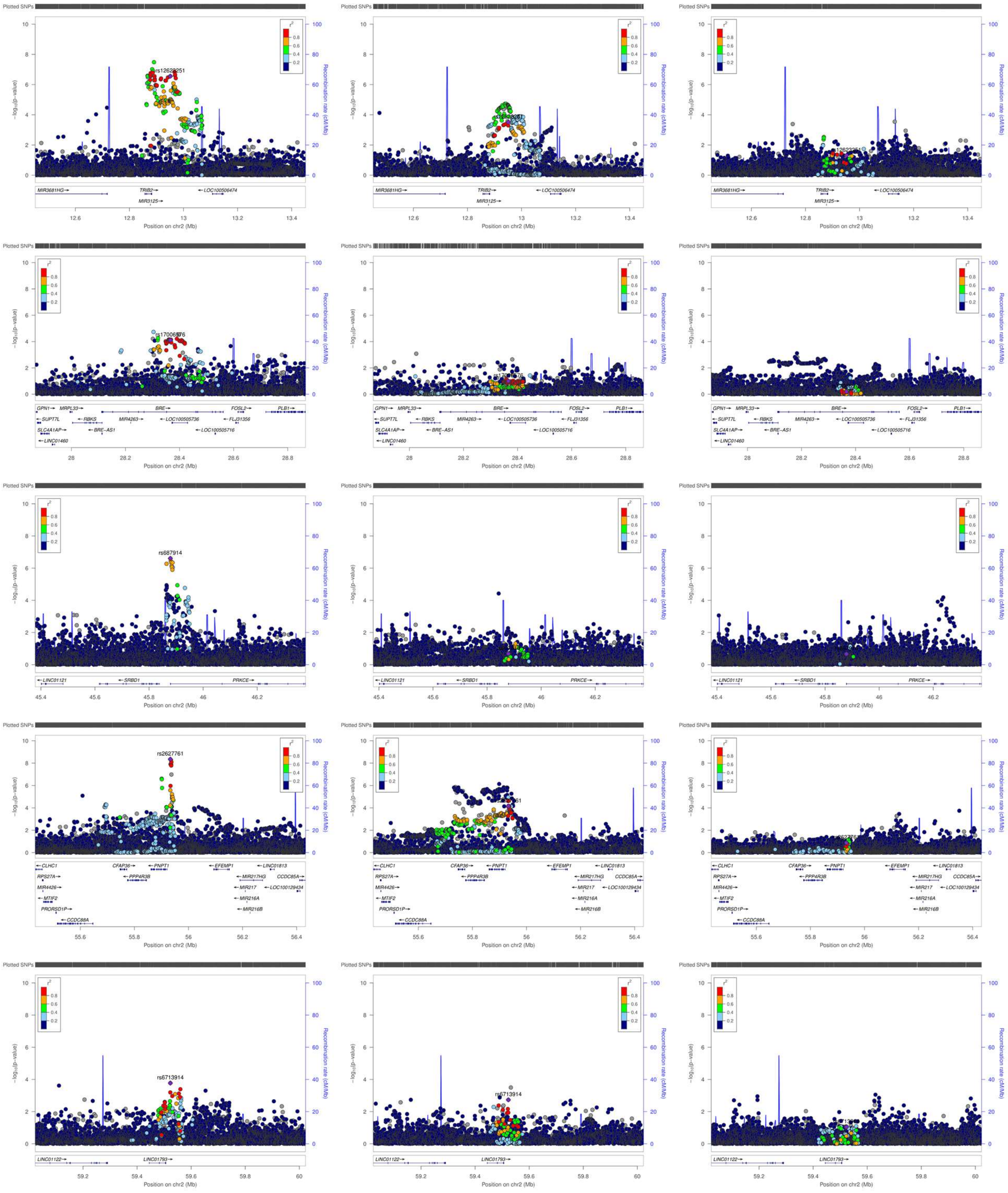

Plotted SNPs
-log10(p-value)
Recombination rate (cM/Mb)
r2
rs12621251
MIR3681HG
TRIB2
MIR3125
LOC100506474
Position on chr2 (Mb)
rs17006576
GPN1
MRPL33
BRE
FOSL2
PLB1
SUPT7L
RBKS
MIR4263
LOC100505736
FLJ31356
SLC4A1AP
BRE-AS1
LOC100505716
LINC01460
rs687914
LINC01121
SRBD1
PRKCE
rs2627761
CLHC1
CFAP36
PNPT1
EFEMP1
LINC01813
RPS27A
PPP4R3B
MIR217HG
CCDC85A
MIR4426
MIR217
LOC100129434
MTIF2
MIR216A
PRORSD1P
MIR216B
CCDC88A
rs6713914
LINC01122
LINC01793

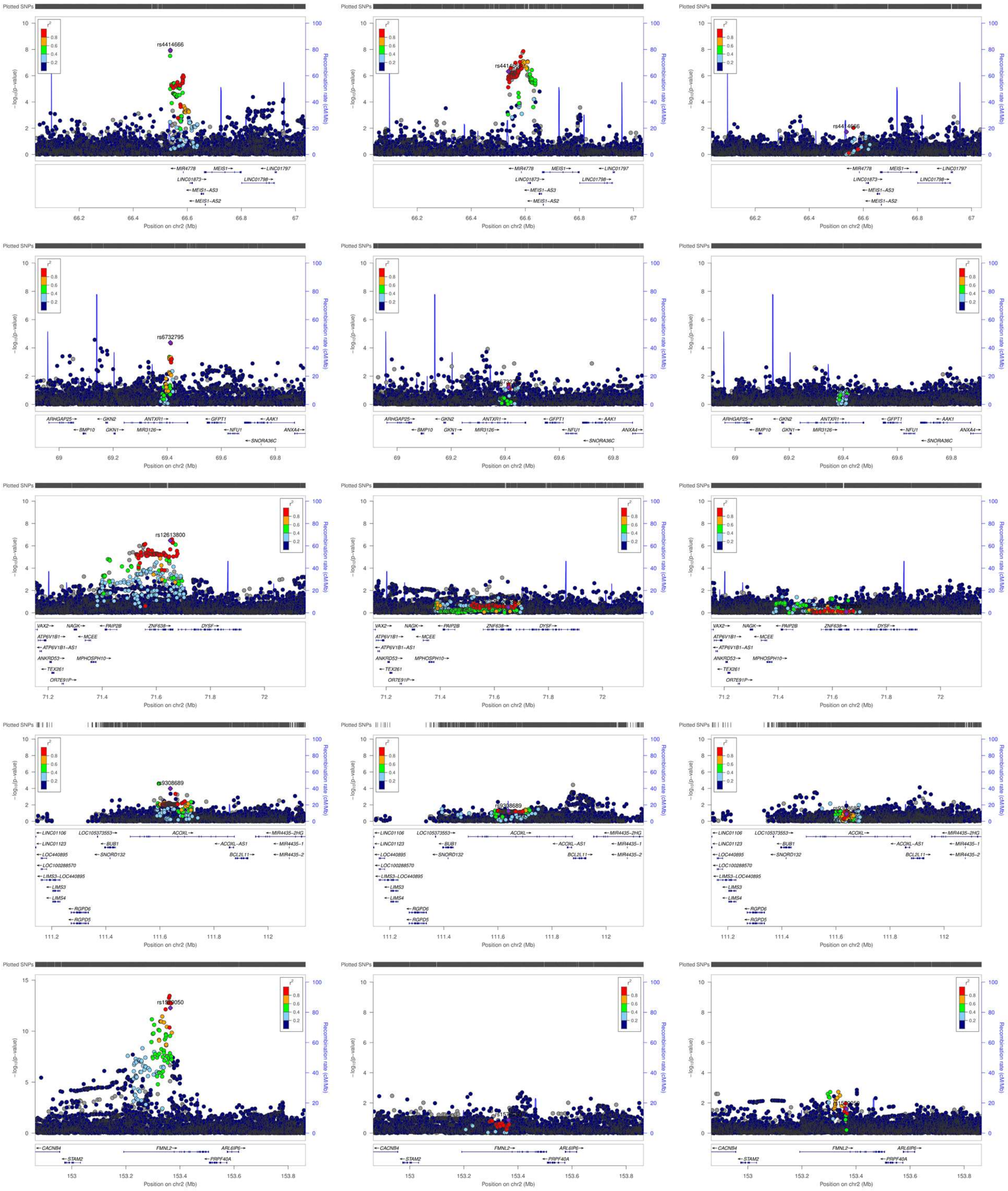

Plotted SNPs
−log10(p-value)
Recombination rate (cM/Mb)
r2
rs4414666
rs6732795
rs12613800
rs9308689
MIR4778
MEIS1
LINC01797
LINC01873
LINC01798
MEIS1-AS3
MEIS1-AS2
Position on chr2 (Mb)
ARHGAP25
GKN2
ANTXR1
GFPT1
AAK1
BMP10
GKN1
MIR3126
NFU1
ANXA4
SNORA36C
VAX2
NAGK
PAIP2B
ZNF638
DYSF
ATP6V1B1
MCEE
ATP6V1B1-AS1
ANKRD53
MPHOSPH10
TEX261
OR7E91P
LINC01106
LOC105373553
ACOXL
MIR4435-2HG
LINC01123
BUB1
ACOXL-AS1
MIR4435-1
LOC440895
SNORD132
BCL2L11
MIR4435-2
LOC100288570
LIMS3-LOC440895
LIMS3
LIMS4
RGPD6
RGPD5
CACNB4
FMNL2
ARL6IP6
STAM2
PRPF40A

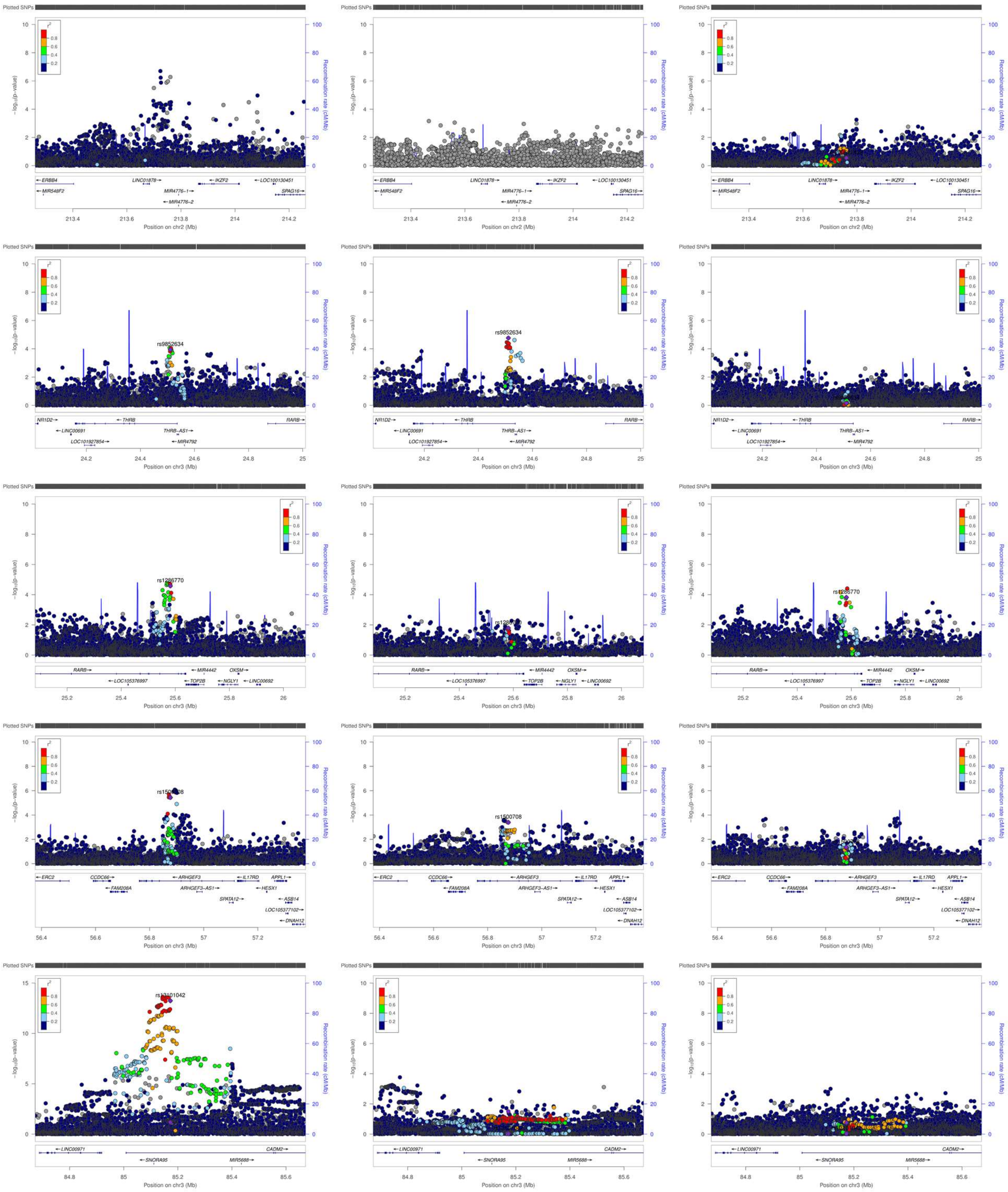

Plotted SNPs
Position on chr2 (Mb)
Position on chr3 (Mb)
Recombination rate (cM/Mb)
rs9852634
rs1286770
rs1500708

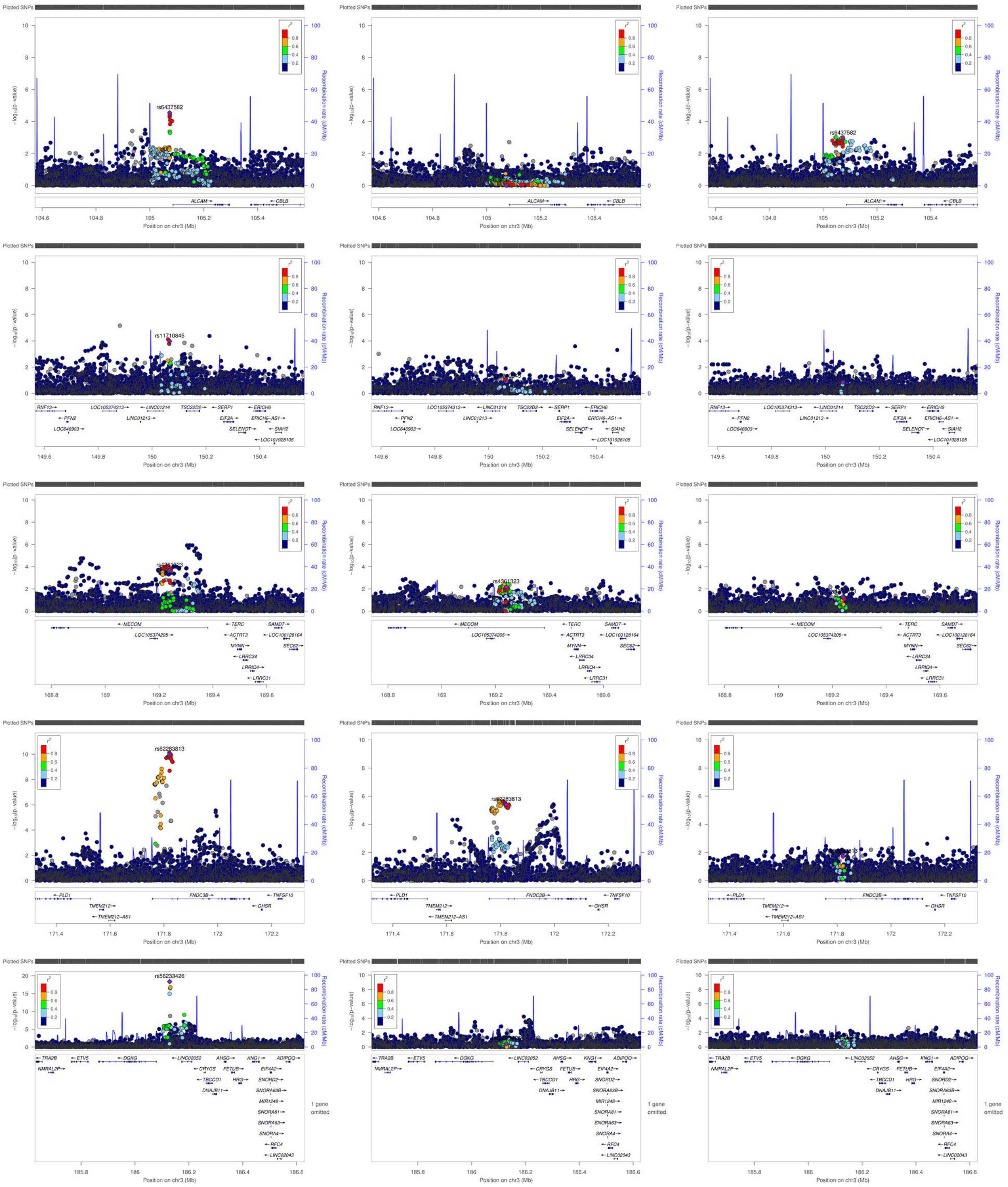

Plotted SNPs
rs6437582
rs11710845
rs62283813
rs56233426
ALCAM
CBLB
MECOM
FNDC3B
PLD1
TNFSF10
GHSR
DGKG
ETV5
TRA2B
AHSG
ADIPOQ
−log10(p-value)
Recombination rate (cM/Mb)
Position on chr3 (Mb)
1 gene omitted

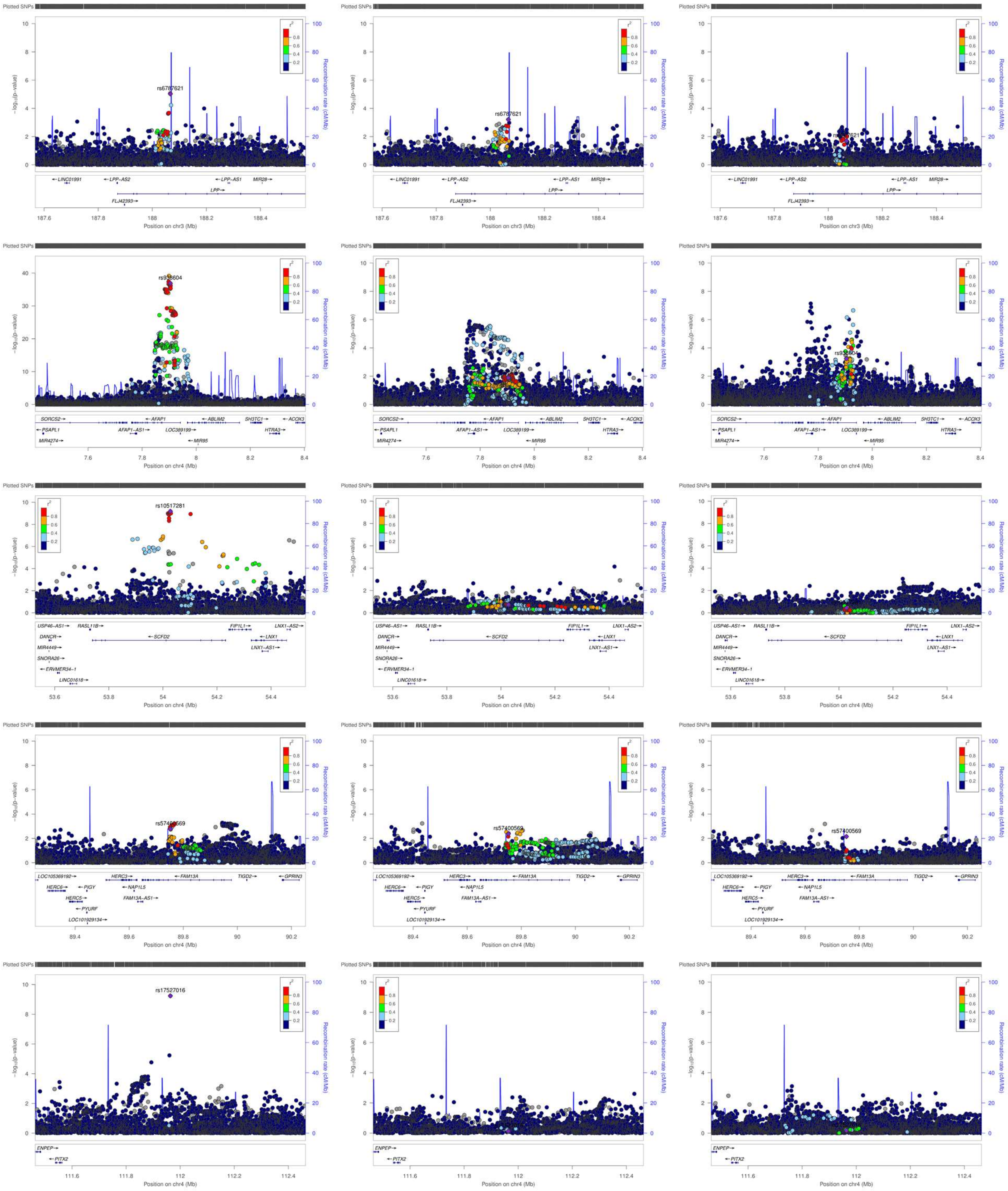

Plotted SNPs
rs6787621
rs938604
rs10517281
rs57400569
rs17527016
r²
0.8
0.6
0.4
0.2
−log₁₀(p-value)
Recombination rate (cM/Mb)
LINC01991
LPP-AS2
LPP-AS1
MIR28
LPP
FLJ42393
Position on chr3 (Mb)
SORCS2
PSAPL1
MIR4274
AFAP1
AFAP1-AS1
LOC389199
ABLIM2
MIR95
SH3TC1
HTRA3
ACOX3
Position on chr4 (Mb)
USP46-AS1
RASL11B
SCFD2
FIP1L1
LNX1-AS2
DANCR
LNX1
MIR4449
LNX1-AS1
SNORA26
ERVMER34-1
LINC01618
LOC105369192
HERC3
FAM13A
TIGD2
GPRIN3
HERC6
PIGY
NAP1L5
HERC5
FAM13A-AS1
PYURF
LOC101929134
ENPEP
PITX2

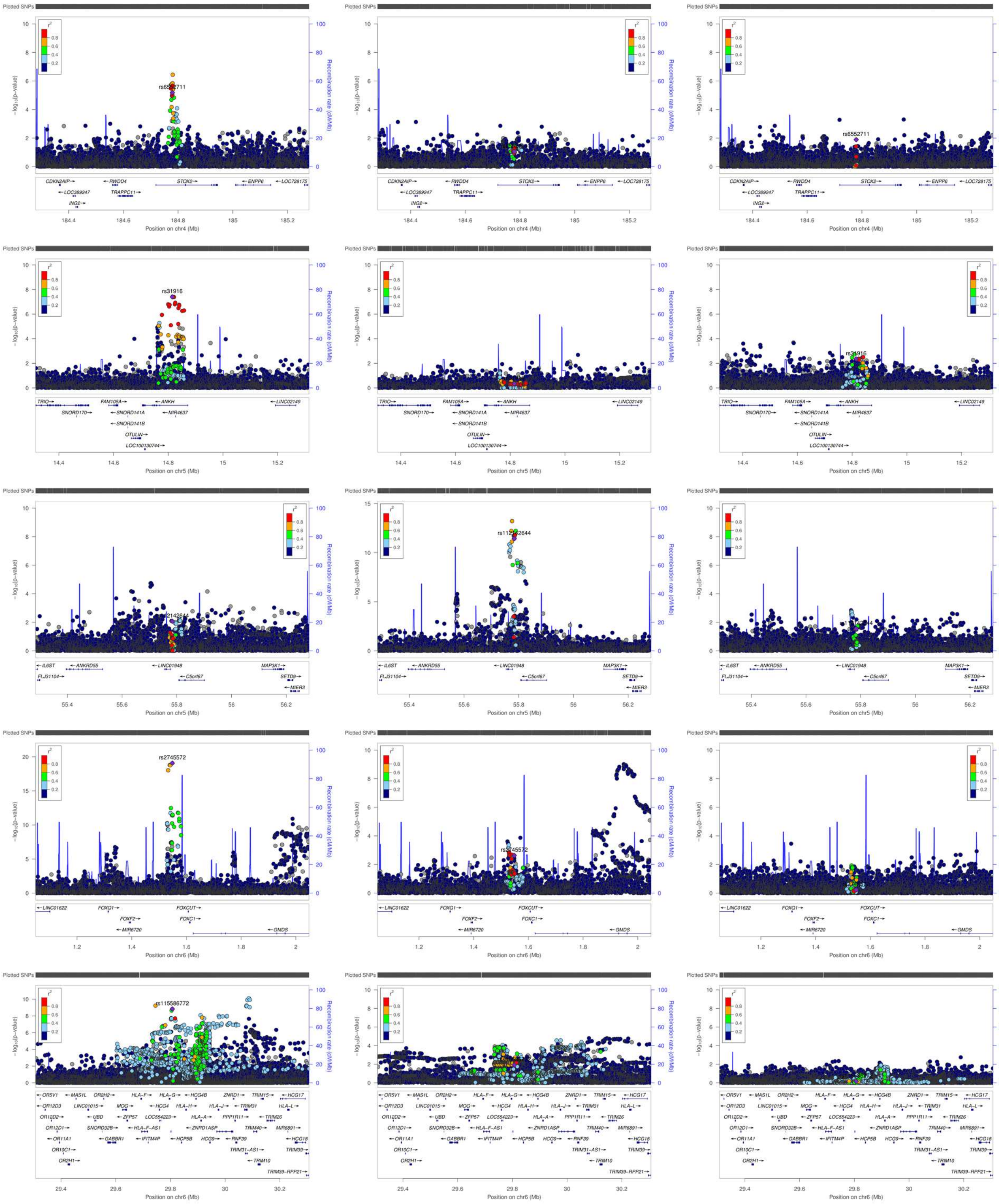
Plotted SNPs
rs6552711
rs31916
rs112142644
rs2745572
rs115586772
Position on chr4 (Mb)
Position on chr5 (Mb)
Position on chr6 (Mb)
Recombination rate (cM/Mb)

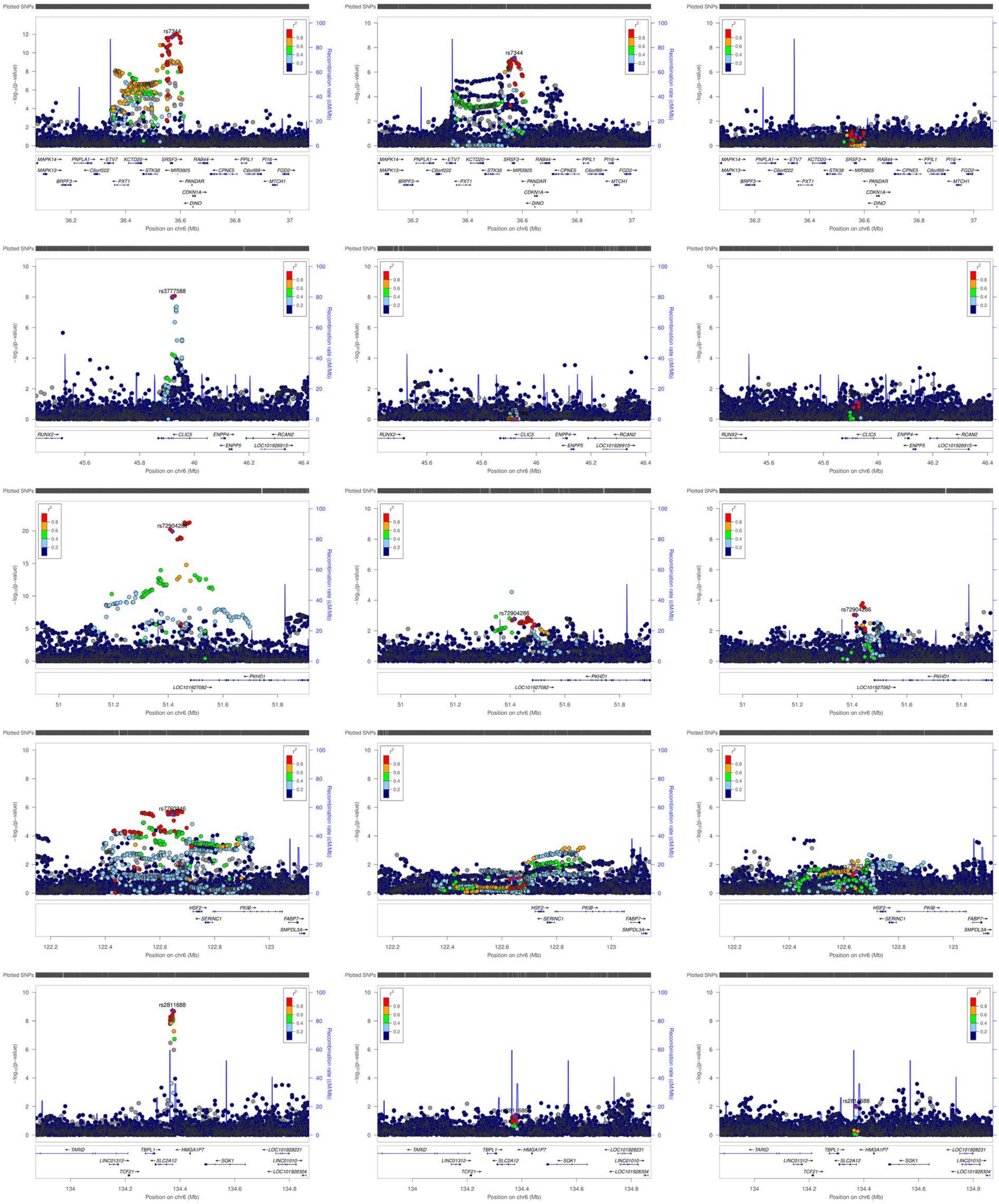

Plotted SNPs
-log10(p-value)
Recombination rate (cM/Mb)
r2
0.8
0.6
0.4
0.2
rs7344
MAPK14
MAPK13
BRPF3
PNPLA1
C6orf222
ETV7
PXT1
KCTD20
STK38
SRSF3
MIR3925
PANDAR
CDKN1A
DINO
RAB44
CPNE5
PPIL1
C6orf89
PI16
MTCH1
FGD2
Position on chr6 (Mb)
rs3777588
RUNX2
CLIC5
ENPP4
ENPP5
RCAN2
LOC101926915
rs72904286
PKHD1
LOC101927082
rs7760946
HSF2
SERINC1
PKIB
FABP7
SMPDL3A
rs2811688
TARID
LINC01312
TCF21
TBPL1
SLC2A12
HMGA1P7
SGK1
LOC101928231
LINC01010
LOC101928304

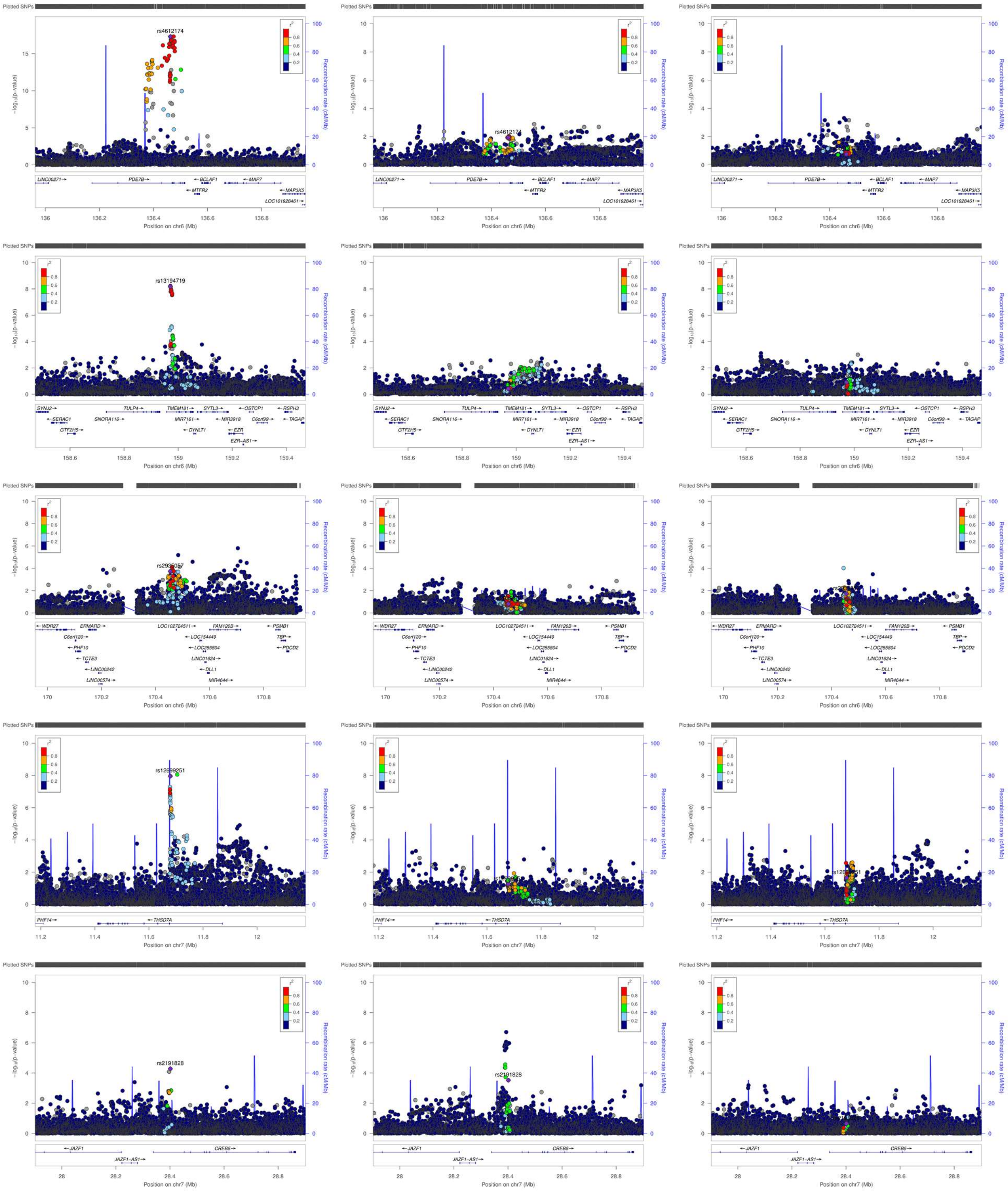
Plotted SNPs
−log10(p−value)
Recombination rate (cM/Mb)
r2
rs4612174
LINC00271→
PDE7B→
←BCLAF1
←MAP7
←MTFR2
←MAP3K5
LOC101928461→
Position on chr6 (Mb)
rs13194719
SYNJ2→
TULP4→
TMEM181→
SYTL3→
←OSTCP1
←RSPH3
←SERAC1
SNORA116→
MIR7161→
←MIR3918
C6orf99→
←TAGAP
GTF2H5→
←DYNLT1
←EZR
EZR−AS1→
←WDR27
ERMARD→
LOC102724511→
FAM120B→
←PSMB1
C6orf120→
←LOC154449
TBP→
←PHF10
←LOC285804
←PDCD2
←TCTE3
LINC01624→
←LINC00242
←DLL1
LINC00574→
MIR4644→
rs12699251
PHF14→
←THSD7A
Position on chr7 (Mb)
rs2191828
←JAZF1
CREB5→
JAZF1−AS1→

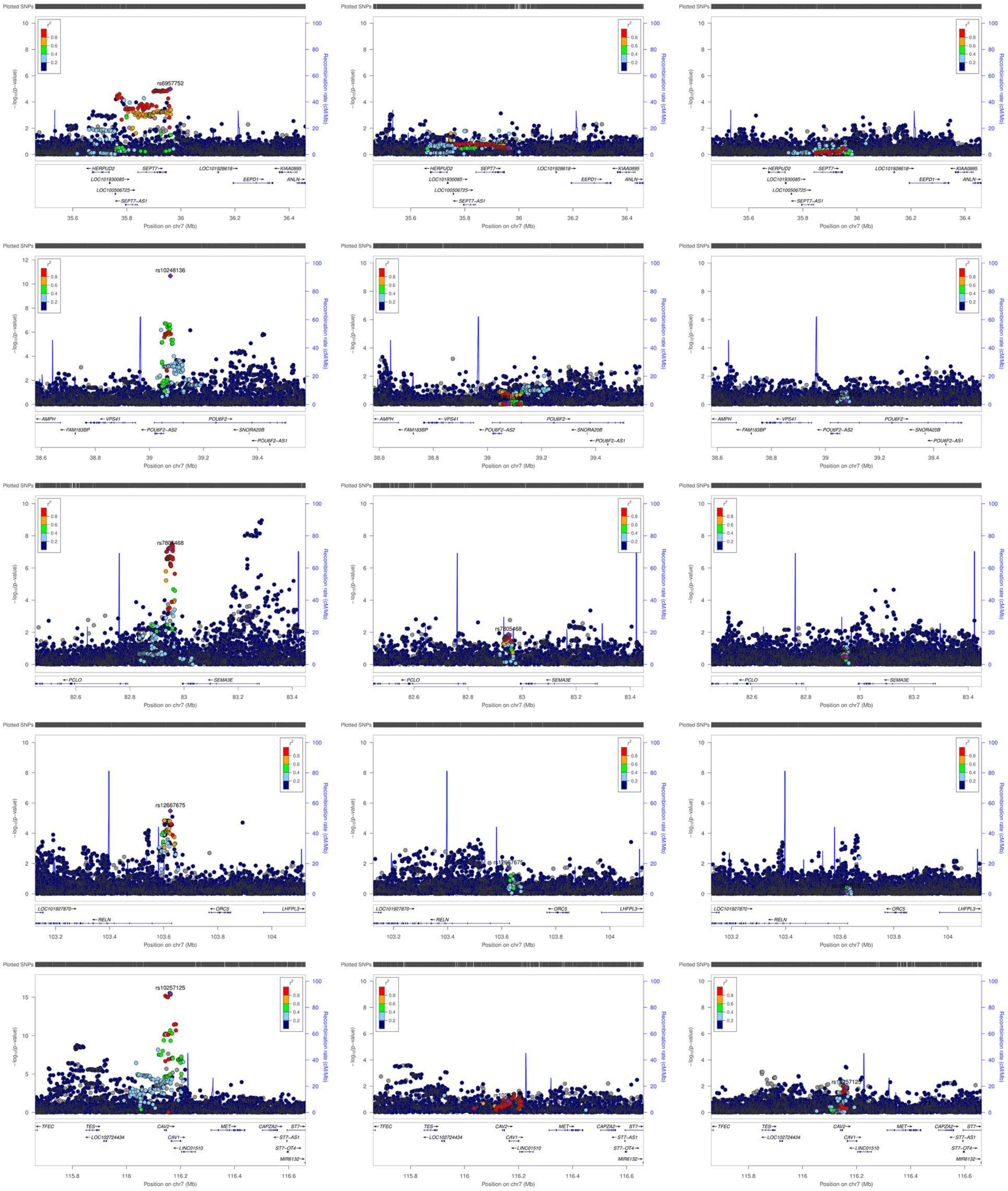
Plotted SNPs
rs6957752
rs10248136
rs7805468
rs12667675
rs10257125
HERPUD2
SEPT7
LOC101928618
KIAA0895
EEPD1
ANLN
LOC101930085
LOC100506725
SEPT7-AS1
AMPH
VPS41
POU6F2
FAM183BP
POU6F2-AS2
SNORA20B
POU6F2-AS1
PCLO
SEMA3E
LOC101927870
RELN
ORC5
LHFPL3
TFEC
TES
CAV2
MET
CAPZA2
ST7
LOC102724434
CAV1
ST7-AS1
LINC01510
ST7-OT4
MIR6132
Position on chr7 (Mb)
Recombination rate (cM/Mb)

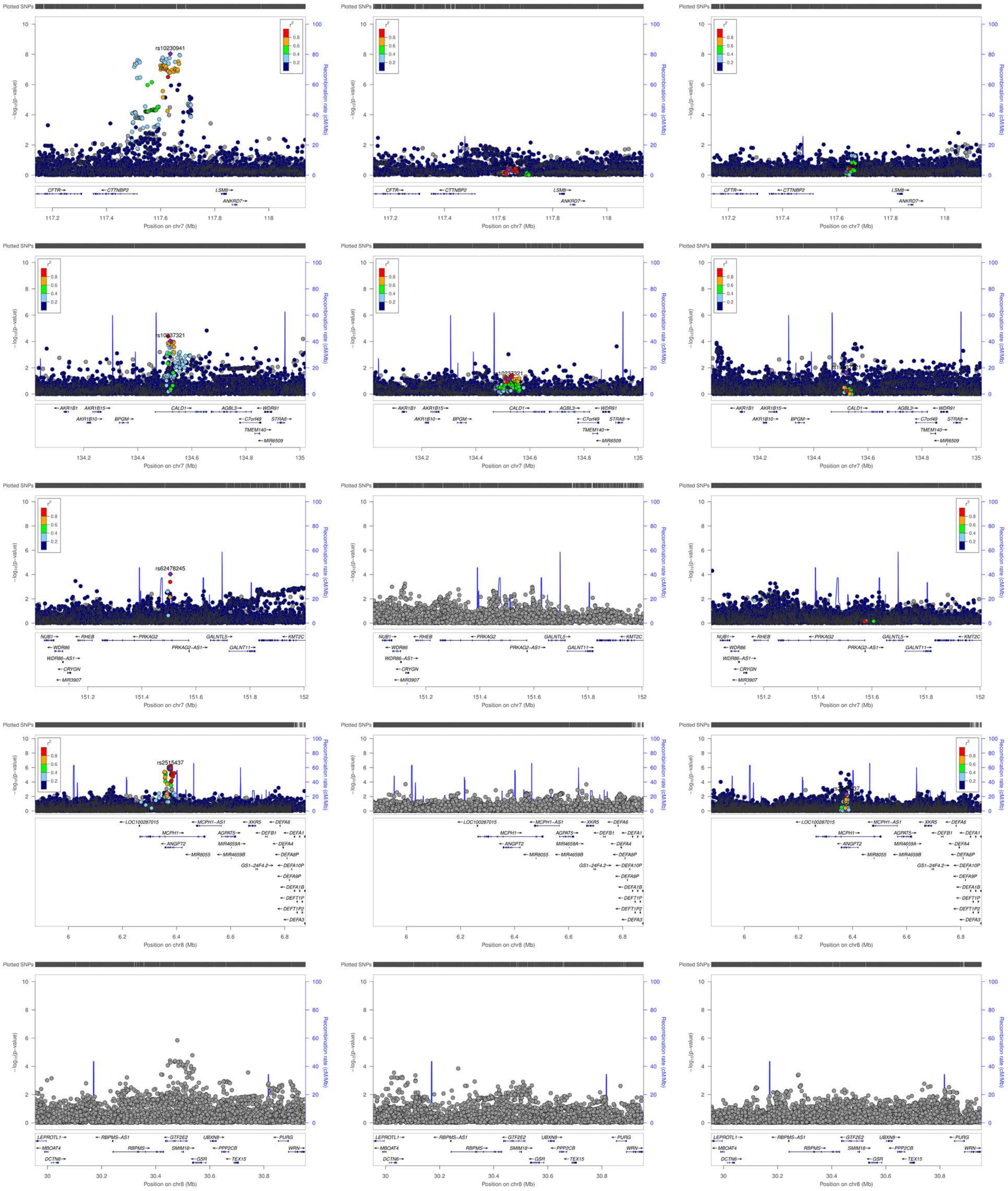

Plotted SNPs
−log10(p-value)
Recombination rate (cM/Mb)
r2
rs10230941
CFTR
CTTNBP2
LSM8
ANKRD7
Position on chr7 (Mb)
rs10237321
AKR1B1
AKR1B15
AKR1B10
BPGM
CALD1
AGBL3
WDR91
C7orf49
STRA8
TMEM140
MIR6509
rs62478245
NUB1
RHEB
WDR86
WDR86−AS1
CRYGN
MIR3907
PRKAG2
PRKAG2−AS1
GALNTL5
GALNT11
KMT2C
rs2515437
LOC100287015
MCPH1−AS1
MCPH1
ANGPT2
AGPAT5
MIR4659A
MIR4659B
MIR8055
XKR5
DEFA6
DEFB1
DEFA1
DEFA4
DEFA8P
DEFA10P
GS1−24F4.2
DEFA9P
DEFA1B
DEFT1P
DEFT1P2
DEFA3
Position on chr8 (Mb)
LEPROTL1
MBOAT4
DCTN6
RBPMS−AS1
RBPMS
GTF2E2
SMIM18
GSR
UBXN8
PPP2CB
TEX15
PURG
WRN

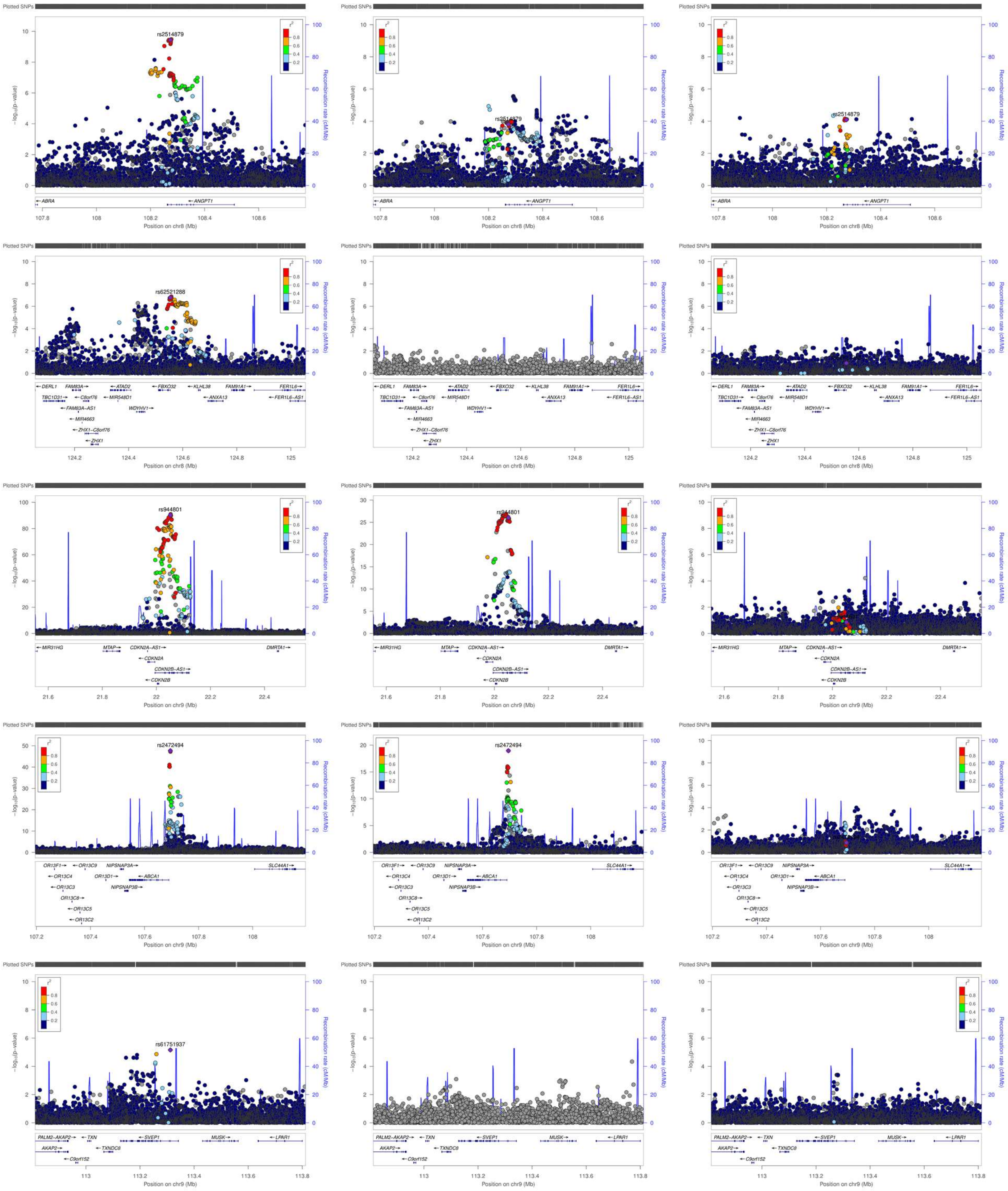

Plotted SNPs
rs2514879
rs62521288
rs944801
rs2472494
rs61751937
−log10(p-value)
Recombination rate (cM/Mb)
r2
Position on chr8 (Mb)
Position on chr9 (Mb)
ABRA
ANGPT1
DERL1
FAM83A
ATAD2
FBXO32
KLHL38
FAM91A1
FER1L6
TBC1D31
C8orf76
MIR548D1
ANXA13
FER1L6-AS1
FAM83A-AS1
WDYHV1
MIR4663
ZHX1-C8orf76
ZHX1
MIR31HG
MTAP
CDKN2A-AS1
DMRTA1
CDKN2A
CDKN2B-AS1
CDKN2B
OR13F1
OR13C9
NIPSNAP3A
SLC44A1
OR13C4
OR13D1
ABCA1
OR13C3
NIPSNAP3B
OR13C8
OR13C5
OR13C2
PALM2-AKAP2
TXN
SVEP1
MUSK
LPAR1
AKAP2
TXNDC8
C9orf152

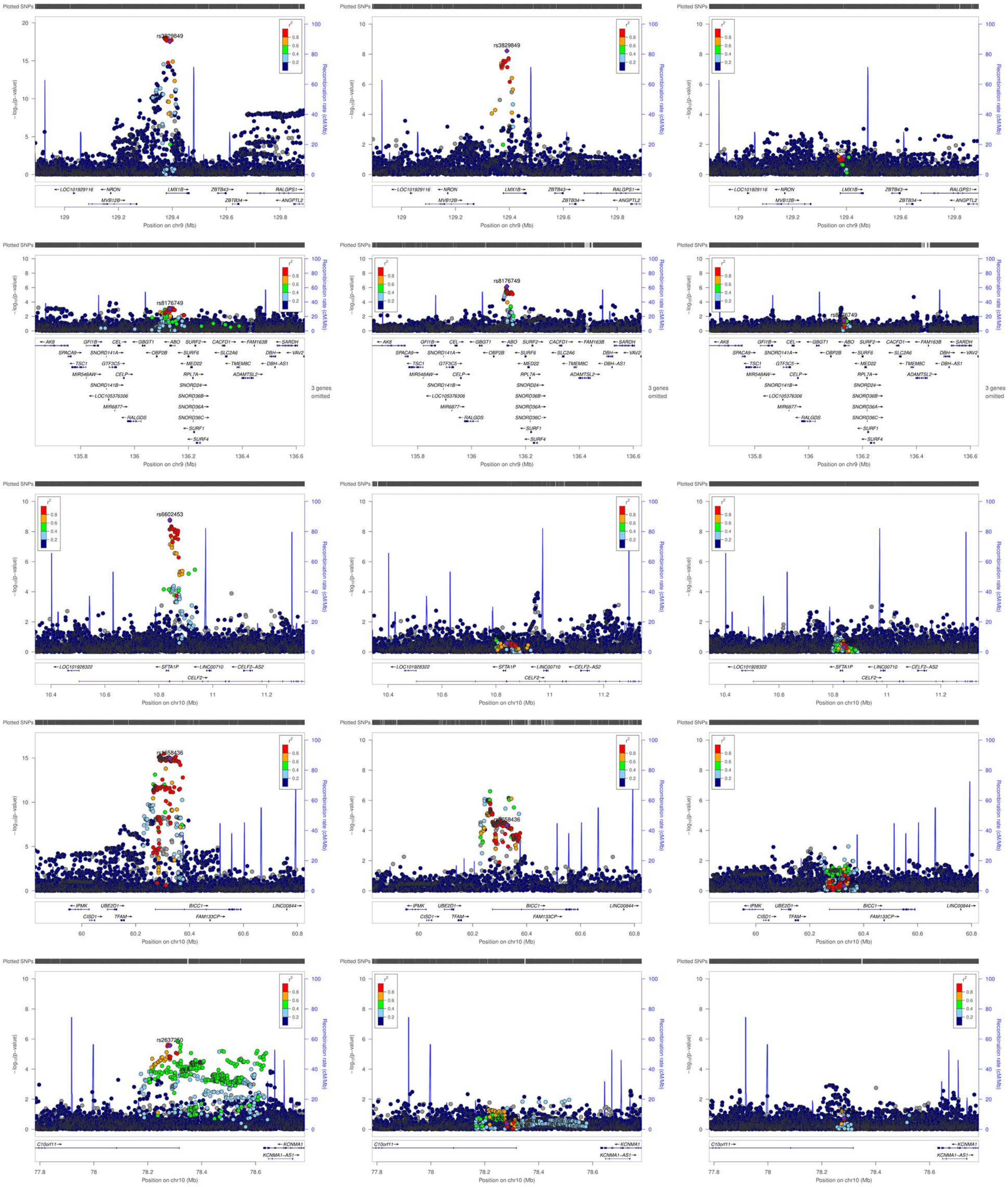
Plotted SNPs
rs3829849
rs8176749
rs6602453
rs2637250
Position on chr9 (Mb)
Position on chr10 (Mb)
Recombination rate (cM/Mb)
3 genes omitted
LMX1B
ZBTB43
ZBTB34
RALGPS1
ANGPTL2
MVB12B
NRON
ABO
SFTA1P
CELF2
BICC1
FAM133CP
KCNMA1
KCNMA1-AS1
C10orf11

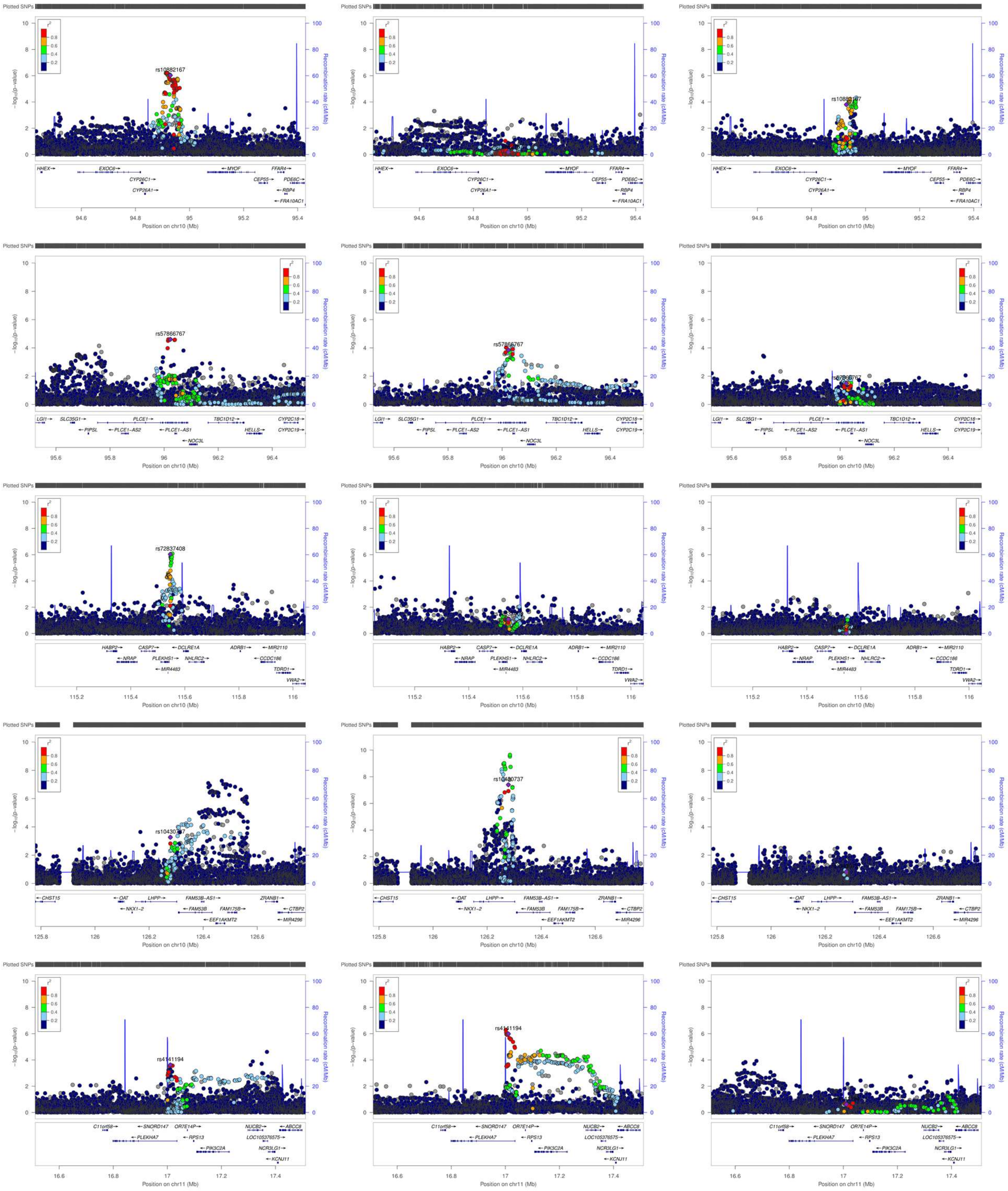

Plotted SNPs
-log10(p-value)
Recombination rate (cM/Mb)
r2
0.8
0.6
0.4
0.2
rs10882167
HHEX
EXOC6
CYP26C1
CYP26A1
MYOF
CEP55
FFAR4
PDE6C
RBP4
FRA10AC1
Position on chr10 (Mb)
rs57866767
LGI1
SLC35G1
PIPSL
PLCE1
PLCE1-AS2
PLCE1-AS1
NOC3L
TBC1D12
HELLS
CYP2C18
CYP2C19
rs72837408
HABP2
NRAP
CASP7
PLEKHS1
MIR4483
DCLRE1A
NHLRC2
ADRB1
MIR2110
CCDC186
TDRD1
VWA2
rs10430737
CHST15
OAT
NKX1-2
LHPP
FAM53B-AS1
FAM53B
FAM175B
EEF1AKMT2
ZRANB1
CTBP2
MIR4296
rs4141194
C11orf58
SNORD147
PLEKHA7
OR7E14P
RPS13
PIK3C2A
NUCB2
LOC105376575
NCR3LG1
KCNJ11
ABCC8
Position on chr11 (Mb)

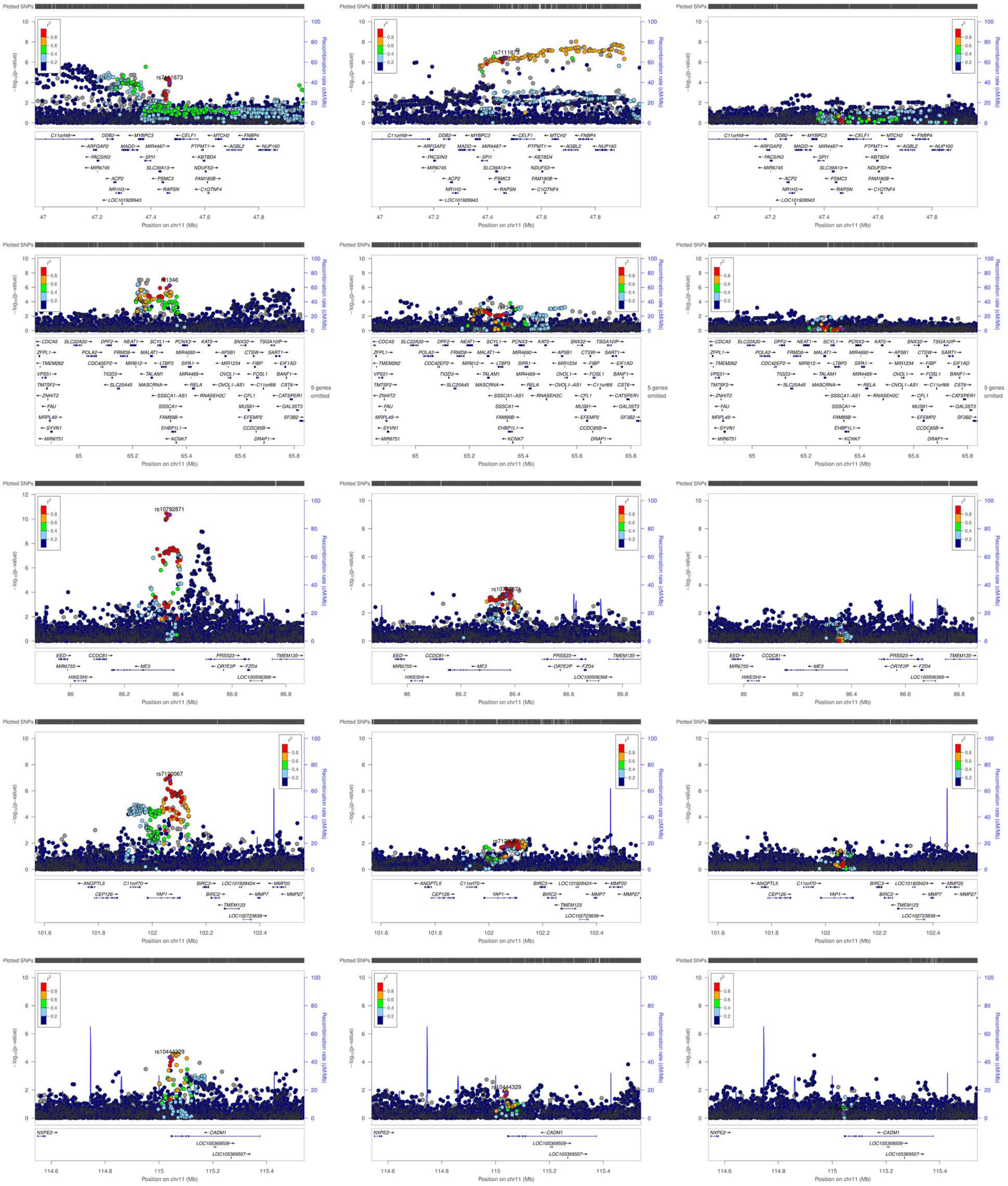

Plotted SNPs
−log10(p-value)
Recombination rate (cM/Mb)
Position on chr11 (Mb)
rs7111873
rs10792871
rs7120067
rs10444329
5 genes omitted

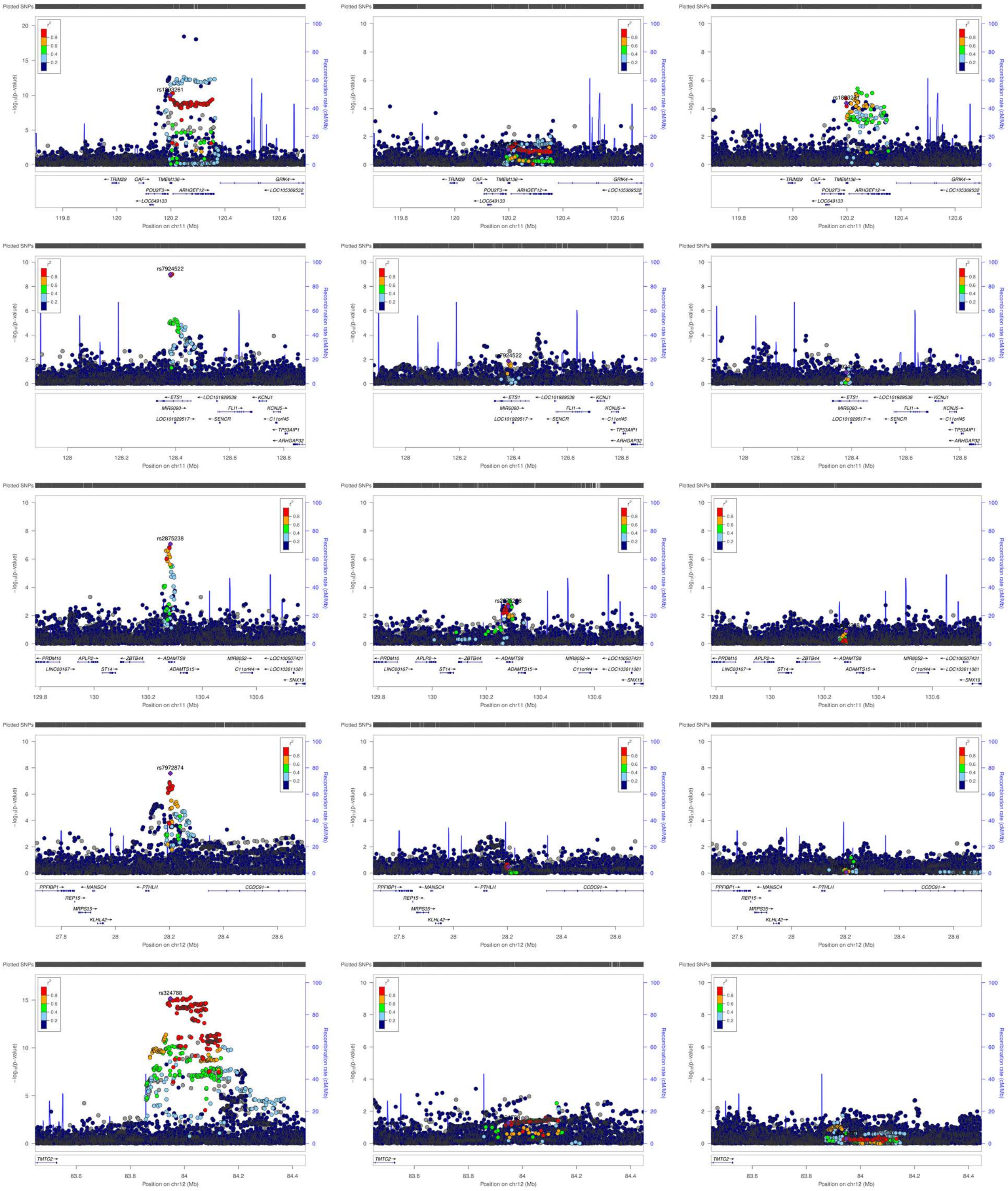

Plotted SNPs
-log10(p-value)
Recombination rate (cM/Mb)
r²
rs1893261
rs7924522
rs2875238
rs7972874
rs324788
Position on chr11 (Mb)
Position on chr12 (Mb)
TRIM29
OAF
TMEM136
POU2F3
ARHGEF12
GRIK4
LOC105369532
LOC649133
ETS1
LOC101929538
KCNJ1
MIR6090
FLI1
KCNJ5
LOC101929517
SENCR
C11orf45
TP53AIP1
ARHGAP32
PRDM10
APLP2
ZBTB44
ADAMTS8
MIR8052
LOC100507431
LINC00167
ST14
ADAMTS15
C11orf44
LOC103611081
SNX19
PPFIBP1
MANSC4
PTHLH
CCDC91
REP15
MRPS35
KLHL42
TMTC2

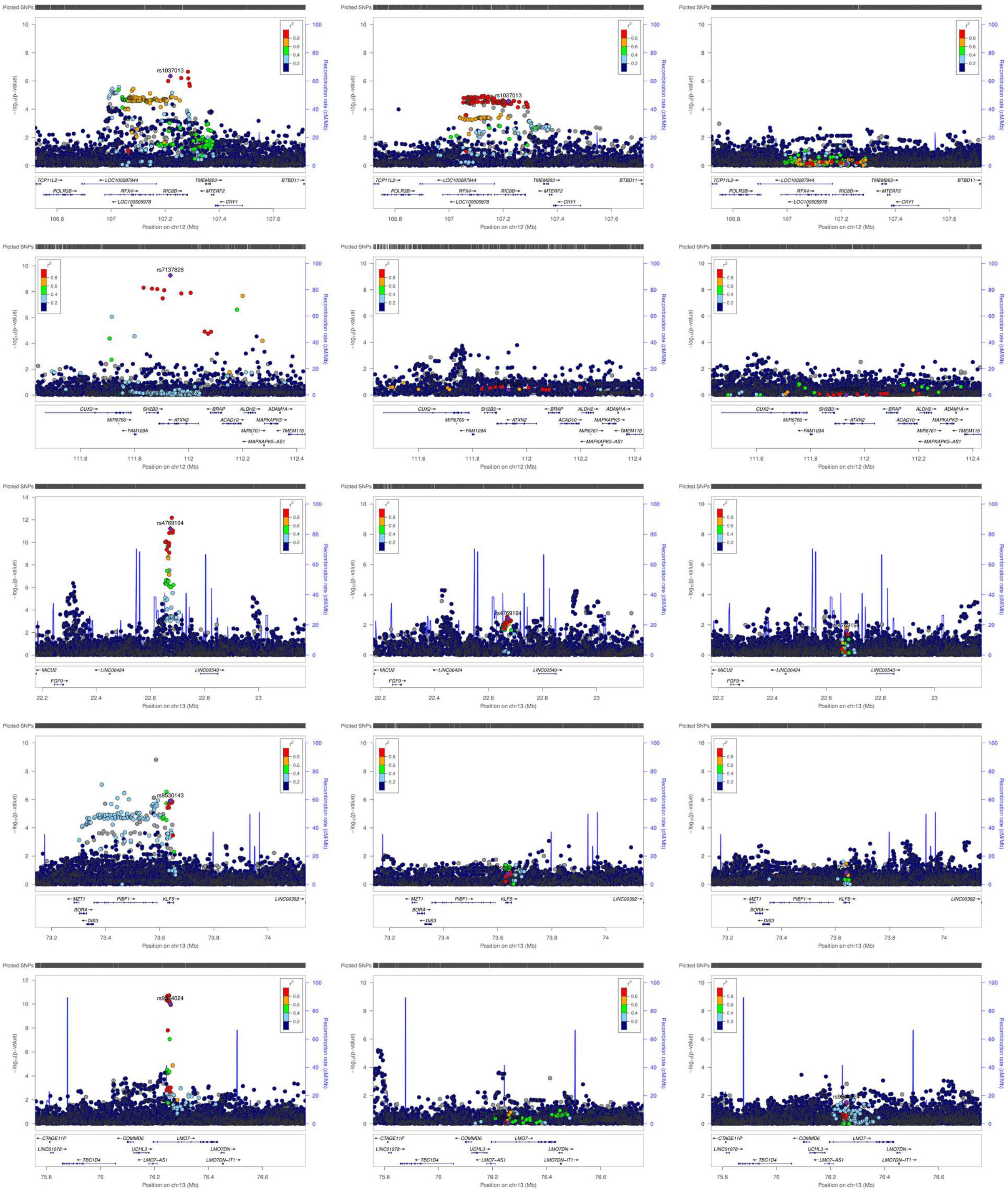

Plotted SNPs
$-\log_{10}$(p-value)
Recombination rate (cM/Mb)
$r^2$
0.8
0.6
0.4
0.2
rs1037013
TCP11L2→
←LOC100287944
TMEM263→
BTBD11→
POLR3B→
RFX4→
RIC8B→
←MTERF2
←LOC100505978
←CRY1
Position on chr12 (Mb)
rs7137828
CUX2→
SH2B3→
←BRAP
ALDH2→
ADAM1A→
MIR6760→
←ATXN2
ACAD10→
MAPKAPK5→
←FAM109A
MIR6761→
←TMEM116
←MAPKAPK5−AS1
rs4769194
←MICU2
←LINC00424
LINC00540→
FGF9→
Position on chr13 (Mb)
rs9530143
MZT1
PIBF1→
KLF5→
LINC00392→
BORA→
←DIS3
←CTAGE11P
←COMMD6
LMO7→
LINC01078→
UCHL3→
LMO7DN→
←TBC1D4
←LMO7−AS1
LMO7DN−IT1→

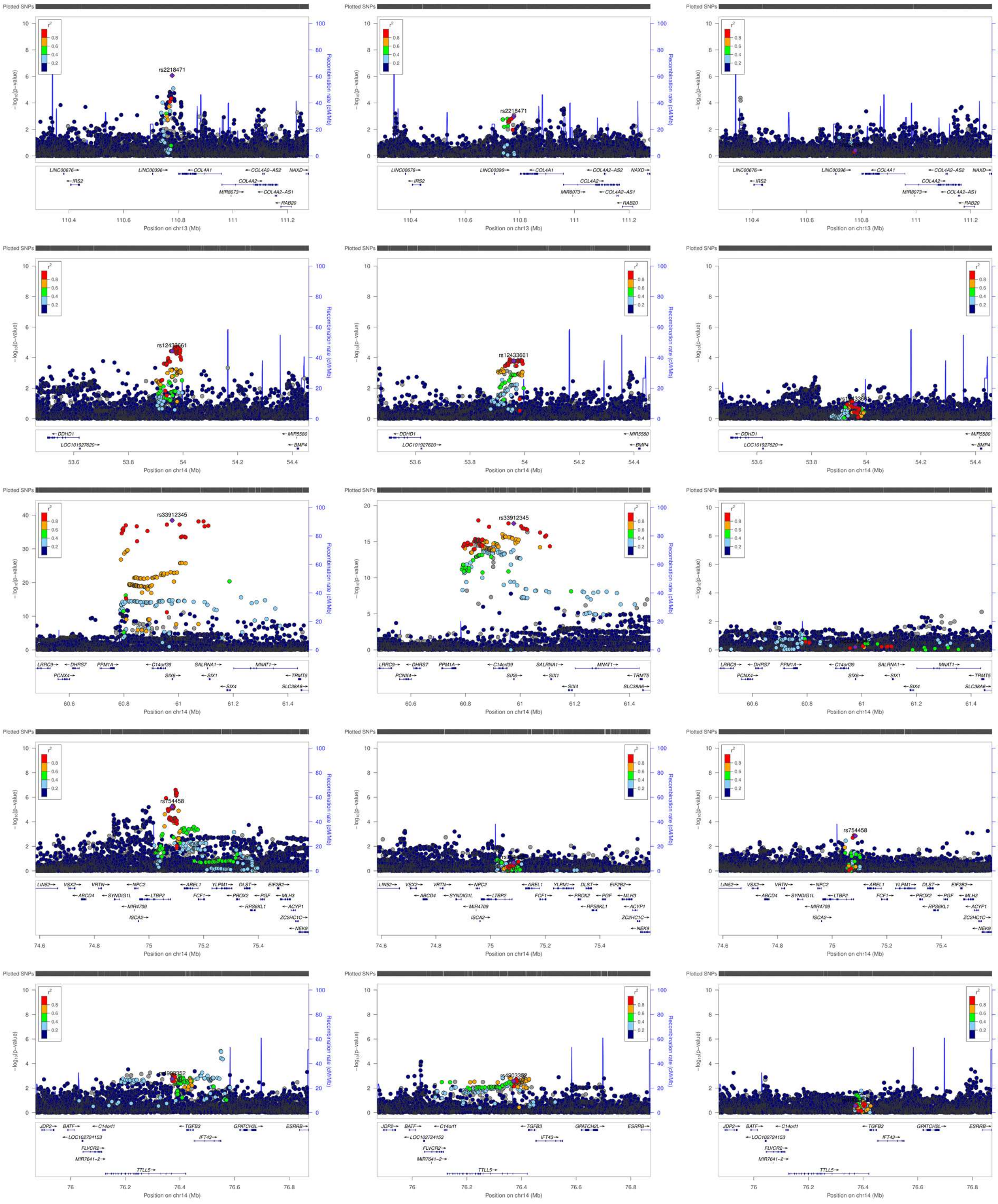

Plotted SNPs
rs2218471
rs12433661
rs33912345
rs754458
-log10(p-value)
Recombination rate (cM/Mb)
Position on chr13 (Mb)
Position on chr14 (Mb)

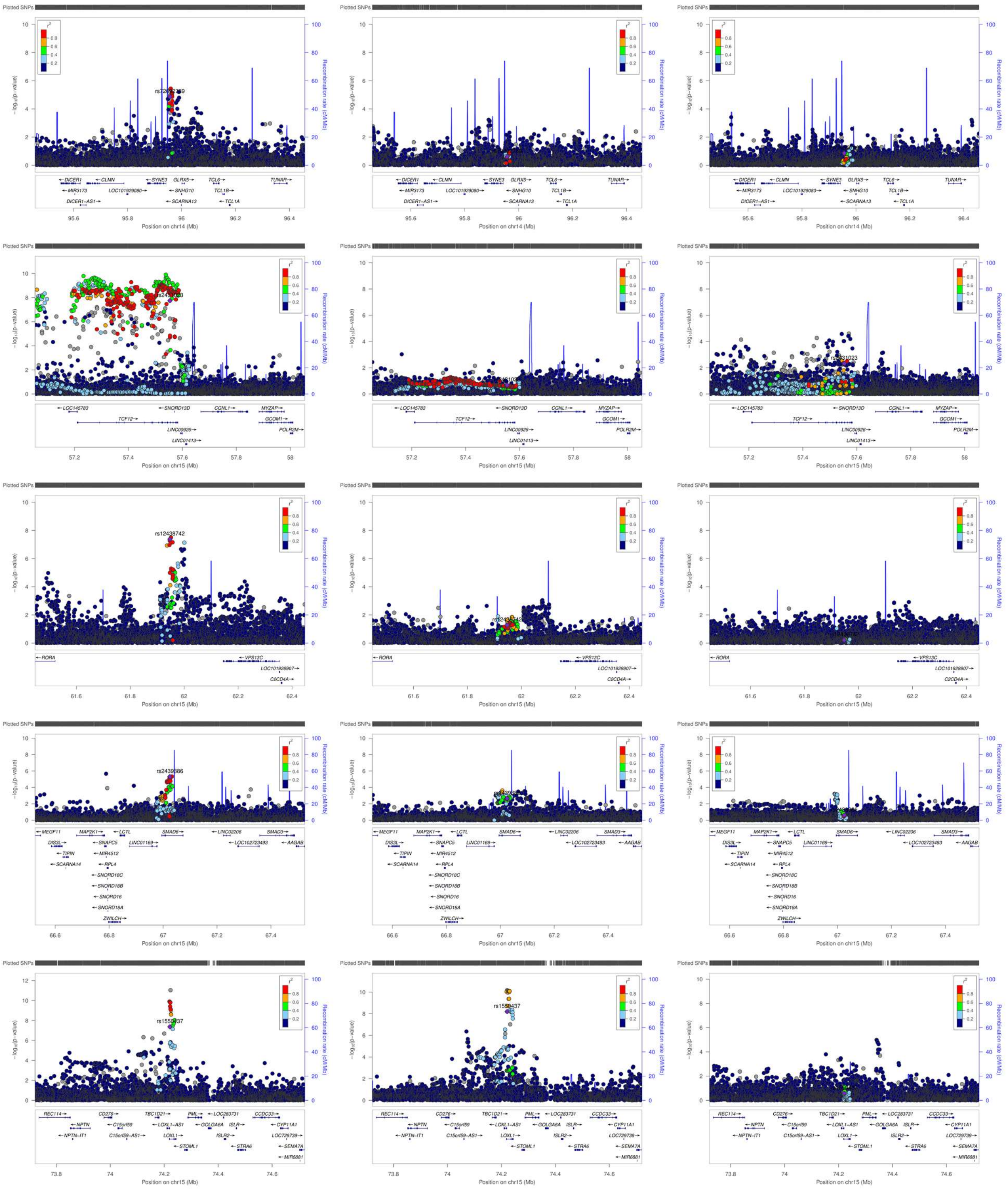

Plotted SNPs
-log10(p-value)
Recombination rate (cM/Mb)
Position on chr14 (Mb)
Position on chr15 (Mb)
rs12438742
rs2439686
rs1550437
DICER1
CLMN
SYNE3
GLRX5
TCL6
TUNAR
MIR3173
LOC101929080
SNHG10
TCL1B
DICER1-AS1
SCARNA13
TCL1A
LOC145783
SNORD13D
CGNL1
MYZAP
TCF12
GCOM1
LINC00926
POLR2M
LINC01413
RORA
VPS13C
LOC101928907
C2CD4A
MEGF11
MAP2K1
LCTL
SMAD6
LINC02206
SMAD3
DIS3L
SNAPC5
LINC01169
LOC102723493
AAGAB
TIPIN
MIR4512
SCARNA14
RPL4
SNORD18C
SNORD18B
SNORD16
SNORD18A
ZWILCH
REC114
CD276
TBC1D21
PML
LOC283731
CCDC33
NPTN
C15orf59
LOXL1-AS1
GOLGA6A
ISLR
CYP11A1
NPTN-IT1
C15orf59-AS1
LOXL1
ISLR2
LOC729739
STOML1
STRA6
SEMA7A
MIR6881

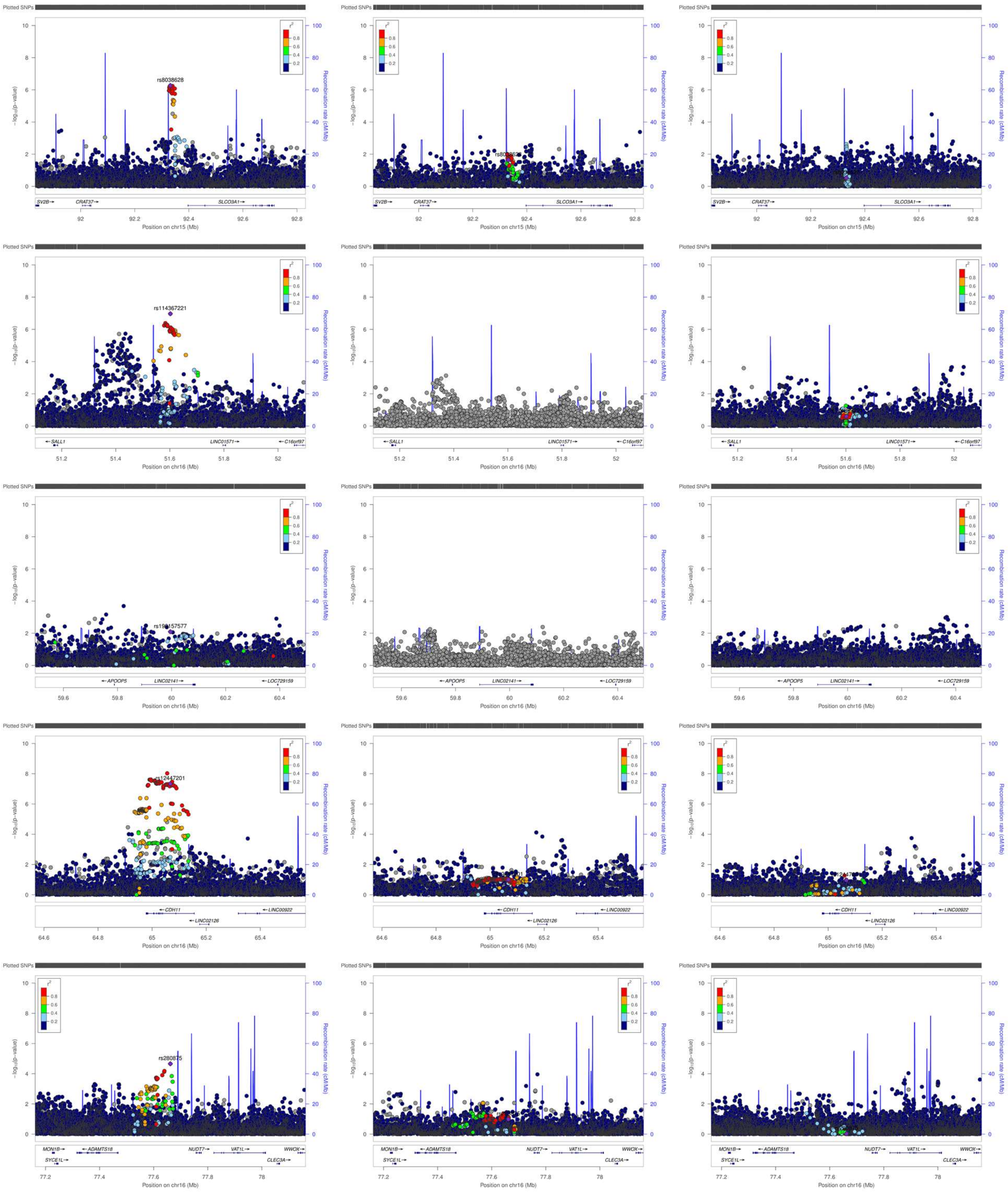

Plotted SNPs
rs8038628
rs114367221
rs12447201
rs280875
-log10(p-value)
Recombination rate (cM/Mb)
Position on chr15 (Mb)
Position on chr16 (Mb)
SV2B
CRAT37
SLCO3A1
SALL1
LINC01571
C16orf97
APOOP5
LINC02141
LOC729159
CDH11
LINC02126
LINC00922
MON1B
SYCE1L
ADAMTS18
NUDT7
VAT1L
WWOX
CLEC3A

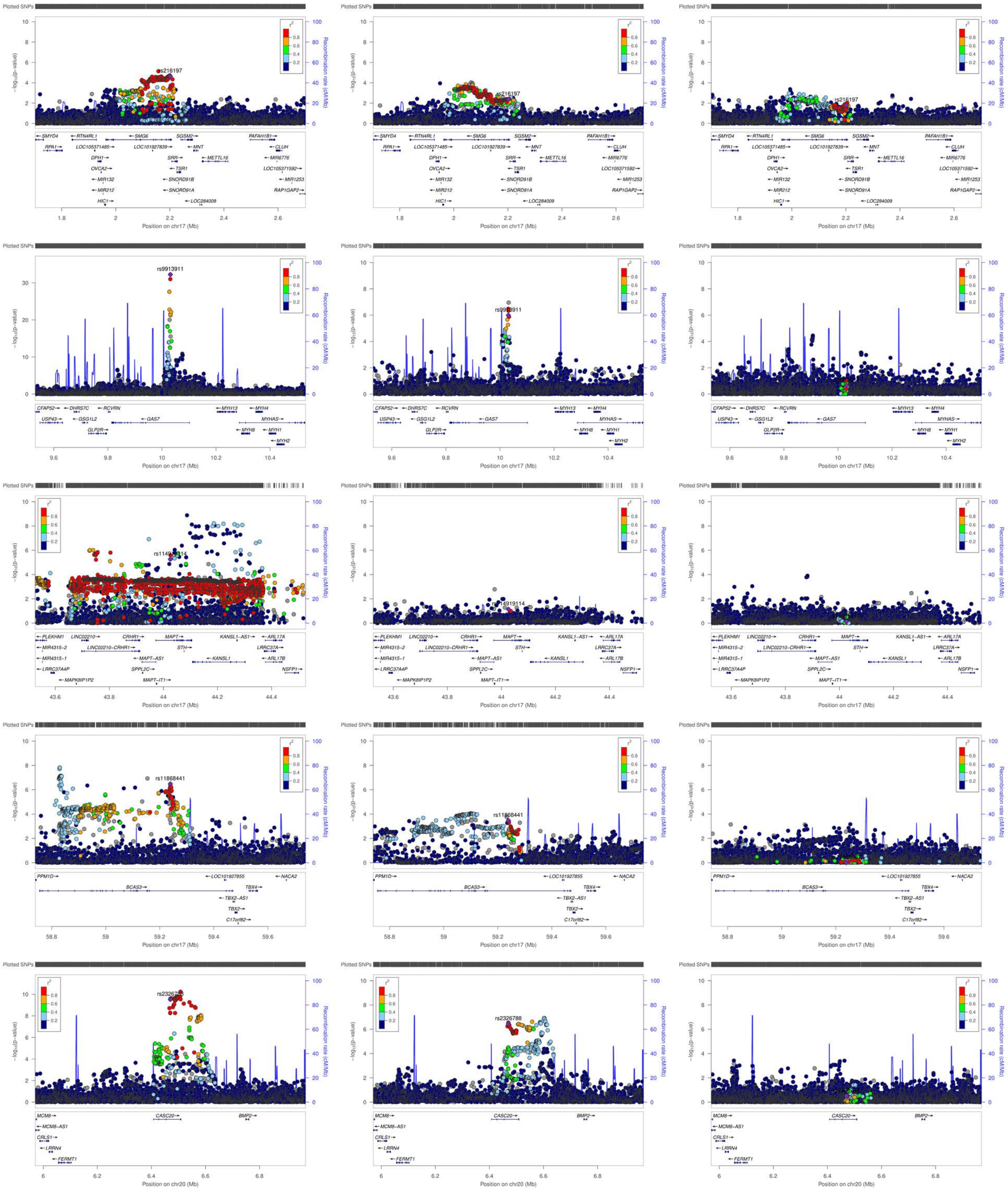
Plotted SNPs
-log10(p-value)
Recombination rate (cM/Mb)
r2
rs216197
rs9913911
rs11868441
rs2326788
SMYD4
RTN4RL1
SMG6
SGSM2
PAFAH1B1
RPA1
LOC105371485
LOC101927839
MNT
CLUH
DPH1
SRR
METTL16
MIR6776
OVCA2
TSR1
LOC105371592
MIR132
SNORD91B
MIR1253
MIR212
SNORD91A
RAP1GAP2
HIC1
LOC284009
Position on chr17 (Mb)
CFAP52
DHRS7C
RCVRN
MYH13
MYH4
USP43
GSG1L2
GAS7
MYHAS
GLP2R
MYH8
MYH1
MYH2
PLEKHM1
LINC02210
CRHR1
MAPT
KANSL1-AS1
ARL17A
MIR4315-2
LINC02210-CRHR1
STH
LRRC37A
MIR4315-1
MAPT-AS1
KANSL1
ARL17B
LRRC37A4P
SPPL2C
NSFP1
MAPK8IP1P2
MAPT-IT1
PPM1D
LOC101927855
NACA2
BCAS3
TBX4
TBX2-AS1
TBX2
C17orf82
MCM8
CASC20
BMP2
MCM8-AS1
CRLS1
LRRN4
FERMT1
Position on chr20 (Mb)

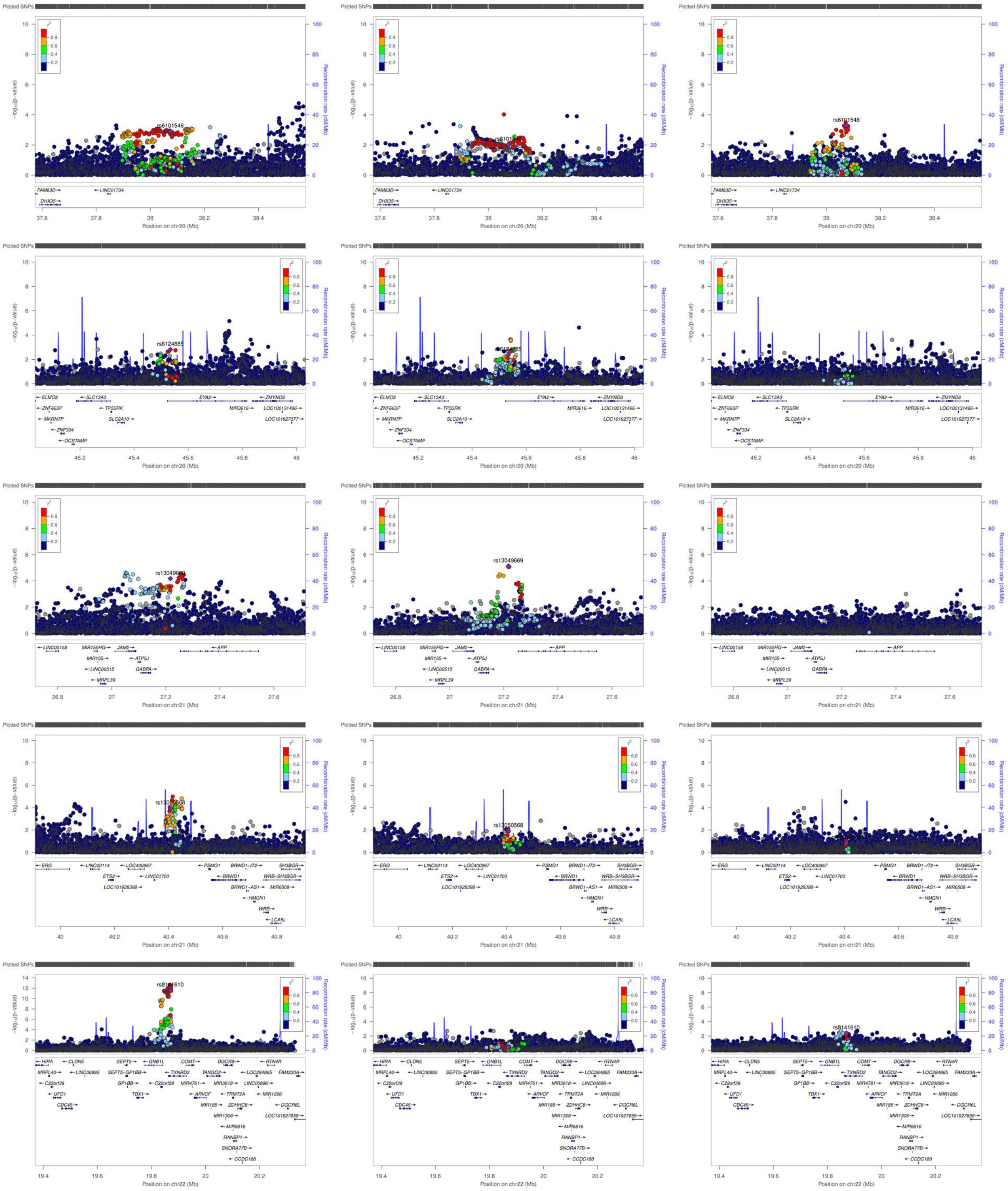

Plotted SNPs
-log10(p-value)
Recombination rate (cM/Mb)
r2
rs6101546
Position on chr20 (Mb)
rs6124885
rs13049669
Position on chr21 (Mb)
rs13050568
rs8141610
Position on chr22 (Mb)

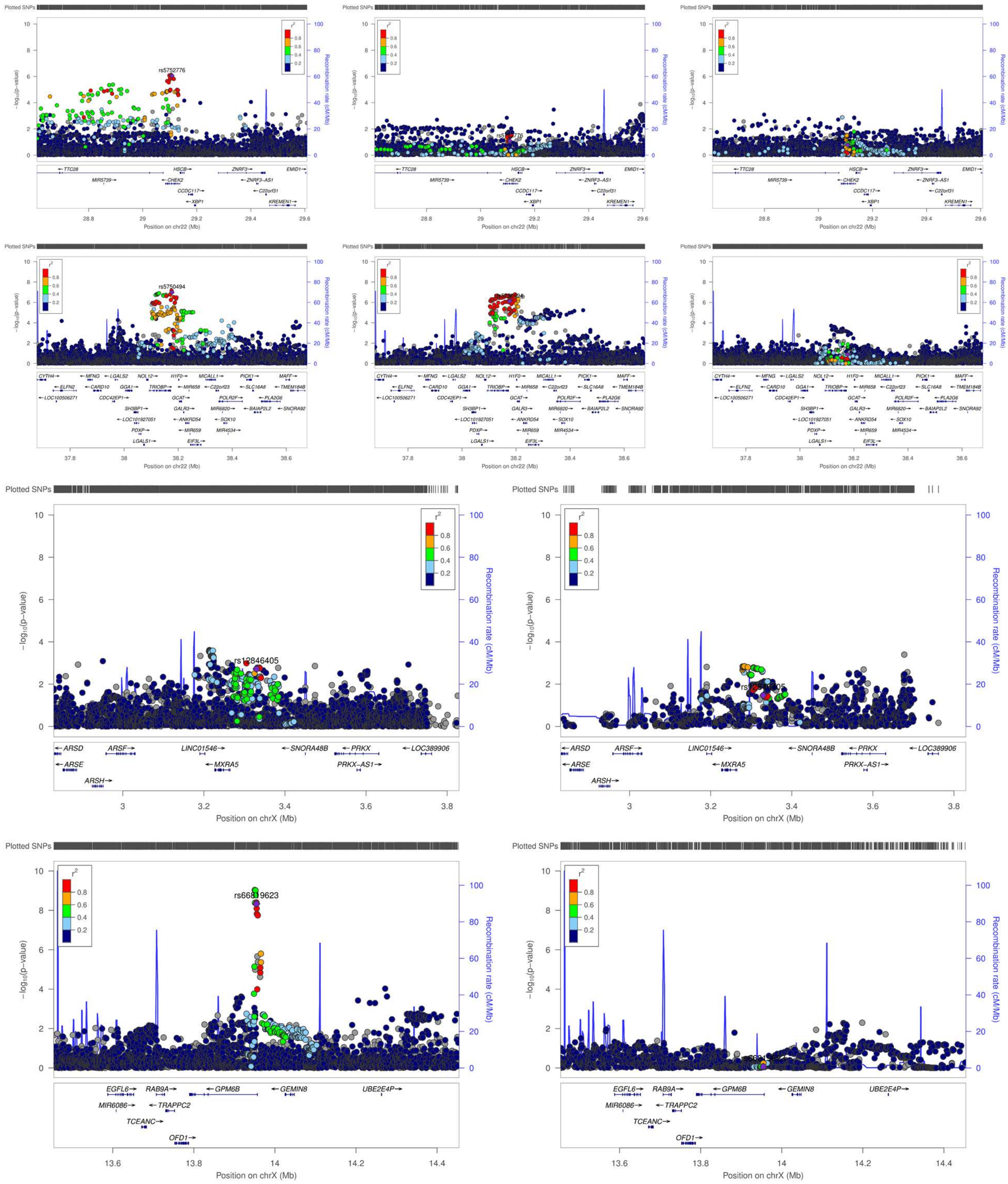

Plotted SNPs
-log10(p-value)
Recombination rate (cM/Mb)
r2
0.8
0.6
0.4
0.2
rs5752776
TTC28
MIR5739
HSCB
CHEK2
CCDC117
XBP1
ZNRF3
ZNRF3-AS1
C22orf31
KREMEN1
EMID1
Position on chr22 (Mb)
rs5750494
CYTH4
ELFN2
LOC100506271
MFNG
CARD10
CDC42EP1
LGALS2
GGA1
SH3BP1
LOC101927051
PDXP
LGALS1
NOL12
TRIOBP
H1F0
GCAT
GALR3
ANKRD54
MIR658
MIR659
EIF3L
MICALL1
C22orf23
POLR2F
MIR6820
SOX10
MIR4534
PICK1
SLC16A8
BAIAP2L2
MAFF
TMEM184B
PLA2G6
SNORA92
rs12846405
ARSD
ARSE
ARSH
ARSF
LINC01546
MXRA5
SNORA48B
PRKX
PRKX-AS1
LOC389906
Position on chrX (Mb)
rs66819623
EGFL6
MIR6086
TCEANC
RAB9A
TRAPPC2
OFD1
GPM6B
GEMIN8
UBE2E4P

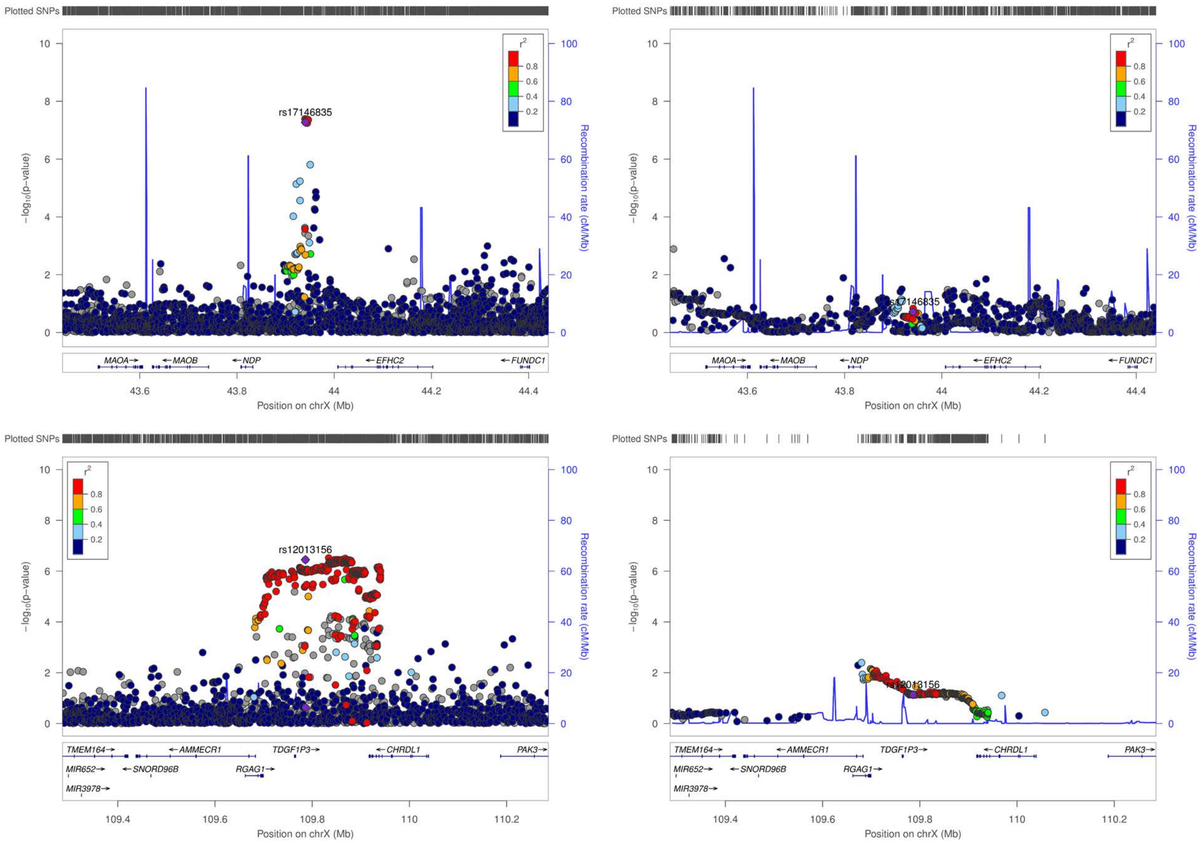

Plotted SNPs
rs17146835
$-\log_{10}(p\text{-value})$
Recombination rate (cM/Mb)
$r^2$
0.8
0.6
0.4
0.2
MAOA→
←MAOB
←NDP
←EFHC2
←FUNDC1
43.6
43.8
44
44.2
44.4
Position on chrX (Mb)
Plotted SNPs
rs17146835
$-\log_{10}(p\text{-value})$
Recombination rate (cM/Mb)
MAOA→
←MAOB
←NDP
←EFHC2
←FUNDC1
Position on chrX (Mb)
Plotted SNPs
rs12013156
$-\log_{10}(p\text{-value})$
Recombination rate (cM/Mb)
TMEM164→
←AMMECR1
TDGF1P3→
←CHRDL1
PAK3→
MIR652→
←SNORD96B
RGAG1→
MIR3978→
109.4
109.6
109.8
110
110.2
Position on chrX (Mb)
Plotted SNPs
rs12013156
$-\log_{10}(p\text{-value})$
Recombination rate (cM/Mb)
TMEM164→
←AMMECR1
TDGF1P3→
←CHRDL1
PAK3→
MIR652→
←SNORD96B
RGAG1→
MIR3978→
Position on chrX (Mb)

**Supplementary Figure 4. Enrichment of the expression of the significant genes identified by MAGMA in GTEX tissues.** The horizontal line indicates the significance threshold (P<1e-03), Bonferroni-corrected for 53 GTEX tissues tested.

**A) Results from the European meta-analysis.**

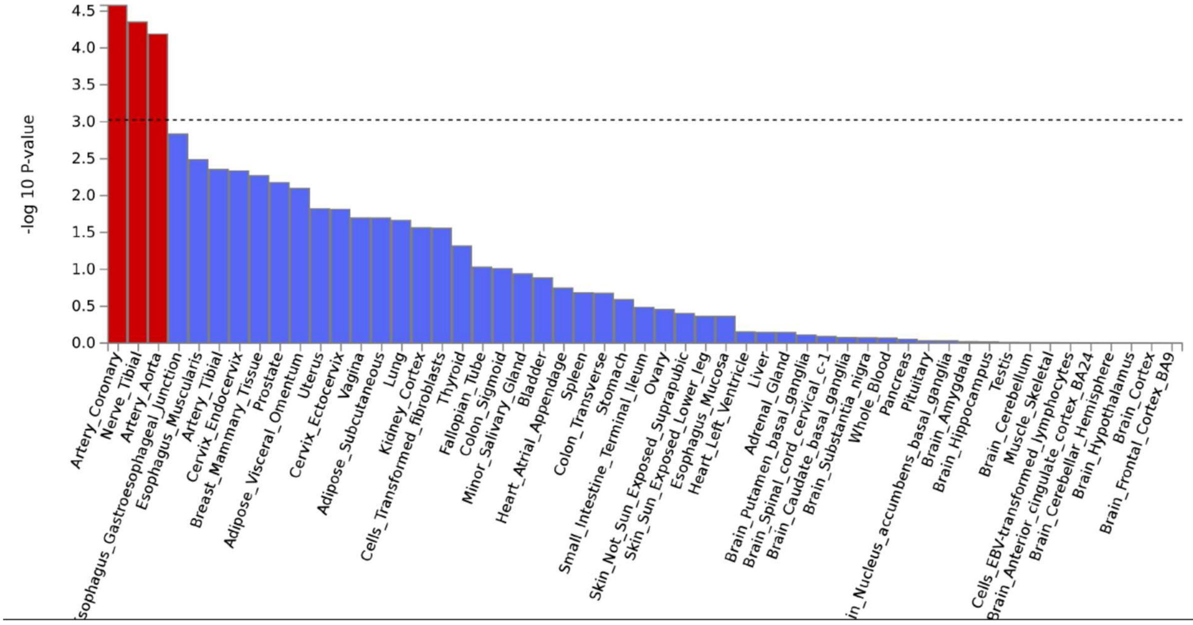


**B) Results from the cross-ancestry meta-analysis.**

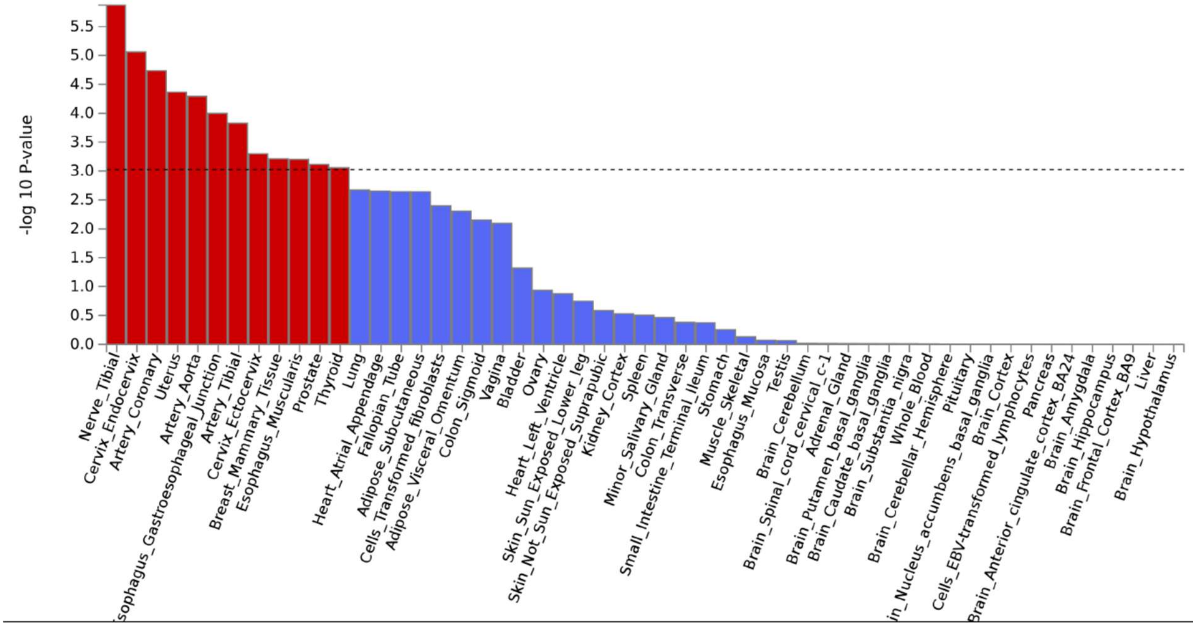

**Supplementary Figure 5. Chromatin and eQTL interactions for the POAG risk loci.** The interactions are shown for each chromosome separately. Orange lines show chromatin interactions and green lines eQTL interactions. Green genes are the eQTL-mapped genes, orange genes are mapped by chromatin interactions, and red genes by both. The outer layers are Manhattan plots with the most significant SNPs shown.

##### Chromosome 1

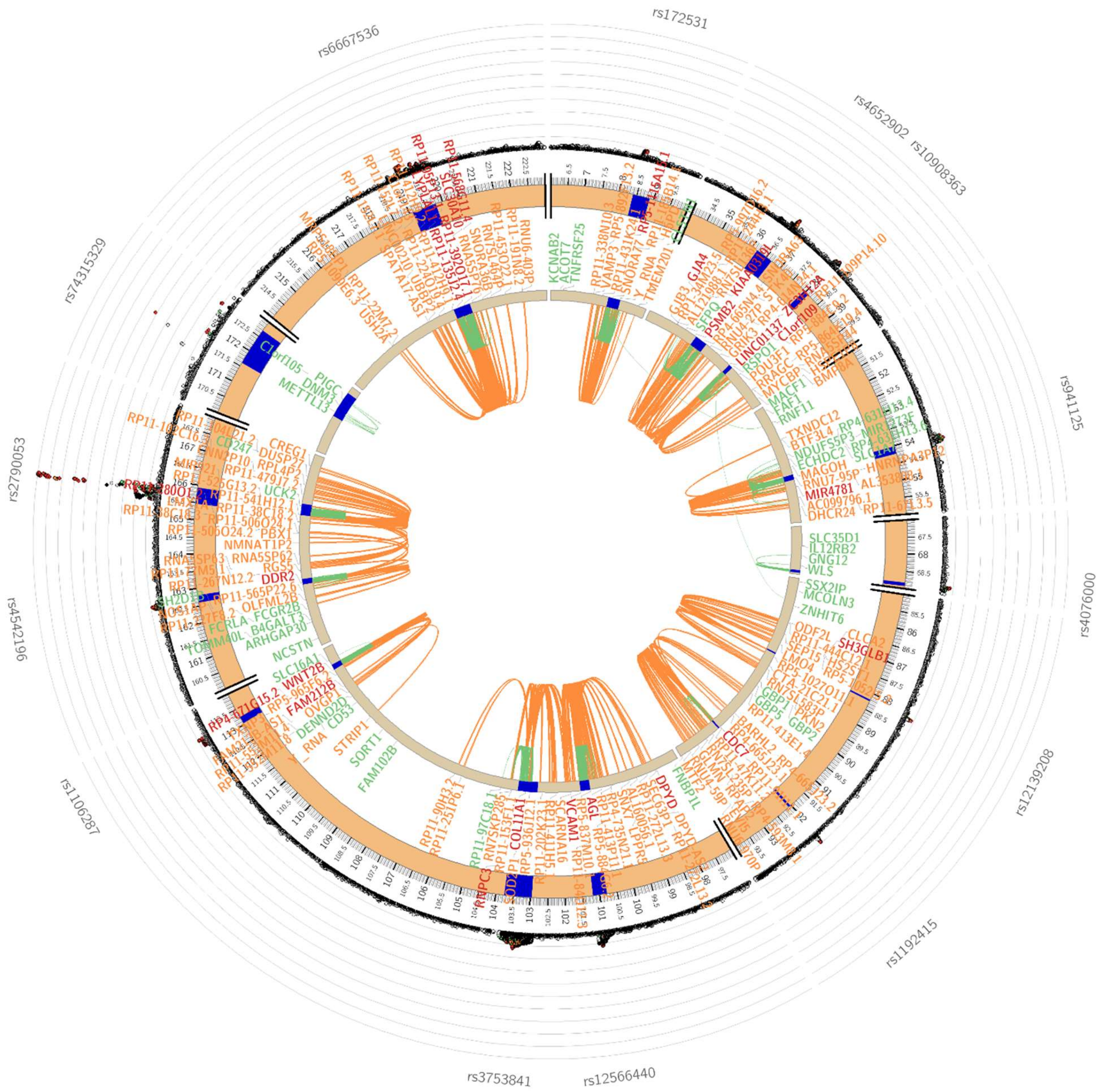

#### Chromosome 2

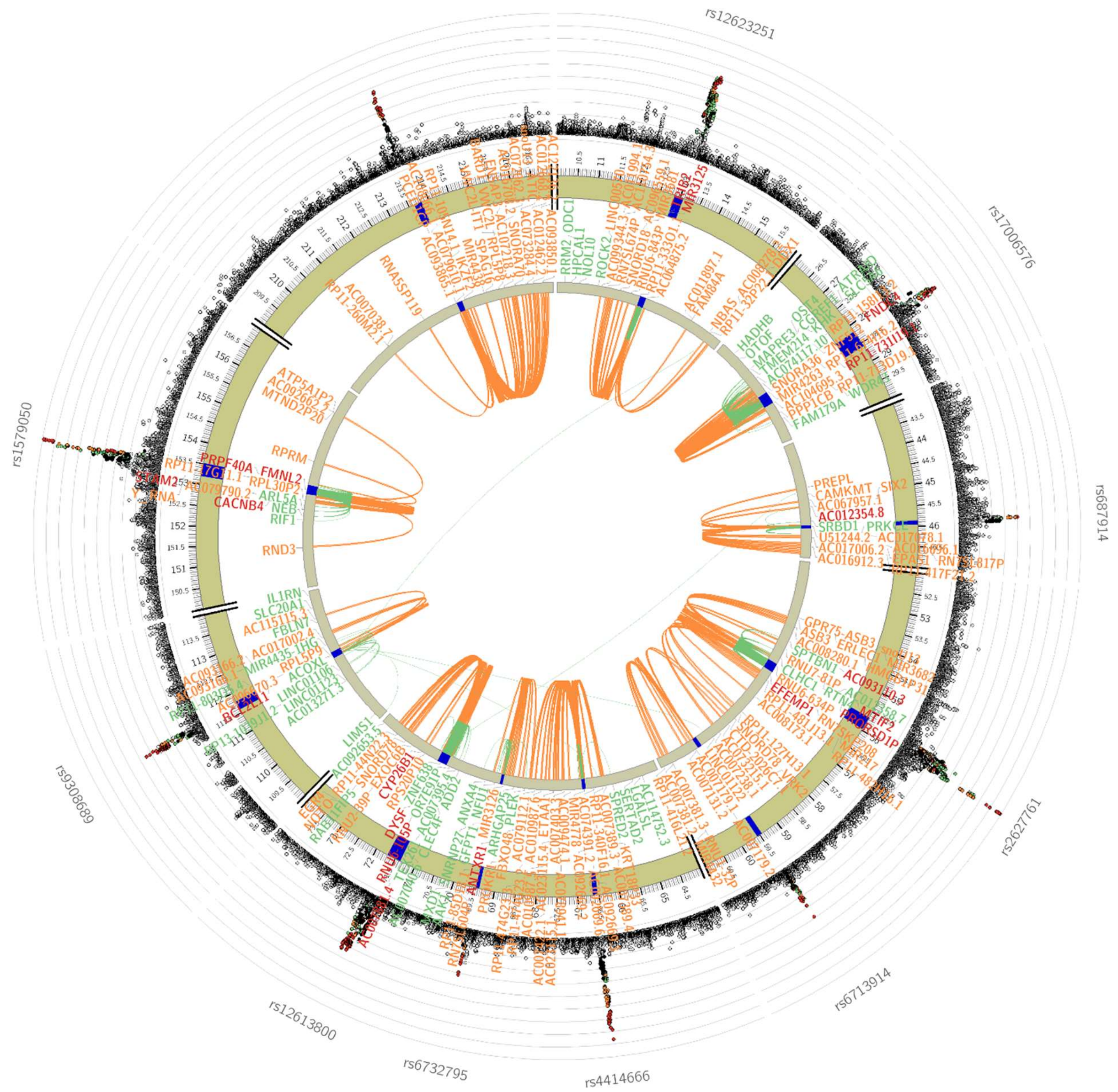

**Chromosome 3**

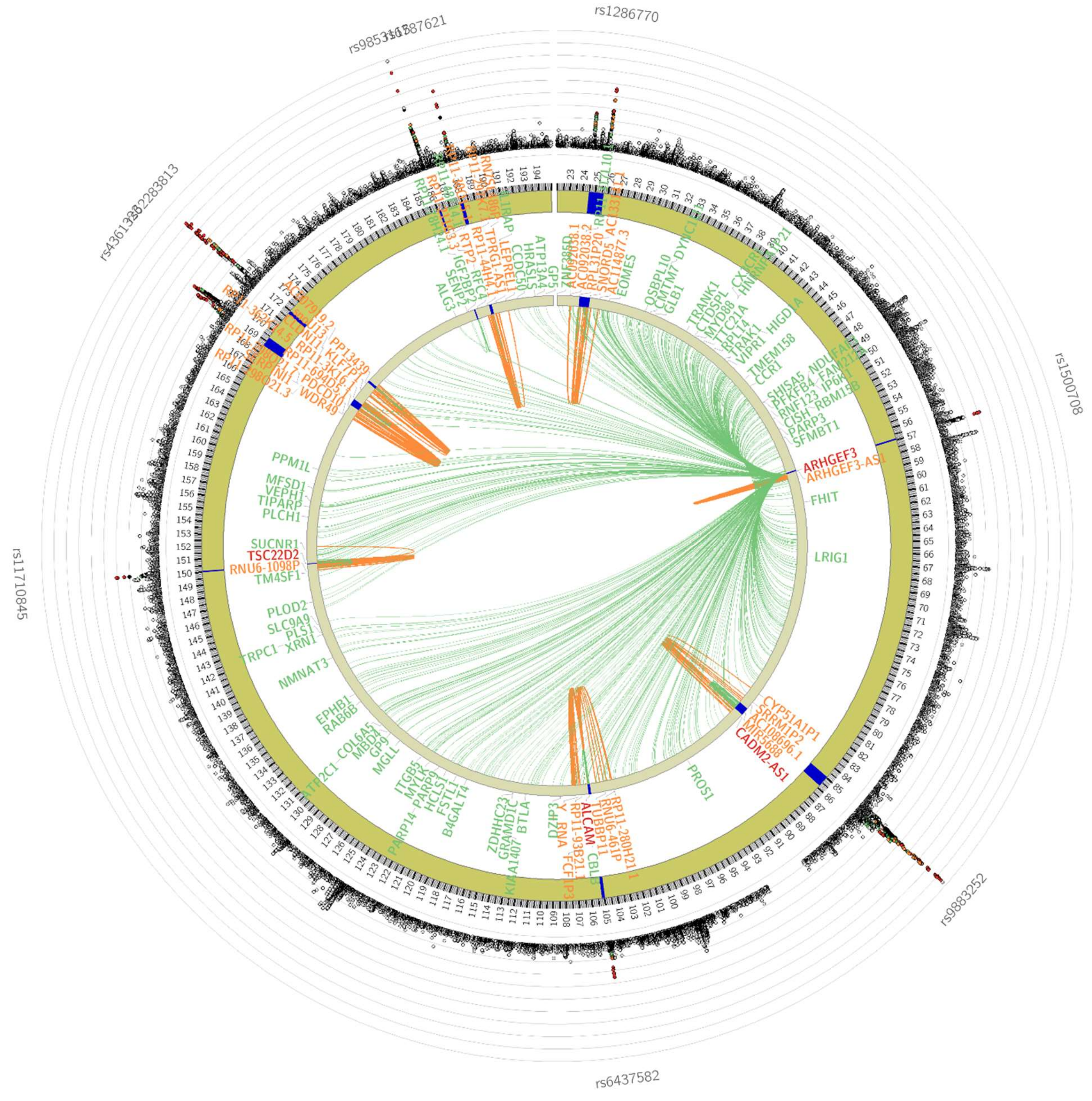

**Chromosome 4**

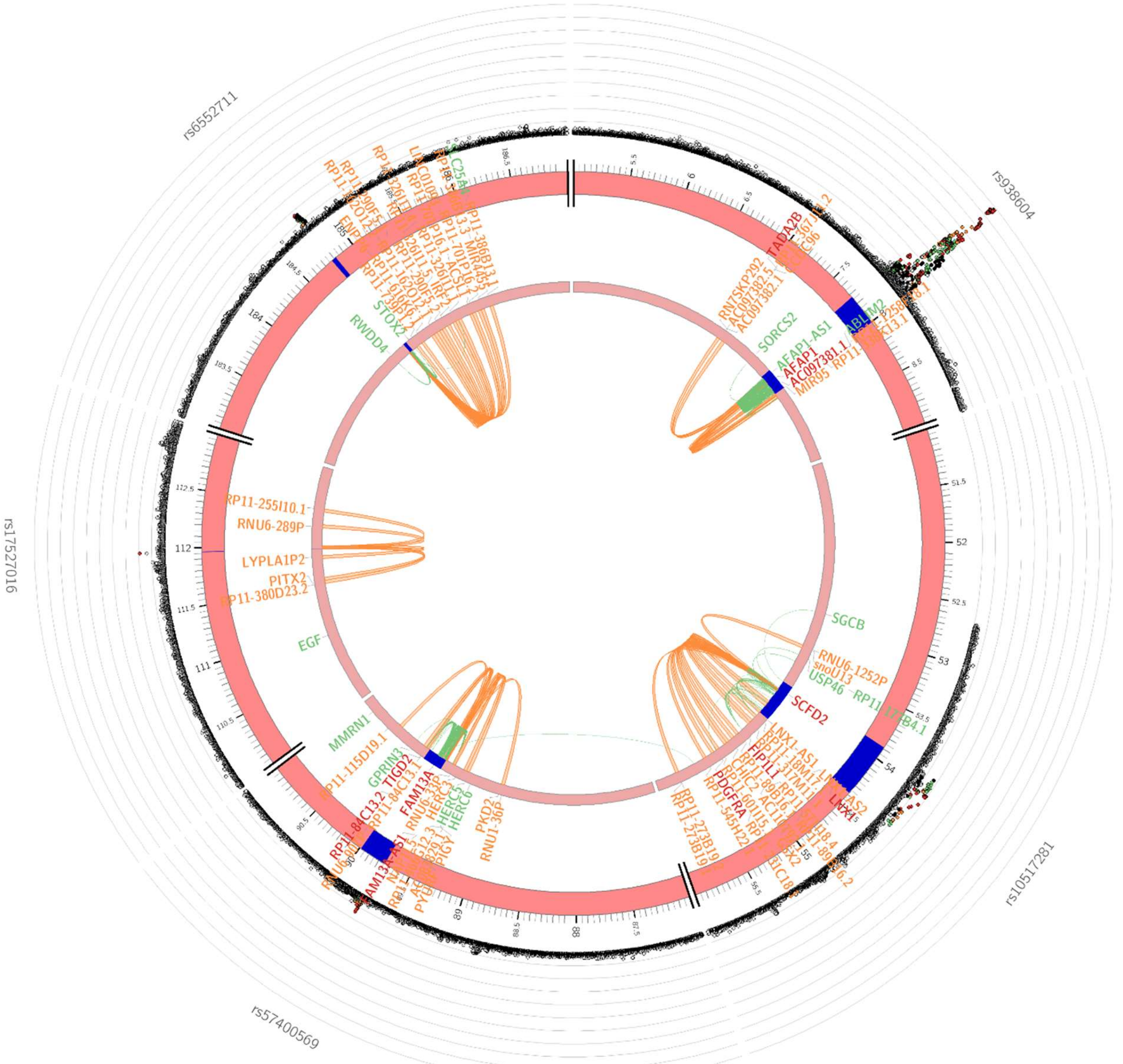

**Chromosome 5**

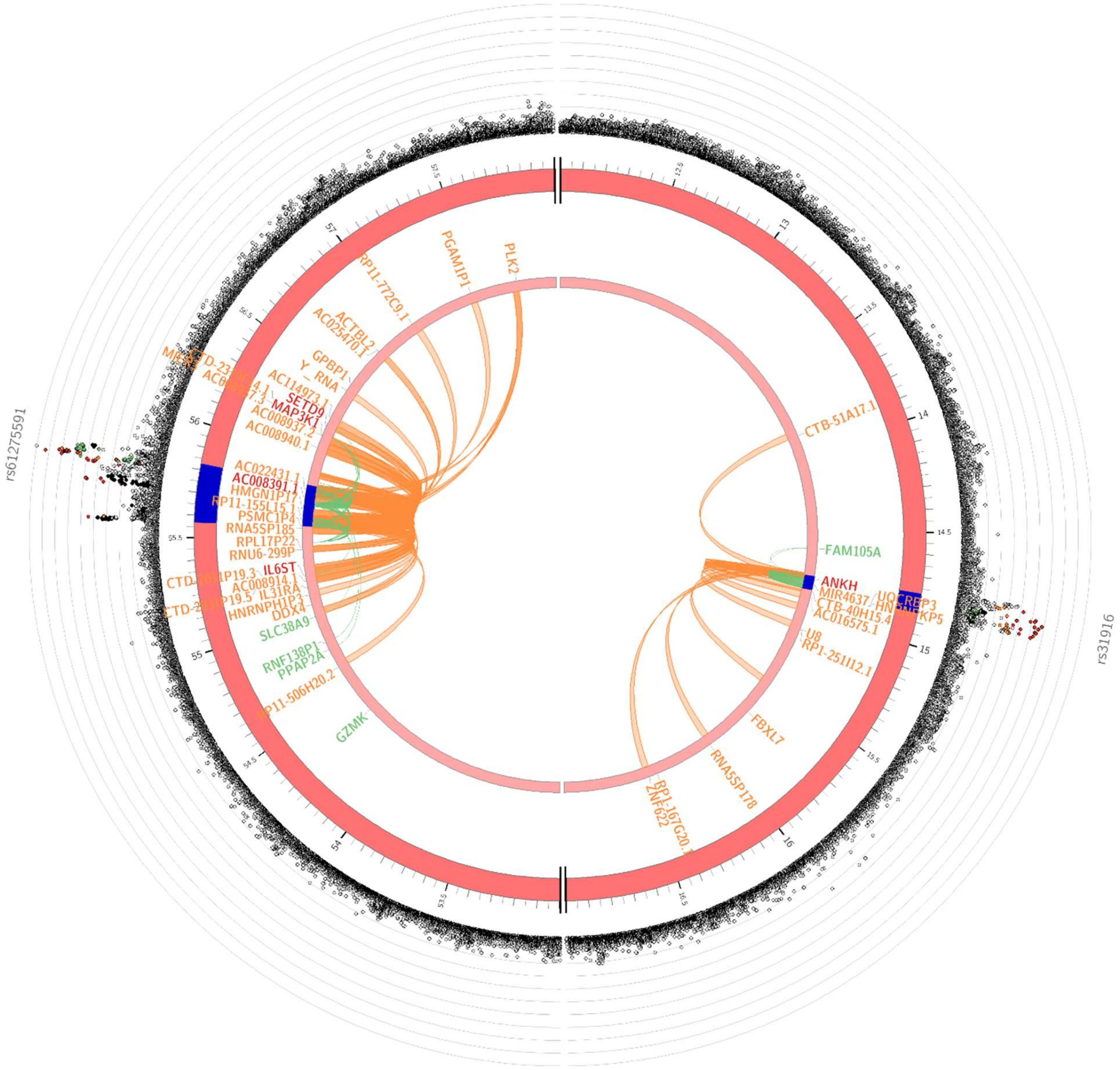

#### Chromosome 6

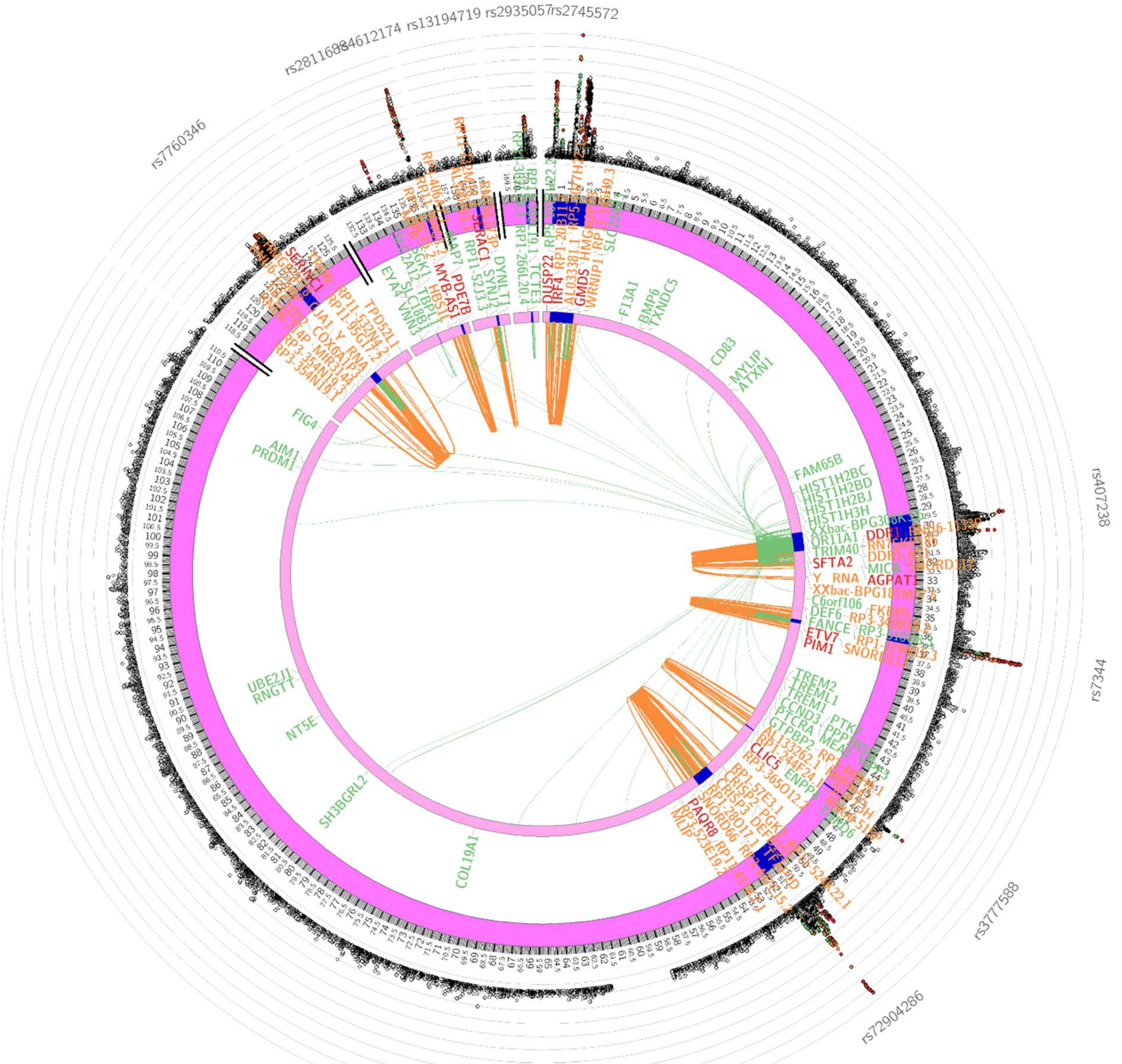

#### Chromosome 7

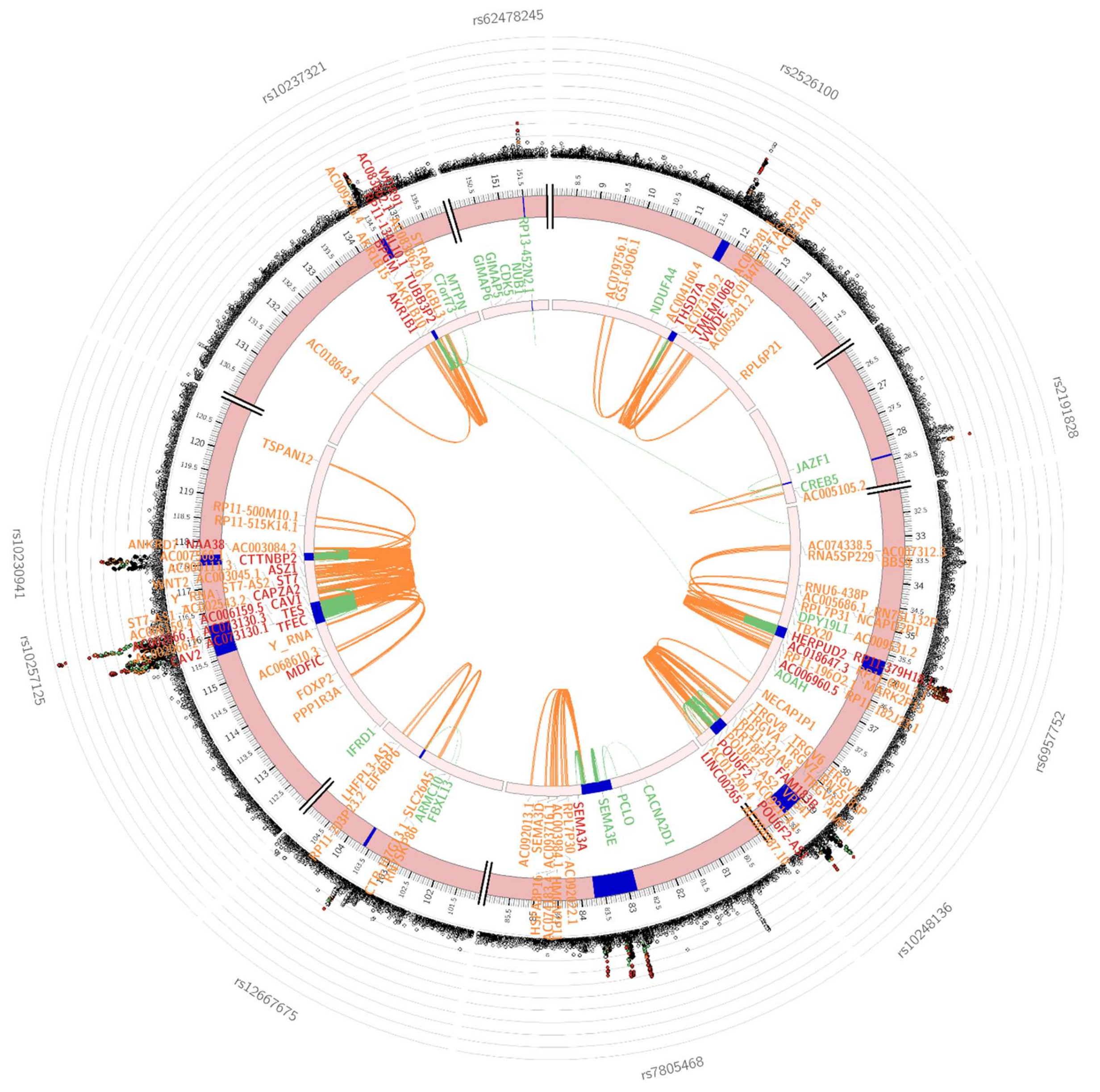

#### Chromosome 8

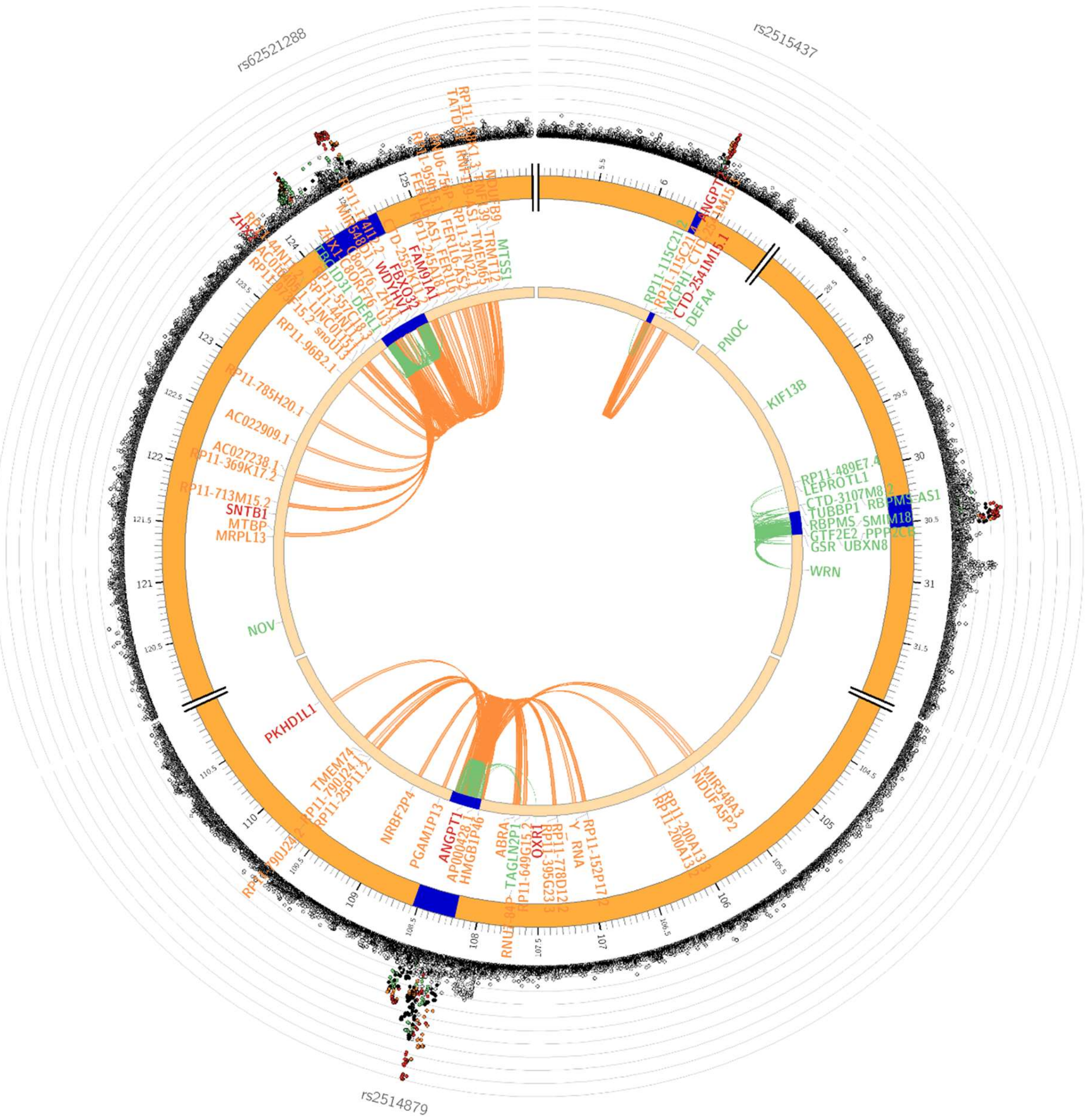

#### Chromosome 9

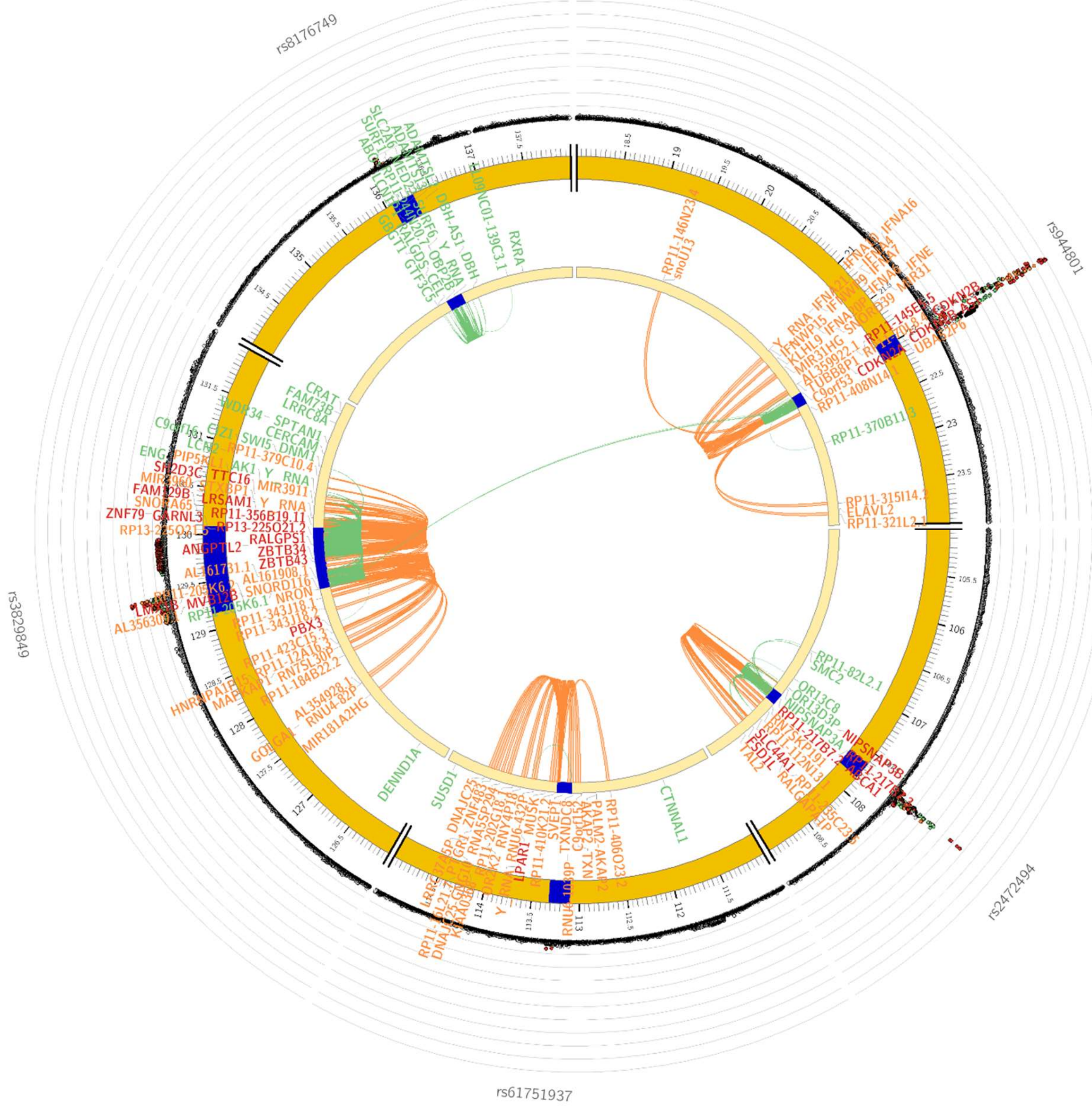

#### Chromosome 10

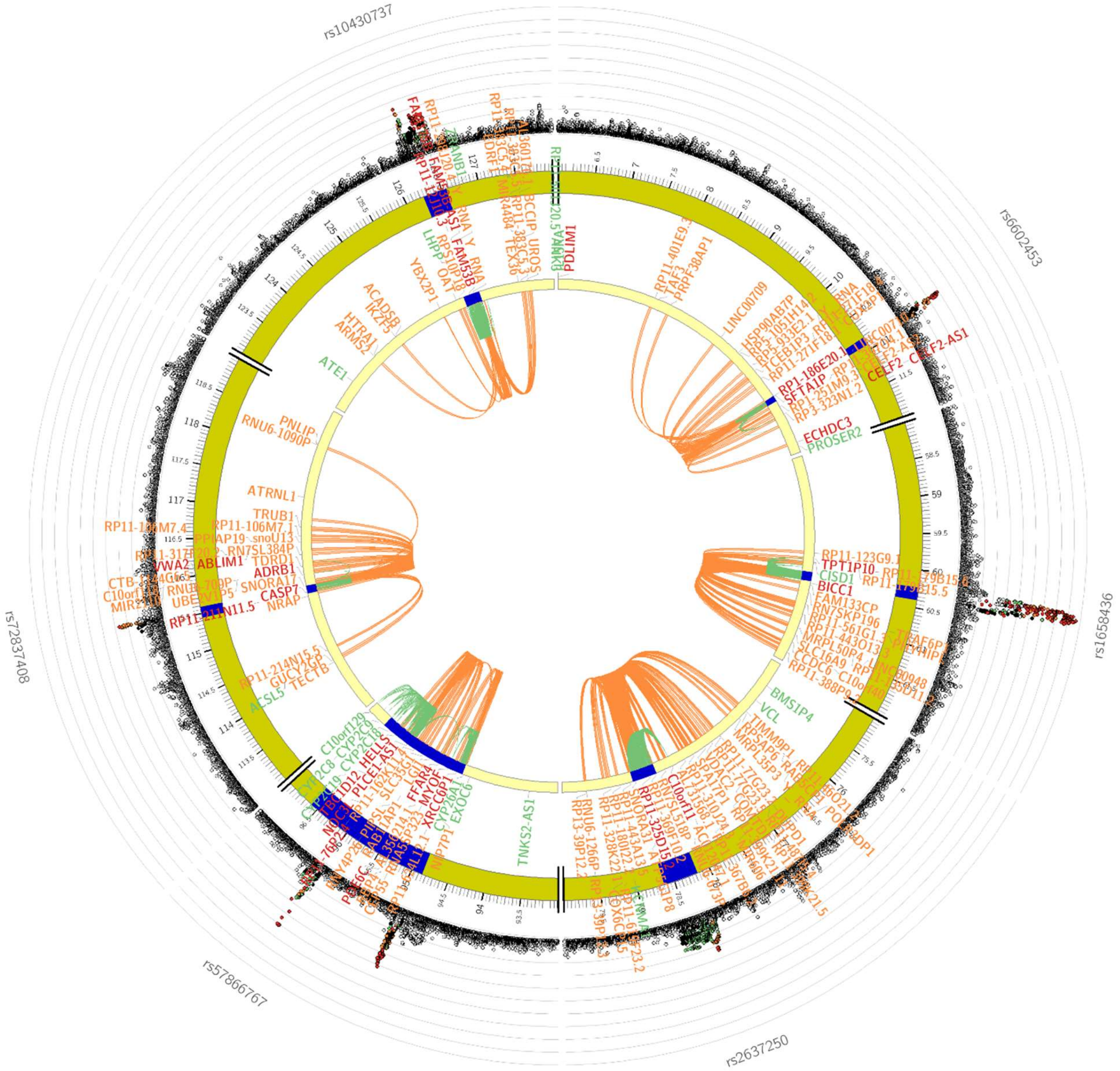

#### Chromosome 11

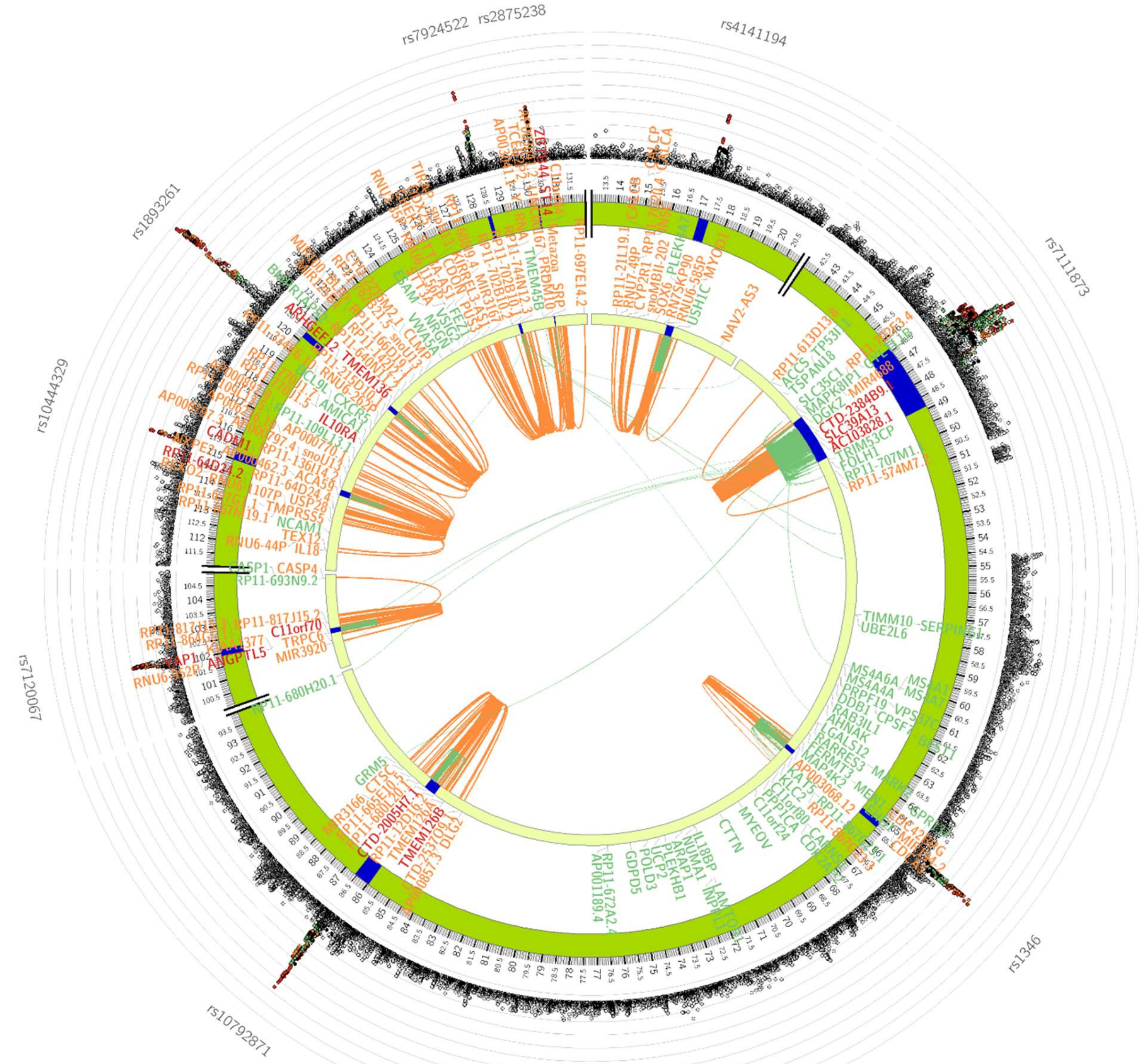

#### Chromosome 12

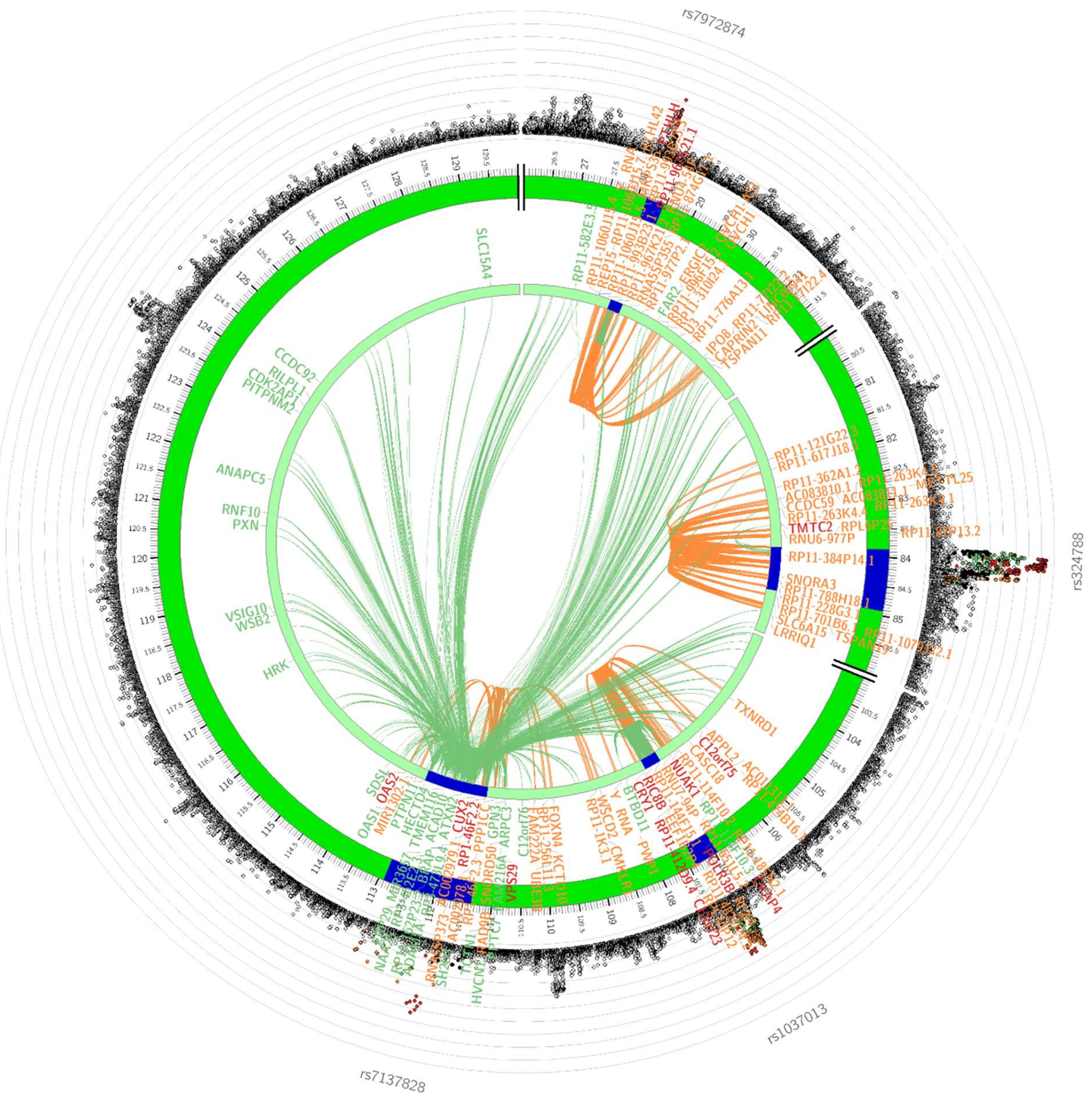

#### Chromosome 13

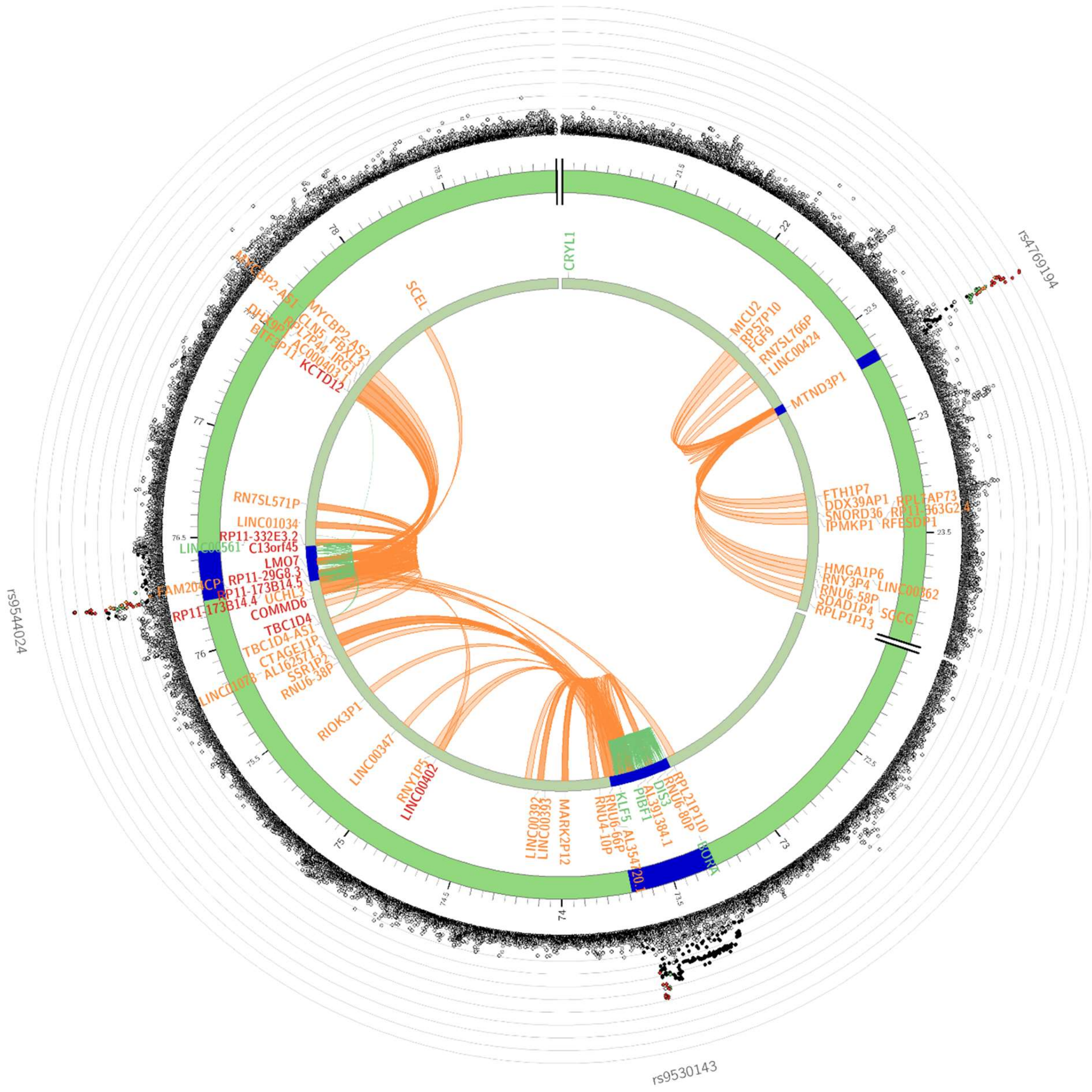

#### Chromosome 14

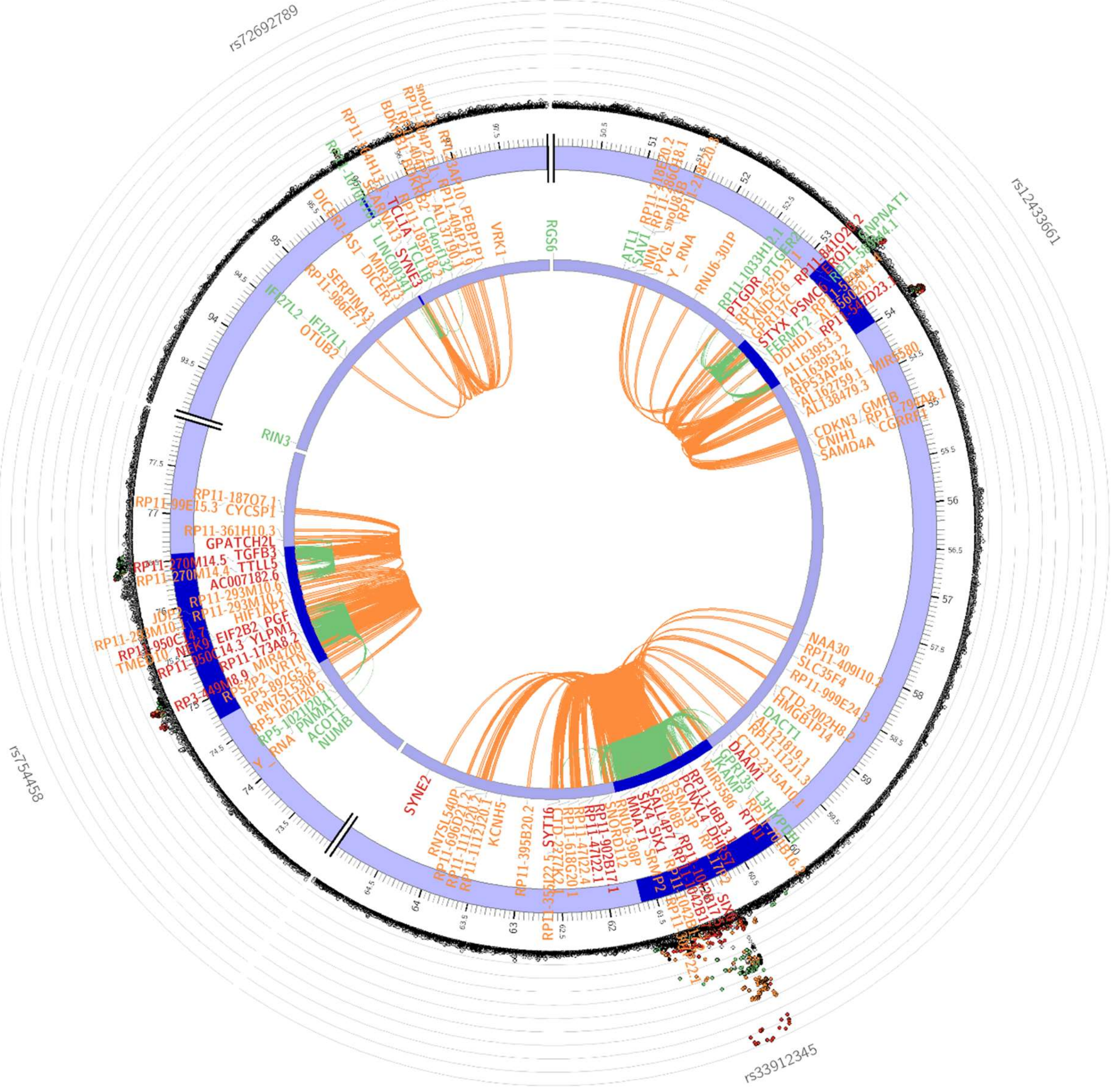

#### Chromosome 15

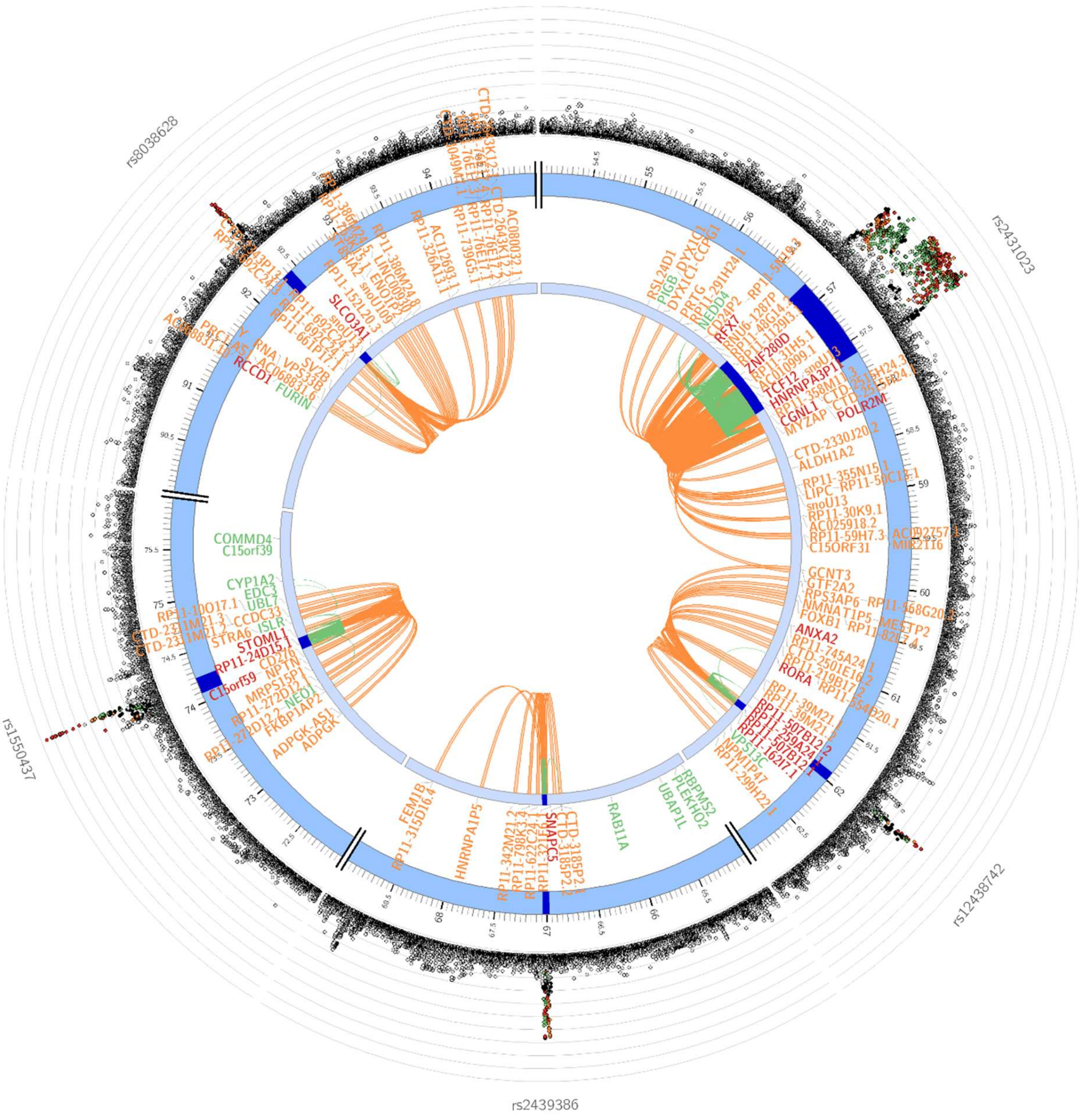

#### Chromosome 16

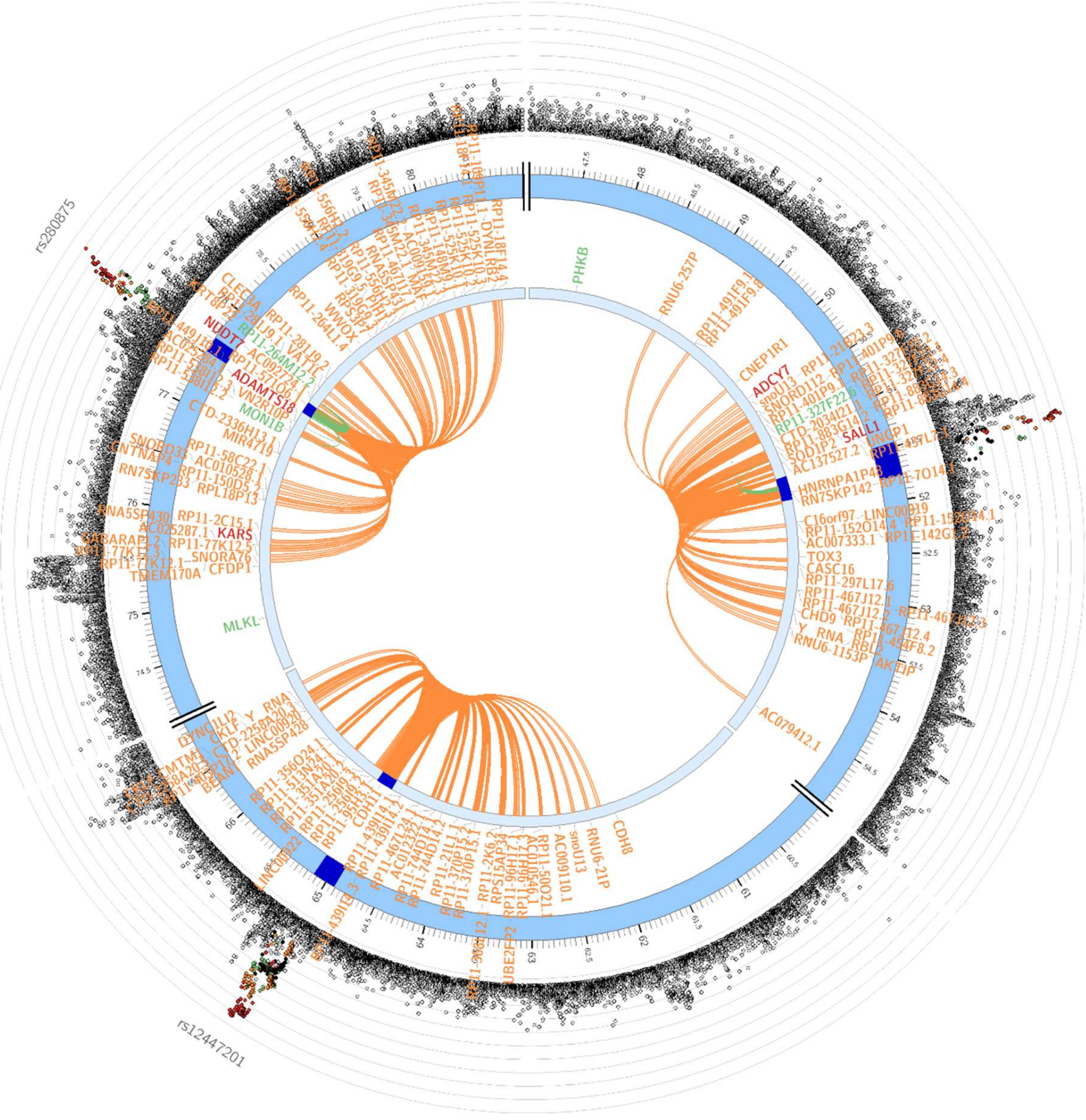

#### Chromosome 17

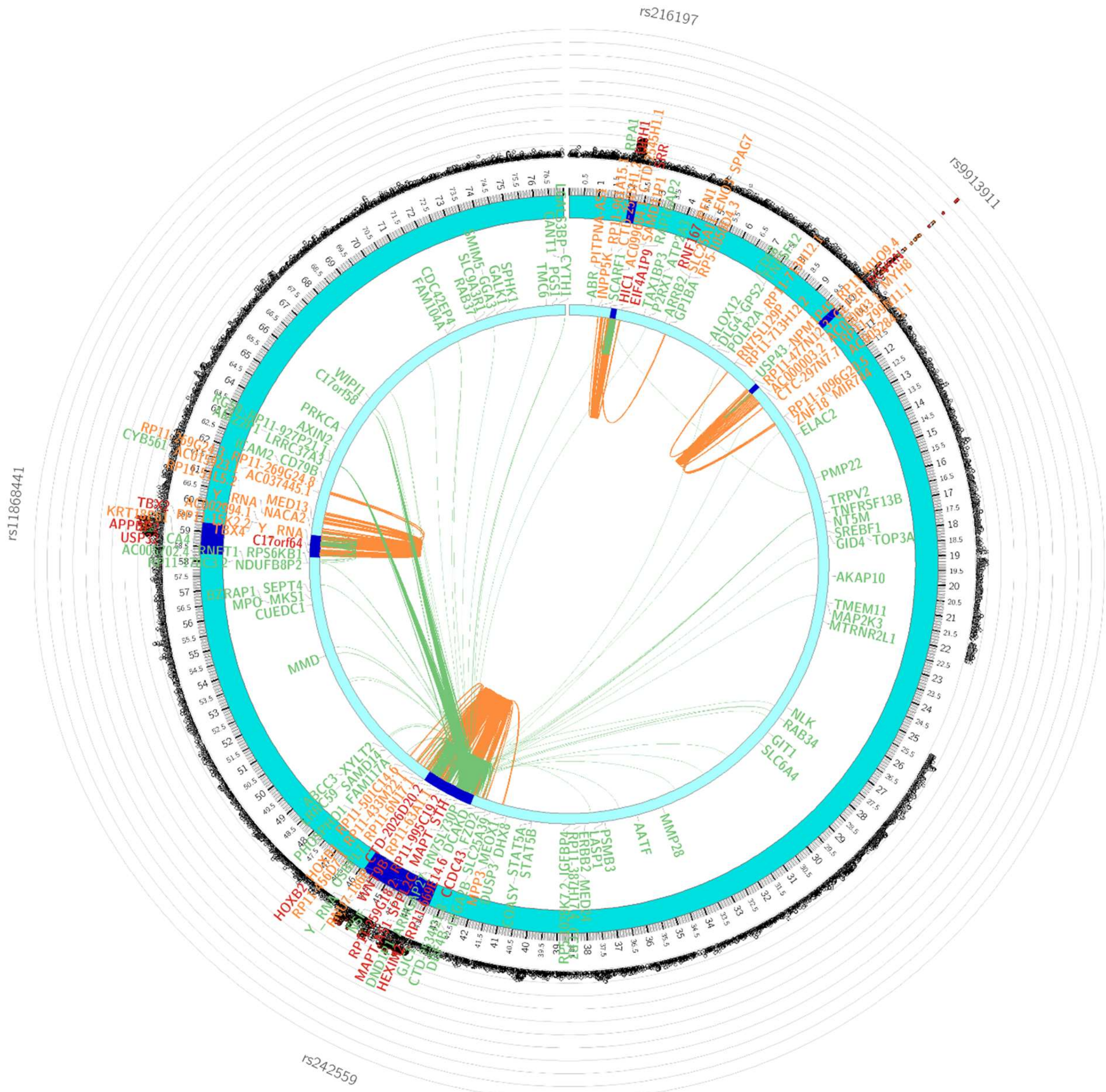

### Chromosome 20

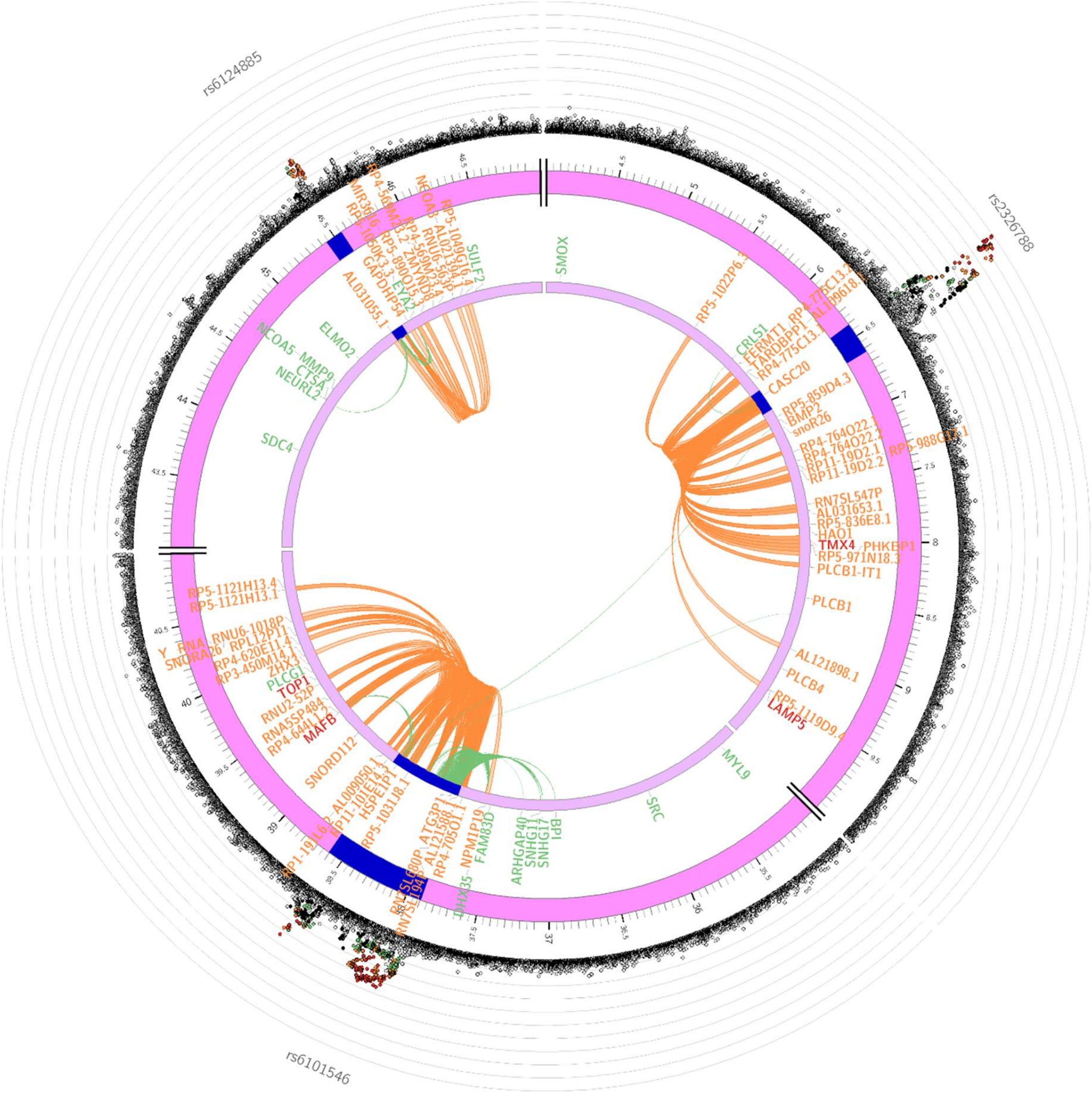

#### Chromosome 21

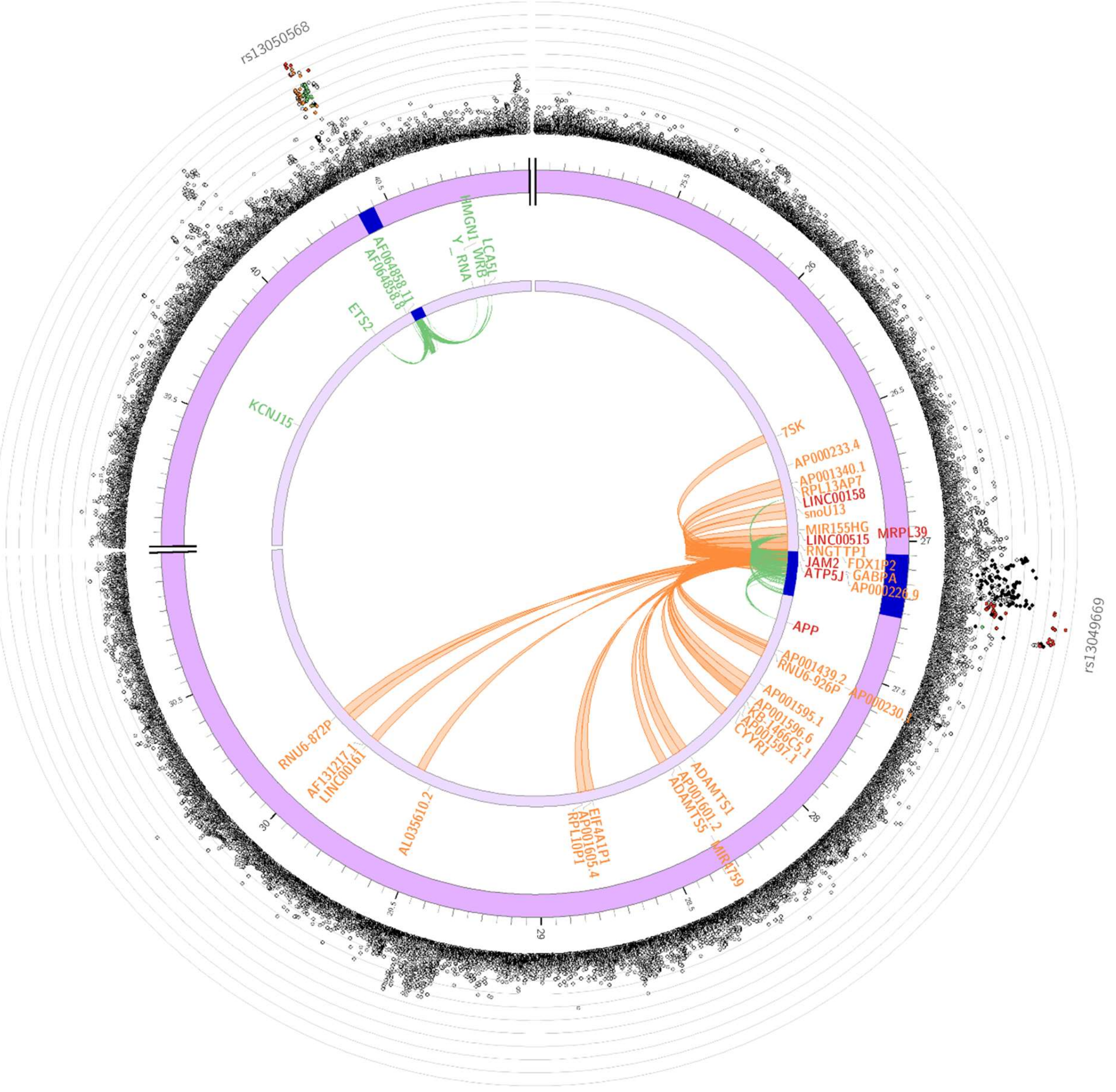

#### Chromosome 22

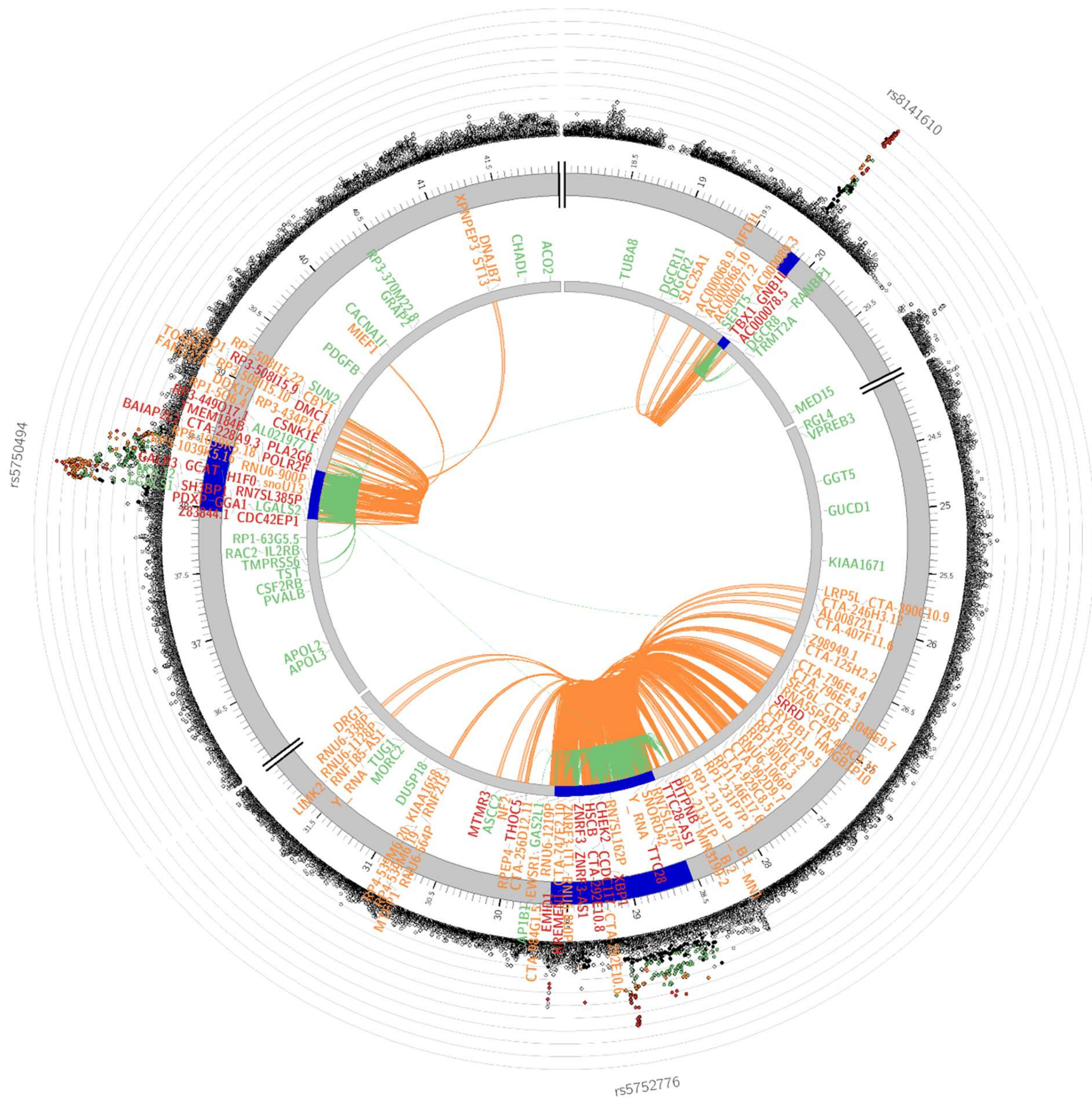

**Chromosome 23**

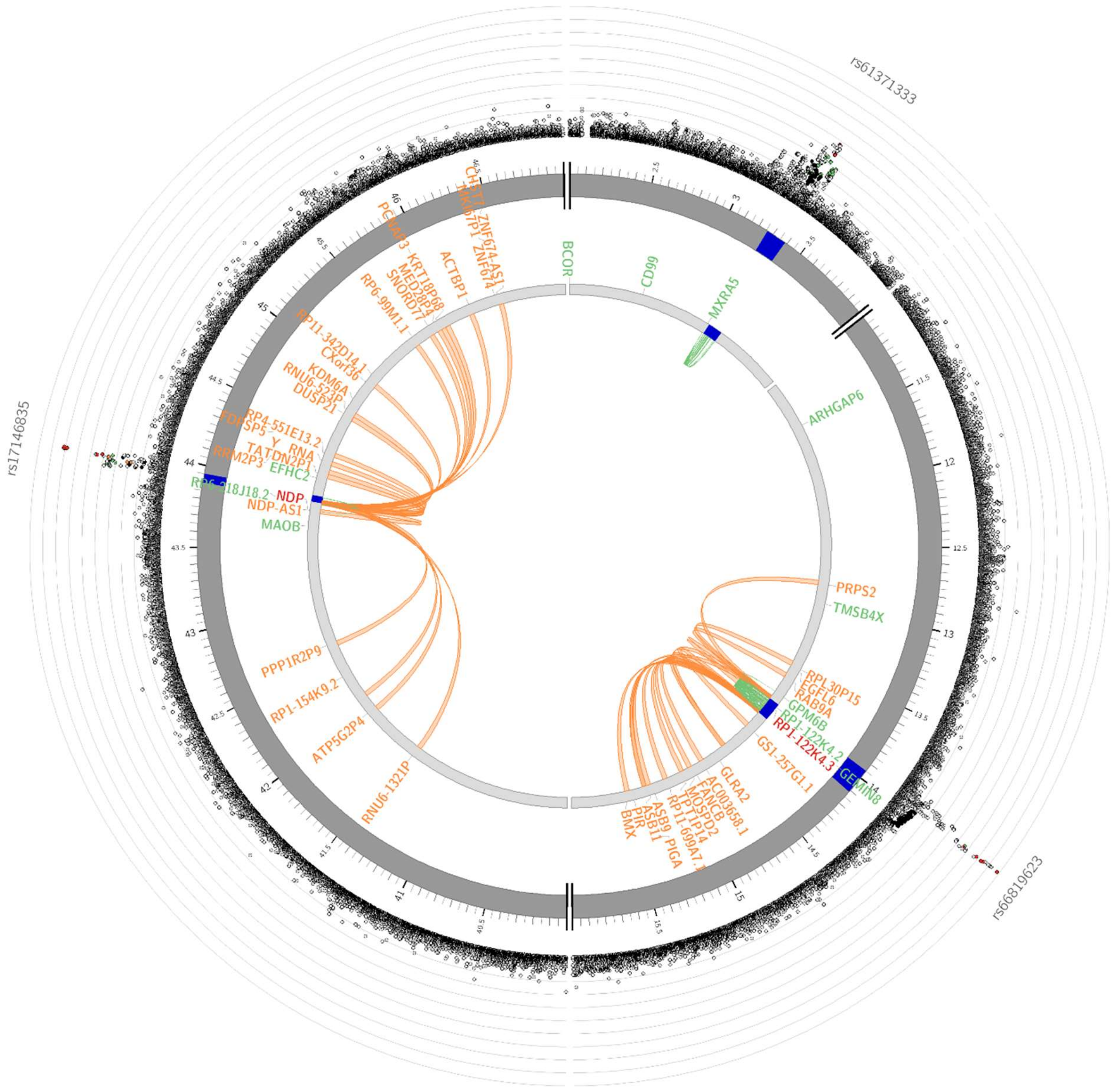

**Supplementary Figure 6. Gene expression heatmaps in eye tissues.**

**A) Gene expression of the novel POAG risk genes.**

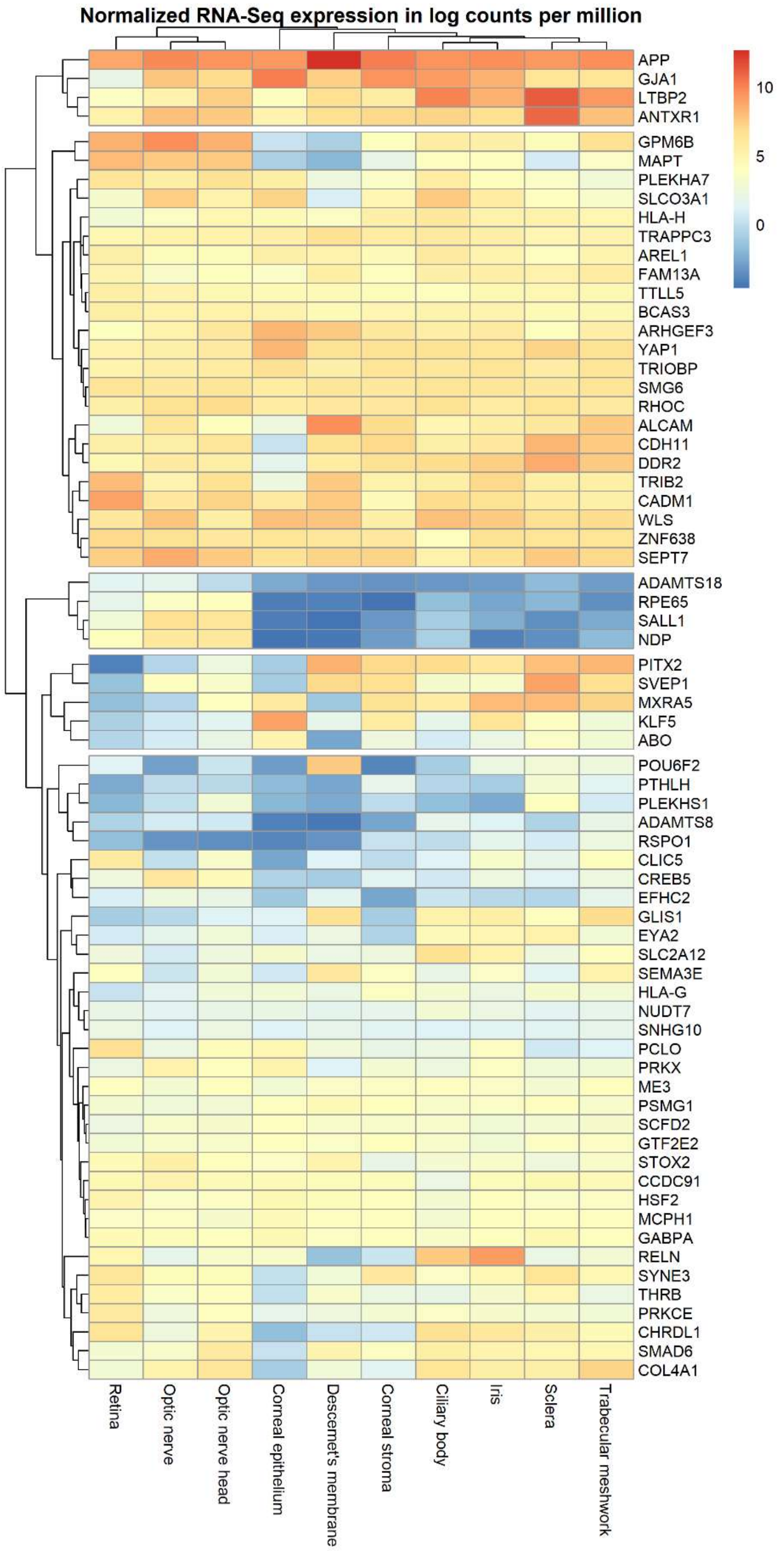

**B) Differential gene expression of the novel POAG risk genes.**

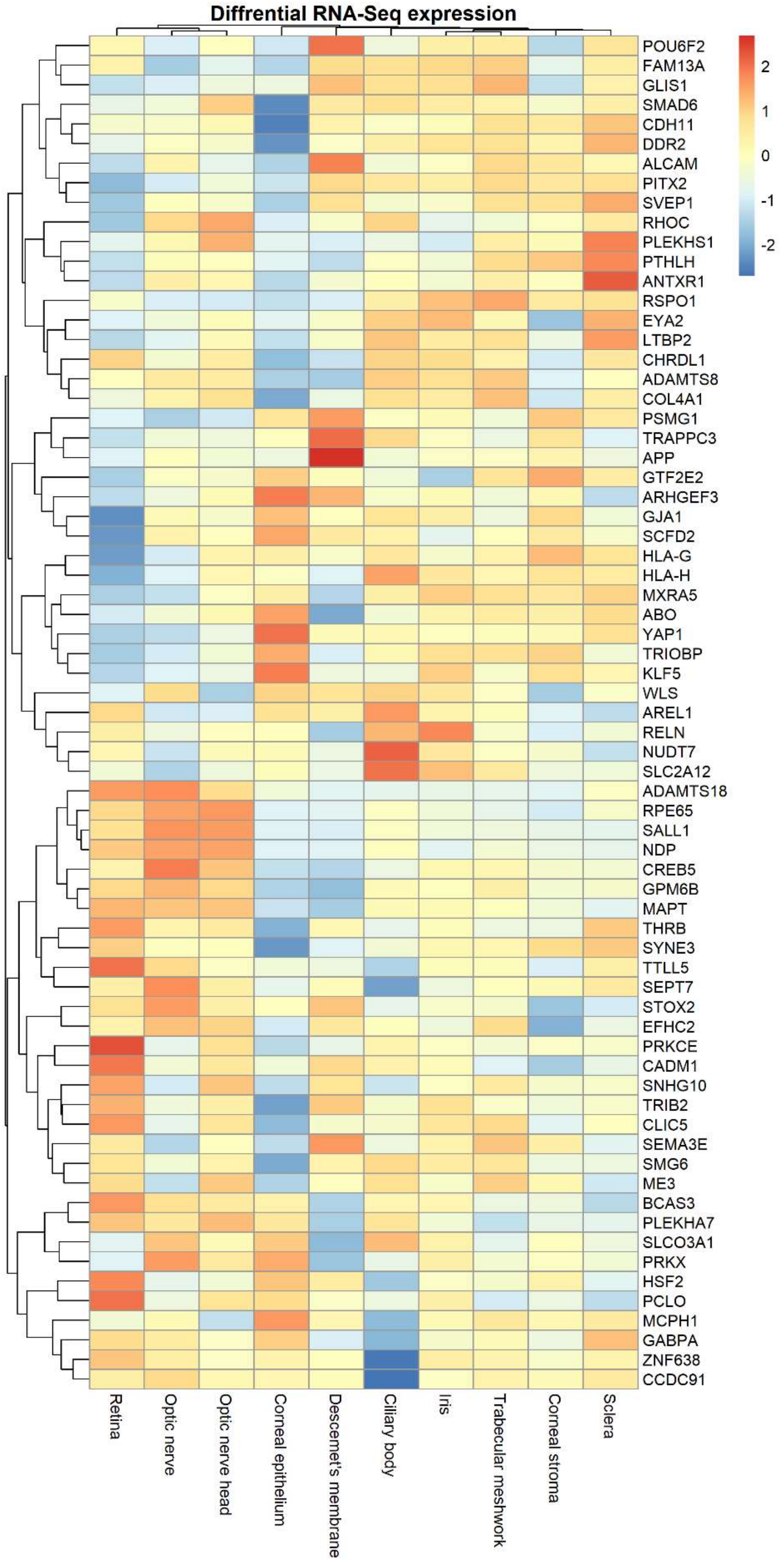
