## Supplementary Notes for "A large cross-ancestry meta-analysis of genome-wide association studies identifies 69 novel risk loci for primary open-angle glaucoma and includes a genetic link with Alzheimer’s disease"

**Study cohorts**

***Australian & New Zealand Registry of Advanced Glaucoma (ANZRAG)***

In total, 3,071 cases and 6,750 unscreened controls of European descent were included from the Australian & New Zealand Registry of Advanced Glaucoma (ANZRAG)^1^. This dataset involves three phases of open-angle glaucoma (OAG) data collection. The first phase was previously published and comprises 1,155 advanced OAG cases and 1,992 controls genotyped on Illumina Omni1M or OmniExpress arrays (Illumina, San Diego, California, USA)^2^. The second phase includes a further 579 advanced OAG cases genotyped on Illumina HumanCoreExome array and 946 controls selected from parents of twins previously genotyped on the same array. The third phase comprises 1,337 OAG cases (11 advanced, 741 non-advanced, and 585 cases with visual field data unavailable) genotyped on Illumina HumanCoreExome array and 3,812 controls selected from a study of endometriosis previously genotyped on the same array. The diagnostic criteria have been described previously^2^. The Approval was obtained from the Human Research Ethics Committees of Southern Adelaide Health Service/Flinders University, University of Tasmania, QIMR Berghofer Medical Research Institute and the Royal Victorian Eye and Ear Hospital. Written informed consent was obtained from all participants. All the methods were carried out in accordance with relevant guidelines and regulations for human subject research, in accordance with the Declaration of Helsinki.

We used the same QC protocol for all the three phases of the ANZRAG OAG GWAS. Briefly, we performed QC using PLINK 1.9^3,4^ by removing individuals with more than 3% missing genotypes, and SNPs with call rate less than 97%, minor allele frequency (MAF) < 0.01, and Hardy-Weinberg equilibrium P < 0.0001 in controls and P < 5 × 10−10 in cases. The same QC protocol was used for case and control datasets before merging to avoid mismatches between the merged datasets. We used PLINK1.9 to compute identity by descent based on autosomal markers, with one of each pair of individuals with relatedness of greater than 0.2 removed within each phase of the ANZRAG data as well as between the three phases. PLINK 1.9 was used to compute principal components for all participants and reference samples of known northern European ancestry (1000 Genomes British, CEU, Finland participants). Participants with PC1 or PC2 values > 6 standard deviations from the mean of known northern European ancestry group were excluded.

Phasing of the genotyped SNPs was conducted using ShapeIT^5^ and imputation was performed using Minimac3 through the Michigan Imputation Server^6^, with the Haplotype Reference Consortium (HRC)^7^ r1.1 as the reference panel. SNPs with imputation quality (r2) > 0.3 and MAF > 0.001 were carried forward for analysis.

We assessed associations between SNPs and OAG status adjusted for sex and the first six principal components under an additive genetic model using the dosage scores obtained from imputation. Association analysis was performed either using SNPTEST v2.5^8,9^ or PLINK 1.9. The same approach was used to investigate the association of SNPs with NTG and HTG subsets within the ANZRAG dataset (821 NTG cases, 1,544 HTG cases, and 6,750 controls).

***NEIGHBOR/MEEI/NHS/HPFS***

All cases and controls met the clinical criteria used previously by the NEIGHBOR and GLAUGEN studies^10,11^. This study was approved by the Massachusetts Eye and Ear Infirmary (MEEI) institutional review board and all subjects signed consent forms approved by the local IRB prior to enrolling in the study. Briefly, POAG cases were defined as individuals for whom reliable visual field (VF) tests showed characteristic VF defects consistent with glaucomatous optic neuropathy. Individuals were classified as affected if the VF defects were reproduced on a subsequent test or if a single qualifying VF was accompanied by a cup-disc ratio (CDR) of 0.7 or more in at least one eye. The majority of cases (over 90%) met this definition, including 96% of the NEIGHBOR cases^11^; and all of the MEEI, NHS, HPFS, and WGHS cases. A small percentage (less than 10%) of the NEIGHBOR cases were defined by cup-to-disc ratio only because visual field data was not available, in some cases because of advanced disease (poor visual acuity) or other medical condition. The CDR definition was > 0.7 in both eyes or CDR asymmetry between the two eyes of 0.2. Patients with signs of secondary causes for elevated IOP such as exfoliation syndrome or pigment dispersion syndrome or critically narrow filtration structures were excluded. The controls were selected to be representative of the age range and gender of the cases. Controls had IOP < 21 mmHg, as measured in a clinical setting, CDR of less than 0.6 and did not have a family history of glaucoma.

***MEEI***

A total of 829 samples that were previously collected and genotyped as part of the GLAUGEN study on the Illumina 660W_Quad_v1 array were included in this analysis^12^. QC was previously performed (SNP call rates >98%). All individuals had genotyping call rate >99%, thus 485 POAG cases and 344 controls were analyzed at 495,160 SNPs. Principal components analysis was run using Eigenstrat and Eigenvectors were tested for association with disease status using a logistic regression model in Stata; Eigenvector 6 was significant. Imputation was performed using Minimac3 through the Michigan Imputation Server^6^, with the Haplotype Reference Consortium (HRC)^7^ r1.1 as the reference panel. SNPs with imputation quality (r2) > 0.3 and MAF > 0.001 were carried forward for analysis.

***NEIGHBOR***

A total of 4,422 samples that were previously collected and genotyped as part of the NEIGHBOR study on the Illumina 660W_Quad_v1 array^10^ were included in this analysis. QC was previously performed (SNP and individual call rates >97%). Pairs of individuals with kinship coefficient >0.0312 were evaluated and the sample with lower genotyping efficiency from each pair removed. This resulted in 4383 individuals (2121 cases, 2262 controls) with genotype calls for 521,687 SNPs. Principal components analysis was run using Eigenstrat and Eigenvectors were tested for association with disease status using a logistic regression model in Stata; Eigenvectors 1,2,3, and 10 were significant. Imputation was performed using Minimac3 through the Michigan Imputation Server^6^, with the Haplotype Reference Consortium (HRC)^7^ r1.1 as the reference panel. SNPs with imputation quality (r2) > 0.3 and MAF > 0.001 were carried forward for analysis.

**NHS/HPFS**

POAG cases in NHS and HPFS have been described elsewhere^13^. This study was approved by the Partners institutional review board and the Harvard School of Public Health Institutional review board. Briefly participants completed surveys every two years that included a question about receiving an eye care provider diagnosis of glaucoma. We retrieved the ophthalmic records from the diagnosing physician and reviewed them in a systematic fashion for POAG. We only included cases without evidence of secondary cause for elevated IOP, open filtration apparatus and reproducible visual field loss on reliable tests consistent with optic nerve disease that could not be explained by other causes.

Many of the participants in these cohorts have been genotyped as part of GWAS for particular diseases. POAG cases in particular were genotyped as part of The Gene, Environment Association Studies (GENEVA) Consortium under the acronym GLAUGEN (Glaucoma Gene and Environment) ^14^. To maximize power, GWAS for different diseases were combined and imputed in a systematic manner to form a combined POAG case control set referred to in this study as NHS/HPFS. In order to decrease false positives in imputation, we used only SNPs that overlap when combining all the datasets. Since different platforms vary in SNP composition, we created two combined datasets based on the type of genotype platform: Affymetrix 6.0, and Illumina Hapmap Series. The details of combining and imputing the data have been previously published^15^. The Affymetrix dataset is comprised of four GWAS: NHS T2D (type 2 diabetes), HPFS T2D, NHS CHD (coronary heart disease), and HPFS CHD. The Illumina HapMap dataset is comprised of 12 datasets: NHS breast cancer, HPFS pancreatic cancer, NHS pancreatic cancer, NHS kidney stone, HPFS kidney stone, NHS2 kidney stone, NHS2 breast cancer, HPFS prostate cancer, NHS glaucoma, and HPFS glaucoma.

POAG cases were identified in both the Affymetrix and Illumina datasaets (incident cases from 1986 – 2010). We implemented several exclusion criteria: we removed all GWAS cancer cases (e.g., breast cancer GWAS cases), and all POAG cases diagnosed with cancer prior to their POAG diagnosis and all controls with prevalent cancer; we removed all controls without an eye exam in 2004-2010, non-Caucasians, and non-confirmed POAG cases (self-reports of glaucoma that were not confirmed by medical record review). The Affymetrix dataset is referred to as NHS/HPFS Affymetrix and the Illumina dataset is referred to as NHS/HPFS Illumina in this study.

We used KING^16^ to calculate kinship coefficients for each pair of individuals both within the Affymetrix dataset, within the Illumina dataset and across the Affymetrix-Illumina datasets, using only the SNPs that overlapped between the two datasets (126,959 SNPs). Unresolved pairs with coefficient>0.0312 were removed (28 controls, 14 from Illumina, 14 from Affymetrix). After filtering, 80 cases and 3,289 controls were included in the Affymetrix dataset, and 382 cases and 2,219 controls in the Illumina dataset. Imputation was performed using Minimac3 through the Michigan Imputation Server^6^, with the Haplotype Reference Consortium (HRC)^7^ r1.1 as the reference panel. SNPs with imputation quality (r2) > 0.3 and MAF > 0.001 were carried forward for analysis.

We calculated PCs with EIGENSTRAT, using all of the SNPs available in the imputed dataset. We checked the top 10 PCs for association with POAG and included only PCs with p<0.05 in the analysis. In both datasets, we controlled for GWAS study (which inherently controlled for gender since each study only contained individuals of one gender or the other), and age at baseline. Since half of the GWAS studies contributing to the Affymetrix NHS/HPFS dataset were studies of type 2 diabetes, the Affymetrix dataset was enriched with diabetes cases. As there is evidence for an association between POAG and T2D^17^, we controlled for diabetes status in the Affymetrix dataset. Association testing was performed in a logistic regression model, adjusting for age, BMI, diabetes status, and the top four PCs.

***European Prospective Investigation into Cancer-Norfolk Eye Study (EPIC-Norfolk Eye Study)***

The European Prospective Investigation into Cancer (EPIC) study is a pan-European prospective cohort study designed to investigate the etiology of major chronic diseases^18^. EPIC-Norfolk , one of the UK arms of EPIC, recruited and examined 25,639 participants between 1993 and 1997 for the baseline examination^19^. Recruitment was via general practices in the city of Norwich and the surrounding small towns and rural areas, and methods have been described in detail previously^20^. Since virtually all residents in the UK are registered with a general practitioner through the National Health Service, general practice lists serve as population registers. Ophthalmic assessment formed part of the third health examination and this has been termed the EPIC-Norfolk Eye Study^21^. In total, 8,623 participants were seen for the Eye Study, between 2004 and 2011. Ophthalmic examination included tonometry (Ocular Response Analyzer; Reichert, New York, USA; software V.3.01), optic disc photography (Nikon D80 camera; Nikon Corporation, Tokyo, Japan), scanning laser ophthalmoscopy (Heidelberg Retinal Tomograph 3; Heidelberg Engineering, Heidelberg, Germany) and nerve fiber layer assessment (Gdx-VCC; Zeiss, Dublin, California, USA). Participants meeting pre-defined criteria and an additional 1:10 participants underwent automated visual field testing (Humphrey 750i Visual Field Analyzer; Carl Zeiss Meditech Ltd, Welwyn Garden City, UK). The EPIC-Norfolk Eye Study was carried out following the principles of the Declaration of Helsinki and the Research Governance Framework for Health and Social Care. The study was approved by the Norfolk Local Research Ethics Committee (05/Q0101/191) and East Norfolk & Waveney NHS Research Governance Committee (2005EC07L). All participants gave written, informed consent.

Ascertainment of POAG in the EPIC Norfolk third health examination has been described previously^22^. Participants with study results suspicious of glaucoma (using pre-defined criteria) were referred for further examination by a glaucoma specialist at the regional University Hospital^21^. Additionally, a diagnosis refinement process was undertaken by a second glaucoma specialist who independently reviewed the test results of all participants classified as glaucoma and a proportion of participants who were not classified as having glaucoma. POAG was defined as the presence of a glaucomatous optic disc together with either a corresponding visual field defect or otherwise unexplained non-specific visual field loss, open angles on gonioscopy, and absence of secondary causes of glaucoma. A glaucomatous disc was defined as one with focal or diffuse neuro-retinal rim thinning, and may possess, though not necessary for the definition, additional characteristic features such as bared circumlinear vessels, disc haemorrhages or nerve fiber layer defects. Pseudoexfoliative and pigmentary glaucoma were defined as secondary glaucoma in this study and therefore did not contribute to POAG cases. In addition to the POAG cases identified among the 8,623 participants attending the 3rd health examination, we included a further 397 POAG cases from the remainder of the cohort via linkage with hospital episode statistic data (ICD10 code H40.1). We defined controls as participants not meeting referral criteria for glaucoma on initial ophthalmic assessment and participants who attended the University Hospital for further examination and were not classified as having or being suspect for any type of glaucoma or ocular hypertension. Additionally, we required controls to have intraocular pressure ≤ 21 mmHg and HRT vertical cup-to-disc ratio ≤ 0.5 to minimise the chance of false negatives.

Genotyping was undertaken using the Affymetrix UK Biobank Axiom Array. SNP exclusion criteria included: call rate < 95%, abnormal cluster pattern on visual inspection, plate batch effect evident by significant variation in minor allele frequency, and/or Hardy-Weinberg equilibrium P < 10-7. Sample exclusion criteria included: DishQC < 0.82 (poor fluorescence signal contrast), sex discordance, sample call rate < 97%, heterozygosity outliers (calculated separately for SNPs with minor allele frequency >1% and <1%), rare allele count outlier, and impossible identity-by-descent values. Following these exclusions, there were no ethnic outliers. After filtering, 664 cases and 5,630 controls were used in this study. Imputation was carried out using the Haplotype reference Consortium as a reference panel. Associations between the selected SNPs and POAG were examined using logistic regression adjusted for age, sex and the first 5 principal components, assuming an additive model and using the imputed dosage data. Analyses were carried out using SNPTEST version 2.5.1.

***UKBB POAG ICD10 code and self report cases***

UK Biobank (UKBB) is a large-scale cohort study that included over 500,000 participants aged between 40-69 years in 2006-2010 from across the United Kingdom. The glaucoma cases definition and association analysis were described previously.^23^ In this study, we defined UKBB POAG ICD-10 cases and self-reported glaucoma cases (exclude ICD-10 POAG cases) using updated phenotype data in April 2019. In brief, we identified ICD-10 POAG cases as those who had an ICD-10 diagnosis of ‘primary open angle glaucoma’, ‘other glaucoma’ or ‘glaucoma, unspecified’. We identified self-reported glaucoma cases as those who (1) responded ‘glaucoma’ to the question ‘Has a doctor told you that you have any of the following problems with your eyes?’; or (2) responded ‘glaucoma’ to the question ‘In the touch screen you selected that you have been told by a doctor that you have other serious illnesses or disabilities, could you now tell me what they are? (non-cancer illness). In both ICD-10 POAG cases and self-reported glaucoma cases, we removed primary angle-closure glaucoma (PACG) cases. We then selected controls as those who reported that did not have any eye diseases. We used separate controls for ICD10 POAG cases and self-reported glaucoma cases in UKBB. With the ratio of 1:5 for ICD10 POAG cases and self-reported glaucoma cases, we spit the controls into two groups randomly with a ratio of 1:5. We then removed related participants ($\hat{\pi}$>0.2) within and between ICD-10 POAG GWAS and self-reported glaucoma GWAS. Finally, we included 1,448 ICD-10 POAG cases and 22,107 controls for ICD-10 POAG GWAS in UKBB, and 7,286 self-reported glaucoma cases and 107,362 controls for self-reported glaucoma GWAS. In the GWAS analysis, the dosage scores from imputation were used in the logistic regression models (PLINK version 2.0) adjusted for sex, age, and the first ten principal components.

***Kaiser Permanente GERA Cohort***

The Genetic Epidemiology Research in Adult Health and Aging (GERA) cohort consists of 110,266 adults who consented to participate in the Research Program on Genes, Environment, and Health, established for members of the Kaiser Permanente Medical Care Plan, Northern California Region (KPNC)^24,25^. The Institutional Review Board of the Kaiser Foundation Research Institute has approved all study procedures.

All GERA subjects included in this study had valid IOP measures, as previously described^26^. Briefly, non-numeric entries for IOP, extreme values (≤5 and >60 mm Hg), and measurements taken on a single eye were removed. Further, IOP measurements that were taken after initial prescription of IOP-lowering medications were excluded to avoid values influenced by treatment. Because IOP-lowering medications are almost always prescribed before IOP-lowering surgical interventions (i.e., laser trabeculoplasty, trabeculectomy, tube shunt procedures, etc.), we did not remove subjects who had these surgical interventions. Patients eligible for inclusion were identified from clinical diagnoses captured in the KPNC electronic health record (EHR) system. These clinical diagnoses were recorded in the EHR system as International Classification of Diseases, Ninth Revision (ICD-9) diagnosis codes. We defined glaucoma cases as having at least: (1) two diagnoses of POAG (ICD-9 codes 365.01, 365.1, 365.10, 365.11, and 365.15); or (2) two diagnoses of NTG (ICD-9 code 365.12); or (3) one diagnosis of POAG and one diagnosis of NTG. In all cases, at least one of the diagnoses was made by a Kaiser Permanente ophthalmologist. Further, the cases did not have any diagnosis of other subtypes of glaucoma (e.g., pseudoexfoliation, pigmentary, or PACG; ICD-9 codes 365.52, 365.13, and 365.2, respectively). After excluding subjects who have one or more diagnosis of any type of glaucoma (ICD-9 code, 365.xx), our control group included all the non-cases. Subjects who had no diagnosis of any type of glaucoma (any ICD-9 code 365.xx other than 365.04) but did have a diagnosis of OHTN (ICD-9 code 365.04), were included as controls. In total, 4,258 POAG and 51,986 controls from GERA (non-Hispanic white and East Asian ethnic groups) were included in this study.

GERA individuals’ DNA samples were extracted using Oragene kits (DNA Genotek Inc., Ottawa, ON, Canada) at KPNC and genotyped at the Genomics Core Facility of UCSF. DNA samples were genotyped at over 665 000 genetic markers on four race/ethnicity-specific Affymetrix Axiom arrays (Affymetrix, Santa Clara, CA, USA) optimized for European, Latino, East Asian, and African-American individuals^27,28^. We performed genotype quality control (QC) procedures for the GERA samples on an array-wise basis^25^. Briefly, we included genetic markers with initial genotyping call rate ≥ 97%, genotype concordance rate > 0.75 across duplicate samples, and allele frequency difference ≤ 0.15 between females and males for autosomal markers. Approximately 94% of samples and over 98% of genetic markers assayed reached QC procedures. Moreover, genetic markers with genotype call rates < 90% were excluded, as well as genetic markers with a MAF < 1%. Following the pre-phasing of genotypes with Shape-IT v2.r72719^5^, imputation was performed using Minimac3 through the Michigan Imputation Server^6^, with the Haplotype Reference Consortium (HRC)^7^ r1.1 as the reference panel. SNPs with imputation quality (r2) > 0.3 and MAF > 0.001 were carried forward for analysis.

***King’s College London (KCL)***

The POAG case–control cohort consists of 576 cases and 287 controls of European ancestry and 298 cases and 194 controls of West African ancestry residing in the United Kingdom. European controls, high-tension glaucoma cases, and a small number of normal-tension cases were recruited in South London at St Thomas' Hospital and the Princess Royal University Hospital, UK. The majority of European normal-tension cases were patients at the University Eye Hospitals in Tuebingen and Wuerzburg, Germany. All African cases, and some of the controls were recruited in South London at St Thomas' Hospital, additional controls were used from the SOUL-D study (The South London Diabetes Study). Patients were included in this study if they had visual field loss in at least one eye attributed to glaucoma by a glaucoma specialist, had a VCDR of more than 0.6, were receiving intraocular-lowering medication (or had previous surgery), and had open drainage angles on gonioscopy. European controls were also examined by an ophthalmologist and were confirmed to be either free from any eye disease, or only had cataracts with no evidence of glaucoma but their visual field was not tested. African controls from St Thomas' followed the same clinical screening, while controls from the SOUL-D study were not screened but did not report glaucoma. Diagnostic criteria for HTG and NTG included a presence of both a VCDR>=0.6 and a visual field defect. In addition to this, HTG cases had a highest recorded, pre-treatment IOP(GAT)>21mmHg, and NTG cases had a highest recorded, pre-treatment IOP(GAT)<21mmHg. All subjects were genotyped in the same batch on the Illumina Human Omni Express Exome 8v1-2 chip. The African cohort was confirmed to be of West-African descent through principal component analysis with samples from the 1000G Nigerian, Gambian and Sierra Leonne populations. Imputation was performed using the Michigan Imputation Server. European and African subjects were imputed separately. The HRC reference panel was used for Europeans and the 1000G phase 3 panel was used for Africans. SNPs were filtered using a minor allele frequency of >0.001 and imputation quality score >0.3. GWAS was conducted in Plink 2 using gene dosage in an additive model, logistic regression (Firth). Age, sex and the first 6 principal components were used as covariates (sex was excluded as a covariate for the sex-stratified analyses).

***Blue Mountain Eye Study (BMES)***

The BMES is a population-based eye assessment of a representative older Australian community sample. During the period 1992-1994, 3654 residents (57% male and 43% females) aged 49-97 were examined (BMES-1). The study was approved by the Western Sydney Area Health Service Human Ethics Committee, and written informed consent was obtained from all participants. Open-angle glaucoma was diagnosed when glaucomatous defects on the Humphrey 30-2 test matched the optic disc changes, without regard to the intraocular pressure level.

The BMES cohort included 107 BMES glaucoma cases and 600 population controls (from BMES) genotyped on Illumina Human 610 Quad Array and imputed against HRC reference panel. The methods used for imputation and statistical analysis of this cohort were the same as the ANZRAG cohort. After quality control (QC), 540,035 SNPs were used as the basis of imputation for the BMES cohort.

***Southampton***

Primary open-angle (POAG) and normal tension glaucoma patients were recruited from the Southampton University Hospital Trust Eye Clinic and satellite regional glaucoma clinics. Ethical approval for the collection of patient information and blood samples was provided by the Southampton and South West Hampshire Local Research Ethics Committee (05/Q1702/8)and Cohort Recruitment commenced in August 2005.Each patient was examined by an experienced glaucoma specialist. Diagnoses were made on the basis of characteristic visual field loss/glaucomatous optic disc damage/increased IOP. Patients presenting with narrow-angle, developmental or secondary glaucoma or any other known abnormalities of the anterior segment were excluded. Patients with unambiguous glaucoma, but normal tension were included in sample collection later. Furthermore, to select for patients with typical POAG or normal-tension glaucoma (NTG), only patients diagnosed over the age of 40 years were included. Both conditions are rare before this age. DNA was extracted according to the standard methods, dissolved in TE buffer, and stored at-20°C.Primary open angle glaucoma patients were genotyped on the AffymetrixSNP6.0 array, all data were exported on the forward strand. These data were combined with the Affymetrix SNP 6.0 data publically available for the WTCCC2 controls. The Genome-wide association data was further filtered to ensure removal of individuals with more than 5% missing genotypes, SNPs with more than 3%missing samples, SNPs with a minor allele frequency of <1% and a Hardy-Weinberg p-value <1x10^-6^.SNPs were all on the forward strand and locations were lifted over from hg18 to hg19 using the UCSC liftover tool. SNPs with complementary alleles were also excluded (A/T and G/C).

The data used for imputation included 941 cases and 1,557 controls, and 533,774 SNPs. Pre-phasing was carried out using Shapeit (v2, r790)and imputation was carried out using Minimac3 through the Michigan Imputation Server^6^, with the Haplotype Reference Consortium (HRC)^7^ r1.1 as the reference panel. SNPs with imputation quality (r2) > 0.3 and MAF > 0.001 were carried forward for analysis. Case-control analysis was carried out using logistic regressions for the selected replication SNPs and Indels with sex as a covariate, using PLINK (v1.90b3b 64-bit (15 Jan2015)).

**Gutenberg Health Study (GHS)**

The GHS is a population-based, prospective, observational cohort study in the Rhine-Main Region in midwestern Germany with a total of 15,010 participants and follow-up after five years. The study sample is recruited from subjects aged between 35 and 74 years at the time of the exam. The sample was drawn randomly from local governmental registry offices and stratified by gender, residence (urban and rural) and decade of age. Exclusion criteria were insufficient knowledge of the German language to understand explanations and instructions, and physical or psychic inability to participate in the examinations in the study center. The study was approved by the Medical Ethics Committee of the University Medical Center Mainz and by the local and federal data safety commissioners. Genotyping was carried out using Affymetrix Genome‐Wide Human SNP 6.0 Array. Imputation was performed using Minimac3 through the Michigan Imputation Server^6^, with the Haplotype Reference Consortium (HRC)^7^ r1.1 as the reference panel. SNPs with imputation quality (r2) > 0.3 and MAF > 0.001 were carried forward for analysis. After QC, 47 cases and 2,731 controls were included in this study. POAG cases were defined based on the ISGEO classification. Association testing was performed using Plink2, assuming an additive genetic model with age, sex, and the top three PCs fitted as covariate.

***Erasmus Rucphen Family (ERF) study***

The ERF study is a family‐based cohort in a genetically isolated population in the southwest of the Netherlands with over 3,000 participants aged between 18 and 86 years^29,30^. In the region of the ERF population, a total of 110 patients with glaucoma who did not participate in the ERF study were recruited in three local hospitals. Their visual fields were tested with standard automated perimetry (Humphrey Field Analyzer c24-2 SITA Standard test program) or the Octopus 101 (G2 program with TOP strategy) (Haag-Streit, Bern, Switzerland). The diagnosis of glaucoma was made by the patient’s ophthalmologist and confirmed by a glaucoma specialist (HGL). It was based on a glaucomatous appearance of the optic disc (notching or thinning of the neuroretinal rim), combined with a matching glaucomatous visual field defect, and open-angles seen by gonioscopy. Participants from the ERF study were used as control group (n = 1999). Genotyping was performed with the 318K array of the Illumina Infinium II whole-genome genotyping assay (HumanHap300-2). Samples with low call rate (<97.5%), with excess autosomal heterozygosity (>0.336), or with sex‐mismatch were excluded. A set of genotyped input SNPs with call rate >98%, with MAF >0.01, and with HWE p-value >10−6 was used for imputation. Imputation with HRC was facilitated by the Michigan Imputation server, file preparation was done using scripts provided online (HRC Imputation preparation and checking: http://www.well.ox.ac.uk/~wrayner/tools/; v4.2.1). Filtered genotypes were uploaded. The server uses SHAPEIT2 (v2.r790) to phase the data and Minimac 3 for imputation to the HRC reference panel (v1.0). We used the imputed dosages returned by the service. Association analyses were performed in RvTest *--meta score* option; to adjust for familial relationships in ERF, the kinship matrix estimated from the genotyped data was used, analyses were also adjusted for age and sex.

***Rotterdam Study I***

The Rotterdam Study is a population-based study established in Rotterdam, the Netherlands^31^. It consists of three cohorts. The original cohort, Rotterdam Study I (RS-I), started in 1990 and includes 7,983 subjects aged 55 years and older. Participants visiting the research center underwent an extensive ophthalmic examination. Details of the eye examinations have been described elsewhere. POAG was defined as glaucomatous visual field loss (GVFL) with glaucomatous optic nerve neuropathy (GON)^32^. DNA was isolated from whole blood according to standard procedures. Genotyping of SNPs was performed using the Illumina Infinium II HumanHap550 array (RS-I). Samples with low call rate (<97.5%), with excess autosomal heterozygosity (>0.336), or with sex-mismatch were excluded, as were outliers identified by the identity-by-state clustering analysis (outliers were defined as being >3 standard deviation (s.d.) from population mean or having identity-by-state probabilities >97%). A set of genotyped input SNPs with call rate >98%, MAF >0.001 and Hardy-Weinberg Equilibrium (HWE) p-value >10-6 was used for imputation. Imputation with HRC was facilitated by the Michigan Imputation server, file preparation was done using scripts provided online (HRC Imputation preparation and checking: http://www.well.ox.ac.uk/~wrayner/tools/; v4.2.1). Filtered genotypes were uploaded. The server uses SHAPEIT2 (v2.r790) to phase the data and Minimac 3 for imputation to the HRC reference panel (v1.0). We used the imputed dosages returned by the service. GWAS analyses were performed using Rvtest *--meta score* option. The analyses were adjusted for age, sex, and the first five principal components. The Rotterdam Study has been approved by the institutional review board (Medical Ethics Committee) of the Erasmus Medical Center and by the review board of The Netherlands Ministry of Health, Welfare and Sports.

***Geisinger***

Geisinger cohort consists of a sample size of 43,982 who consented to participate in the MyCode Community Health Initiative program. Within this dataset, the POAG case-control group consists of 664 cases and 5904 controls of European ancestry. All samples were genotyped on Illumina Human Omni Express genotyping platform and imputation was carried out using Michigan Imputation Server with HRC reference panel. Markers with imputation quality (r2) >= 0.3 and MAF >= 0.01 were carried forward for analysis. Additionally a HWE p-value < 1e-07, a marker call rate of 99% and a sample call rate of 90% were applied as a further filtration criteria. The analysis was run in unrelated samples who are 18 years or older and was adjusted for the covariates of first four principal components and age.

***The Chinese University of Hong Kong (CUHK)***

Patients with POAG were defined using the same criteria as described for the Singaporean POAG collection. All subjects of the Hong Kong study population were recruited from the Chinese University of Hong Kong after ethics committee approved the study protocol following the tenets of the Declaration of Helsinki. A total of 217 POAG patients of Han Chinese ancestry and 397 controls of Hong Kong and Guangdong Chinese of Han ancestry were used in this study. The Hong Kong control subjects were recruited in a hospital-based manner. They were all given complete ocular examinations, and confirmed to have no sign of glaucoma, angle closure or narrow angle, or other major eye diseases except for mild cataract and mild refractive errors. Control subjects were recruited from elderly people aged ≥ 60 years to ensure they were at least free of early-onset major eye diseases. They had IOP < 21 mmHg, and had no known family history of glaucoma. Additional healthy controls were recruited from local communities in the Guangdong province of Southern China^33^. Genotyping was carried out using Illumina Humancnv370-quad V3.0. Imputation was performed using Minimac3 and 1000G phase 3 as the reference panel. SNPs with imputation quality (r2) > 0.3 and MAF > 0.001 were carried forward for analysis. Association testing was performed in a logistic regression model with age, sex, and the top six PCs fitted as covariate.

***Singapore Chinese***

POAG including NTG patients were from the Singapore National Eye Center glaucoma clinics. Ethical approval for the collection of patient information and blood samples was provided by the local institutional ethics review committee (CIRB) and study was conducted in accordance with revised Declaration of Helsinki. Patients with POAG over the age of 40 years old were recruited. Patients with POAG were defined by the following criteria: the presence of glaucomatous optic neuropathy (defined as a loss of neuroretinal rim with a vertical cup: disc ratio of >0.7 or an inter-eye asymmetry of >0.2, and/ or notching attributable to glaucoma) with compatible visual field loss, open angles on gonioscopy, and absence of secondary causes of glaucomatous optic neuropathy. POAG patients with a mean IOP without treatment that was consistently <21 mm Hg on diurnal testing were classified as having NTG, whereas those with a mean IOP without treatment that was consistently >21 mm Hg on diurnal testing were classified as having HTG. Diurnal IOP measurements were recorded hourly between 8.00 am to 5.00 pm with non-contact air puff tonometry (Topcon CT-80 Computerized Non-Contact Tonometer, Topcon). Patients were excluded if they were unable to give written informed consent or had either neurological/retinal disease that had visual field sequelae, or secondary glaucoma (such as pigmentary, uveitic, neovascular or post-traumatic).

1224 Singaporean Chinese POAG patients were recruited under the above criteria, and their genomic DNA samples were genotyped using the Illumina Human OmniExpress Beadchips. 1033 cases passed quality control filters and were brought forward for downstream analyses. The final sample cohort with clear sub-diagnosis comprised of 289 NTG and 709 HTG subjects. Controls were ascertained from an on-going population based study of Chinese persons aged 40 years and older (the Singapore Chinese Eye Study [SCES])^34^. The SCES is a population-based, cross-sectional study of Chinese adults residing in the south- western part of Singapore. 2587 SCES samples were genotyped with Illumina Human OmniExpress or Human610-Quad BeadChips. 2371 that passed quality check were used in subsequent genetic analyses. The imputation and phasing of genotypes were carried out using IMPUTE2 (http://mathgen.stats.ox.ac.uk/impute/impute_v2.html) with cosmopolitan population haplotypes based on data from 2535 individuals from 26 distinct populations around the world obtained from the 1000 Genomes project Phase 3 (Jun 2014) release for reference panel construction. SNPs with imputation quality (r2) > 0.3 and MAF > 0.001 were carried forward for analysis.

***Japanese Tohoku***

All the POAG patients were recruited at the Institutes related to Tohoku University. The study protocol followed the tenets of the Declaration of Helsinki and was approved by the Institutional Review Board of the Tohoku Graduate School of Medicine. All participants in this study were at the age of 35 years or older and of Japanese residents. All of the subjects with POAG were diagnosed by glaucoma specialists and fulfilled the following diagnostic criteria: presence of glaucomatous optic disk changes, including neuroretinal rim thinning, notching, or cupping; presence of visual field defect that could be attributed to the optic disk changes; and no history of secondary, angle closure, or congenital glaucoma. The control subjects recruited at the Tohoku Medical Megabank Organization as a part of prospective cohort study were considered to have no POAG based on self-report.

A total of 1267 patients with POAG and 2517 healthy controls were recruited above the criteria. DNA samples from the participants were collected from whole blood according to standard procedures. Japonica array^TM^ (Toshiba, Tokyo, Japan) is a custom-designed array optimized for the Japanese population based on the information from the reference panel from 1,070 Japanese. Genome-wide genotype data set was obtained using this SNP array for 1267 patients with POAG according to the manufacturer’s instructions. 38 case samples that did not satisfy the quality control criteria (DishQC > 0.82 and call rates > 0.970) and 38 case samples from pairs with a relatedness measure (pi-hat) value >0.2 were excluded, thus resulting in a dataset comprising 1191 cases. As a control, genotype data from 2434 healthy subjects collected previously through a prospective cohort study were used.

SNP quality control was applied for the imputation procedure. We used info score using IMPUTE2, and considered a SNP with info score great than 0.9 as an acceptable well-imputed variant in this study. Autosomal SNPs that were assigned to ‘Recommended’ by the Ps_Classification program in the SNPolisher package (Affymetrix) were selected. We applied the following thresholds for quality control in further data cleaning: Hardy–Weinberg equilibrium with a P value <0.001 for control samples, call rate for each SNP > 0.970, and minor allele frequencies <0.005. As a result, a total of 608329 SNPs on autosomal chromosomes passed the quality control filters and were used for whole genome imputation. Prephasing was first conducted with these SNPs by SHAPEIT (v2.r837). Genotype imputation was performed on the phased genotypes with IMPUTE2 (ver. 2.2.2) by using a phased reference panel of the 1000 Genomes project Phase3. SNPs with imputation quality (r2) > 0.3 and MAF > 0.001 were carried forward for analysis.

***Biobank Japan***

The Biobank Japan (BBJ; <http://biobankjp.org>) is a large, patient-based biobank which started at the Institute of Medical Science, the University of Tokyo in 2003, consisted of approximately 267,000 individuals with 51 target diseases including glaucoma. The definition of POAG, NTG, and HTG are fully described in our previous reports^35,36^. As the controls of present study, we randomly selected 30,000 samples from the participants of the BBJ using ‘--thin-indiv-count’ option implemented by PLINK1.9, and integrated them with genotype data of general Japanese subjects obtained from Pharma SNP consortium (N=946). After the quality controls, 3,979 cases and 30,278 controls were included in the association analysis. All of these participants were genotyped by (a) a combination of Illumina Human OmniExpress BeadChip and Infinium HumanExome BeadChip or (b) Infinium OmniExpressExome. We phased genotypes using Eagle2, and imputed using 1000 Genomes Project Phase 3 (N = 2,504) using minimac3 as we described elsewhere^37,38^. Association analysis was performed in logistic regression model by PLINK2 using age, sex, top six PCs as covariates.

***GIGA (TZ, SAC, SAB)***

The Genetics In Glaucoma patients from African descent study (GIGA) is a multicenter case–control study comprising POAG patients and controls from South Africa and Tanzania. Participants from Black African and South African Coloured ancestry were ascertained from the ophthalmology outpatient department of the Groote Schuur Hospital in Cape Town, South Africa (*N*_cases_ = 327; *N*_controls_= 194), and from hospitals in Tanzania: Muhimbili National Hospital and CCBRT Disability Hospital in Dar es Salaam (*N*_cases_ = 395; *N*_controls_ = 382). The study was conducted according to the guidelines for human research by the National Institute for Medical Research in Tanzania. Ethical approval was obtained from the institutional review boards at each study site, and written informed consent was provided by each participant.

POAG cases met category 1 or 2 of the ISGEO classification for open-angle glaucoma^39^. In brief, cases had either a definite visual field defect and Vertical Cup Disc Ratio (VCDR) ≥ 0.7, or VCDR > 0.8 in the absence of a visual field test. Other inclusion criteria were an open angle on gonioscopy and age of onset older than 35 years. Glaucoma patients diagnosed with secondary causes were excluded from this study. Controls were recruited at the same ophthalmology clinics and underwent identical examinations as the POAG cases. Inclusion criteria were: no signs of glaucoma, IOP ≤ 21 mmHg; VCDR< 0.5, and a VCDR inter-eye asymmetry < 0.2, no family history of glaucoma, and age> 55 years. Case and control status was adjudicated by two experienced ophthalmologists.

1162 participants were genotyped using either the Illumina HumanOmniExpressExome Beadchip (964,193 variants; Illumina, Inc., San Diego, CA, USA; *n* = 999) or the Illumina HumanOmni2.5Exome Beadchip (2,406,945 variants; Illumina, Inc., San Diego, CA, USA; *n* = 163). Extensive quality control (QC) was performed on the genotyped data in PLINK v1.07^3^. Variants with a call rate < 95%, as well as variants showing an extreme deviation from Hardy–Weinberg equilibrium (*P* < 1 × 10^−6^), or MAF < 0.01 were excluded. All SNPs were mapped to genome build hg19/GRCh37. Individual level QC was performed by exclusion of individuals with a call rate < 95%, discordant sex in self-report versus genetically determined sex, excess or reduced heterozygosity, relatedness (PI-HAT > 0.25) or duplicative samples based on identity by descent (IBD) sharing calculations. The final dataset consisted of 663 and 476 successfully genotyped POAG cases and controls, respectively.

Imputation of unknown genetic variation was performed by means of the “cosmopolitan” approach of using all available populations in a reference panel. The 1000 Genomes Project phase III version 5 was used as an imputation reference panel for GIGA^40^. The pipeline implemented at the Michigan Imputation Server ([https://imputationserver.sph.umich.edu](https://imputationserver.sph.umich.edu/)) was used for prephasing and imputation (Minimac) of GIGA genotypes^6^.

***Eyes of Africa Genetic Consortium***

The Eyes of Africa Genetic consortium is a Pan-African study of genetic determinants of POAG, and comprises studies recruited from Ghana, Nigeria, South Africa and the USA, totaling a sample size of 2320 POAG cases and 2121 controls. The methods of ascertaining POAG cases has been described in detail elsewhere^41^. In brief, POAG cases met the following inclusion criteria: glaucomatous optic neuropathy (VCDR > 0.7 or notch in the neuroretinal rim), and visual field loss (examined by frequency doubling technology or standard automated perimetry) consistent with optic nerve damage in at least one eye. Controls were participants with no known first-degree relative with glaucoma, IOP less than 21 mmHg in both eyes without treatment, and no evidence of glaucomatous optic neuropathy in either eye. Genotyping of cases and controls was performed on the Illumina OmniExpressExome array. Imputation was carried out using IMPUTE2 and 1000G phase1 version 3 as the reference panel. SNPs with imputation quality (r2) > 0.3 and MAF > 0.001 were carried forward for analysis.

#### 23andMe Research Team

#### Michelle Agee, Stella Aslibekyan, Robert K. Bell, Katarzyna Bryc, Sarah K. Clark, Sarah L. Elson, Kipper Fletez-Brant, Pierre Fontanillas, Nicholas A. Furlotte, Pooja M. Gandhi, Karl Heilbron, Barry Hicks, David A. Hinds, Karen E. Huber, Ethan M. Jewett, Yunxuan Jiang, Aaron Kleinman, Keng-Han Lin, Nadia K. Litterman, Jennifer C. McCreight, Matthew H. McIntyre, Kimberly F. McManus, Joanna L. Mountain, Sahar V. Mozaffari, Priyanka Nandakumar, Elizabeth S. Noblin, Carrie A.M. Northover, Jared O'Connell, Steven J. Pitts, G. David Poznik, J. Fah Sathirapongsasuti, Anjali J. Shastri, Janie F. Shelton, Suyash Shringarpure, Chao Tian, Joyce Y. Tung, Robert J. Tunney, Vladimir Vacic, Amir S. Zare

### Eye and Vision Consortium members

Sarah BARMAN, Jenny BARRETT, Paul BISHOP, Peter BLOWS, Catey BUNCE, Roxana CARARE, Usha CHAKRAVARTHY, Michelle CHAN, Antonietta CHIANCA, Valentina CIPRIANI, David CRABB, Philippa CUMBERLAND, Alexander DAY, Parul DESAI, Bal DHILLON, Andrew DICK, Cathy EGAN , Sarah ENNIS, Paul FOSTER, Marcus FRUTTIGER, John GALLACHER, David (Ted) GARWAY-HEATH, Jane GIBSON, Dan GORE, Jeremy GUGGENHEIM, Chris HAMMOND, Alison HARDCASTLE, Simon HARDING, Ruth HOGG, Pirro HYSI, Pearse A KEANE, Sir Peng Tee KHAW, Anthony KHAWAJA, Gerassimos LASCARATOS, Andrew LOTERY, Phil LUTHERT, Tom MACGILLIVRAY, Sarah MACKIE, Keith MARTIN, Michelle MCGAUGHEY, Bernadette MCGUINNESS, Gareth MCKAY, Martin MCKIBBIN, Danny MITRY, Tony MOORE, James MORGAN, Zaynah MUTHY, Eoin O'SULLIVAN, Chris OWEN, Praveen PATEL, Euan PATERSON, Tunde PETO, Axel PETZOLD, Jugnoo RAHI, Alicja RUDNICKA, Jay SELF, Sobha SIVAPRASAD, David STEEL, Irene STRATTON, Nicholas STROUTHIDIS, Cathie SUDLOW, Caroline THAUNG, Dhanes THOMAS, Emanuele TRUCCO, Adnan TUFAIL, Marta UGARTE, Veronique VITART, Stephen VERNON , Ananth VISWANATHAN , Cathy WILLIAMS, Katie WILLIAMS, Jayne WOODSIDE, Max YATES, Jennifer YIP, Yalin ZHENG, Haogang ZHU, Robyn TAPP, Denize ATAN.

**The NEIGHBORHOOD consortium members**

R. Rand Allingham Murray Brilliant, Donald L. Budenz, Jessica Cooke Bailey, John H. Fingert, Douglas Gaasterland, Teresa Gaasterland, Jonathan L Haines, Michael Hauser, Robert P. Igo Jr, Jae Hee Kang, Peter Kraft, Richard K. Lee, Paul R. Lichter, Yutao Liu, Louis R Pasquale, Syoko Moroi, Jonathan Myers, Margaret Pericak-Vance, Anthony Realini, Doug Rhee, Julia E. Richards, Robert Ritch, Joel S. Schuman, William K. Scott, Kuldev Singh, Arthur J. Sit, Douglas Vollrath, Janey L. Wiggs, Gadi Wollstein & Donald J. Zack.

**FinnGen Project members**

Anu Jalanko, Jaakko Kaprio, Kati Donner, Mari Kaunisto, Nina Mars, Alexander Dada, Anastasia Shcherban, Andrea Ganna, Arto Lehisto, Elina Kilpeläinen, Georg Brein, Ghazal Awaisa, Jarmo Harju, Kalle Pärn, Pietro Della Briotta Parolo, Risto Kajanne, Susanna Lemmelä, Timo P. Sipilä, Tuomas Sipilä, Ulrike Lyhs, Vincent Llorens, Teemu Niiranen, Kati Kristiansson, Lotta Männikkö, Manuel González Jiménez, Markus Perola, Regis Wong, Terhi Kilpi, Tero Hiekkalinna, Elina Järvensivu, Essi Kaiharju, Hannele Mattsson, Markku Laukkanen, Päivi Laiho, Sini Lähteenmäki, Tuuli Sistonen, Sirpa Soini, Adam Ziemann, Anne Lehtonen, Apinya Lertratanakul, Bob Georgantas, Bridget Riley-Gillis, Danjuma Quarless, Fedik Rahimov, Graham Heap, Howard Jacob, Jeffrey Waring, Justin Wade Davis, Nizar Smaoui, Relja Popovic, Sahar Esmaeeli, Jeff Waring, Athena Matakidou, Ben Challis, David Close, Slavé Petrovski, Antti Karlsson, Johanna Schleutker, Kari Pulkki, Petri Virolainen, Lila Kallio, Arto Mannermaa, Sami Heikkinen, Veli-Matti Kosma, Chia-Yen Chen, Heiko Runz, Jimmy Liu, Paola Bronson, Sally John, Sanni Lahdenperä, Susan Eaton, Wei Zhou, Minna Hendolin, Outi Tuovila, Raimo Pakkanen, Joseph Maranville, Keith Usiskin, Marla Hochfeld, Robert Plenge, Robert Yang, Shameek Biswas, Steven Greenberg, Eija Laakkonen, Juha Kononen, Juha Paloneva, Urho Kujala, Teijo Kuopio, Jari Laukkanen, Eeva Kangasniemi, Kimmo Savinainen, Reijo Laaksonen, Mikko Arvas, Jarmo Ritari, Jukka Partanen, Kati Hyvärinen, Tiina Wahlfors, Andrew Peterson, Danny Oh, Diana Chang, Edmond Teng, Erich Strauss, Geoff Kerchner, Hao Chen, Hubert Chen, Jennifer Schutzman, John Michon, Julie Hunkapiller, Mark McCarthy, Natalie Bowers, Tim Lu, Tushar Bhangale, David Pulford, Dawn Waterworth, Diptee Kulkarni, Fanli Xu, Jo Betts, Jorge Esparza Gordillo, Joshua Hoffman, Kirsi Auro, Linda McCarthy, Soumitra Ghosh, Meg Ehm, Kimmo Pitkänen, Tomi Mäkelä, Anu Loukola, Heikki Joensuu, Juha Sinisalo, Kari Eklund, Lauri Aaltonen, Martti Färkkilä, Olli Carpen, Paula Kauppi, Pentti Tienari, Terhi Ollila, Tiinamaija Tuomi, Tuomo Meretoja, Anne Pitkäranta, Joni Turunen, Katariina Hannula-Jouppi, Sampsa Pikkarainen, Sanna Seitsonen, Miika Koskinen, Antti Palomäki, Juha Rinne, Kaj Metsärinne, Klaus Elenius, Laura Pirilä, Leena Koulu, Markku Voutilainen, Markus Juonala, Sirkku Peltonen, Vesa Aaltonen, Andrey Loboda, Anna Podgornaia, Aparna Chhibber, Audrey Chu, Caroline Fox, Dorothee Diogo, Emily Holzinger, John Eicher, Padhraig Gormley, Vinay Mehta, Xulong Wang, Johannes Kettunen, Katri Pylkäs, Marita Kalaoja, Minna Karjalainen, Reetta Hinttala, Riitta Kaarteenaho, Seppo Vainio, Tuomo Mantere, Seppo Vainio, Anne Remes, Johanna Huhtakangas, Juhani Junttila, Kaisa Tasanen, Laura Huilaja, Marja Luodonpää, Nina Hautala, Peeter Karihtala, Saila Kauppila, Terttu Harju, Timo Blomster, Hilkka Soininen, Ilkka Harvima, Jussi Pihlajamäki, Kai Kaarniranta, Margit Pelkonen, Markku Laakso, Mikko Hiltunen, Mikko Kiviniemi, Oili Kaipiainen-Seppänen, Päivi Auvinen, Reetta Kälviäinen, Valtteri Julkunen, Anders Malarstig, Åsa Hedman, Catherine Marshall, Christopher Whelan, Heli Lehtonen, Jaakko Parkkinen, Kari Linden, Kirsi Kalpala, Melissa Miller, Nan Bing, Stefan McDonough, Xing Chen, Xinli Hu, Ying Wu, Annika Auranen, Airi Jussila, Hannele Uusitalo-Järvinen, Hannu Kankaanranta, Hannu Uusitalo, Jukka Peltola, Mika Kähönen, Pia Isomäki, Tarja Laitinen, Teea Salmi, Anthony Muslin, Clarence Wang, Clement Chatelain, Ethan Xu, Franck Auge, Kathy Call, Kathy Klinger, Marika Crohns, Matthias Gossel, Kimmo Palin, Manuel Rivas, Harri Siirtola & Javier Gracia Tabuenca.

**GIGA study Group members**

Neema Kanyaro, Cyprian Ntomoka, Julius J. Massaga, Joyce K. Ikungura.
